# Circadian Disruption Drives Extracellular Matrix Remodeling to Facilitate Pulmonary Metastatic Colonization

**DOI:** 10.64898/2026.04.20.719725

**Authors:** Ignacio Aiello, Guido Hokama, Alessandro Ceci, Camila Senna, Diego A. Golombek, Natalia Paladino, Carla V. Finkielstein

## Abstract

Circadian clocks impose temporal architecture on signaling networks, and disruption of this architecture predisposes to cancer metastasis. Through direct pharmacological and genetic perturbation of core clock components, we establish that circadian desynchronization in mouse lung fibroblasts eliminates temporal migration gating, creating constitutive motility responses to TNF-α and TGF-β, and identify YAP/TEAD as the obligate, non-redundant convergence node through which clock-regulated ECM mechanical signals and cytokine-driven transcriptional programs jointly drive cellular motility. Chronic jet lag (CJL) in mice generates nocturnal TNF-α elevation (ZT12-21) that drives sustained matrix metalloprotease expression while simultaneously reorganizing Hippo and TGF-β signaling toward temporal convergence during daytime hours, enabling YAP/TEAD-dependent transcription to synergize with TGF-β signaling and drive epithelial-to-mesenchymal transition (EMT) programs that normally remain temporally restricted. Functional validation demonstrates CJL doubles metastatic colonization incidence (40% to 90%) following B16F10 melanoma inoculation. Critically, established metastases amplify these molecular changes: metastatic burden under CJL creates maximal TGF-β expression (ZT15-21), constitutive YAP activity, and sustained EMT marker expression, while eliminating M1/M2 macrophage temporal organization. Analysis of TCGA-SKCM metastatic melanoma datasets confirms that clock-disrupted human tumors exhibit selective strengthening of YAP/TAZ, EMT, and inflammatory pathway coupling, establishing that this convergence architecture is conserved in human disease. Together, these findings demonstrate that circadian disruption transforms from a facilitator of initial metastatic colonization into a driver of progressive metastatic burden by eliminating the temporal segregation that normally constrains pro-metastatic programs to discrete, non-overlapping windows, creating self-perpetuating cycles wherein pathway convergence facilitates colonization and established tumors amplify pro-metastatic signaling to maintain permissive microenvironmental conditions.

## INTRODUCTION

Circadian timekeeping systems orchestrate temporal coherence across virtually all physiological and behavioral processes, from sleep-wake cycles and body temperature regulation to endocrine secretion and immune function. This temporal organization enables organisms to anticipate and adapt to predictable environmental transitions inherent to the Earth’s 24-h rotation. At the molecular level, circadian oscillations emerge from interlocked transcriptional-translational feedback loops wherein positive regulators CLOCK and BMAL1 drive expression of negative regulators PER1-3 and CRY1-2, which in turn suppress CLOCK:BMAL1 activity in a cycle requiring approximately 24 h ^1^. The suprachiasmatic nuclei (SCN) of the anterior hypothalamus function as the master pacemaker, receiving photic information directly from melanopsin-expressing retinal ganglion cells and subsequently broadcasting temporal cues to peripheral oscillators throughout the brain and body via neurohumoral signals ^2^. Synchronization among this hierarchical network of cellular clocks maintains internal temporal order, but disruption of these relationships, whether through genetic perturbation, environmental desynchrony, or pathological processes, generates a state of temporal misalignment that predisposes to cardiometabolic disorders, obesity, and cancer ^2^.

The lung represents a particularly intriguing model for investigating circadian regulation of tissue homeostasis. Clock gene expression in pulmonary tissue exhibits robust circadian rhythmicity, with entrainment by multiple zeitgebers including feeding schedules, glucocorticoid hormones, and temperature cycles ^3–5^. Characterized as a “weak oscillator” displaying periodicities ranging from 20-28 hours ^6^, the lung’s temporal architecture reflects its remarkable cellular heterogeneity. This organ comprises type I and type II pneumocytes, Clara cells, fibroblasts, vascular endothelium, airway smooth muscle cells, and diverse immune populations, predominantly alveolar macrophages and dendritic cells. Structural integrity derives from the extracellular matrix (ECM), which not only provides physical scaffolding but actively regulates cell growth, morphology, and migration through dynamic reciprocal interactions with cellular components. Emerging evidence demonstrates that circadian clock dysfunction profoundly influences pulmonary function across diverse pathologies, including chronic obstructive pulmonary disease (COPD), idiopathic pulmonary fibrosis, and asthma ^3,7–11^. Notably, aberrant ECM composition and architecture associate strongly with chronic inflammation, creating a self-reinforcing cycle of tissue damage and impaired function ^12–14^.

Chronic inflammatory states frequently trigger maladaptive tissue repair processes that generate pathological healing responses and alter the mesenchymal cell populations essential for tissue regeneration ^15^. Central to this pathobiology is epithelial-to-mesenchymal transition (EMT), a reversible cellular reprogramming program wherein polarized epithelial cells lose their apicobasal architecture and acquire migratory and invasive capabilities ^16,17^. While essential for embryonic development and physiological wound healing, EMT contributes to pathological fibrosis and cancer metastasis ^18^. Rather than representing a binary switch, EMT progression involves intermediate hybrid states characterized by co-expression of epithelial and mesenchymal markers ^19,20^. This transition responds to diverse signaling factors that induce master transcriptional regulators including SNAIL (snail family zinc finger 1), TWIST (twist family bHLH transcription factor 1), and ZEB1 (Zinc finger E-box Binding homeobox 1), which coordinate the morphological and functional conversion ^21^. Multiple signaling cascades including Hippo, TGF-β (Transforming Growth Factor-β), EGF (Epidermal Growth Factor), and Wnt pathways govern this cellular plasticity ^22^.

The Hippo pathway represents a highly conserved signaling network controlling organ size, cell proliferation, and tissue regeneration. Pathway activity integrates diverse intrinsic signals (energy status, cellular stress, polarity) and extrinsic cues (mechanical forces from ECM, cell-cell contacts, matrix composition) ^23,24^. In mammals, pathway activation proceeds through a kinase cascade wherein the MST1/2 (Mammalian STerile 20-like kinase 1 and 2) heterodimer phosphorylates MOB1 (MOB kinase activator 1A) and LATS1/2 kinases (LArge Tumor Suppressor 1 and 2). Activated LATS1/2 directly phosphorylates YAP (Yes-Associated Protein) and its paralog TAZ (Transcriptional Coactivator with PDZ-binding Motif) at multiple sites, promoting 14-3-3 binding and cytoplasmic sequestration ^25,26^. Additional phosphorylation by CK1 (Casein Kinase 1) triggers β-TrCP (beta-Transducin repeat Containing Protein)-mediated ubiquitination and proteasomal degradation ^27^. Despite lacking intrinsic DNA-binding domains, YAP/TAZ function as powerful transcriptional co-activators through interaction with TEAD family (Transcriptional Enhanced Associate Domain) transcription factors, driving expression of genes promoting proliferation, survival, and EMT ^28,29^. ECM stiffness emerges as a critical regulator of Hippo signaling: compliant matrices activate the pathway and suppress YAP/TAZ, while rigid ECM inhibits Hippo components, allowing nuclear YAP/TAZ accumulation ^30,31^. This mechanotransduction link positions Hippo signaling as a key integrator of physical microenvironmental properties with cellular fate decisions.

Parallel to Hippo signaling, the TGF-β pathway exerts profound influence over EMT, fibrosis, and cancer progression. TGF-β ligand binding to type I and II serine/threonine kinase receptors triggers SMAD2/3 phosphorylation and complex formation with SMAD4, enabling nuclear translocation and transcriptional activation of target genes including those encoding EMT transcription factors and ECM components ^32,33^. Inhibitory SMAD7 provides negative feedback by competing with SMAD2/3 for receptor binding and recruiting ubiquitin ligases to promote receptor degradation ^34^. The balance between activating (SMAD2/3/4) and inhibitory (SMAD7) SMADs determines pathway output, with this equilibrium subject to extensive regulation. In the metastatic context, TGF-β exhibits paradoxical roles: tumor-suppressive in early stages through growth inhibition, but pro-metastatic in advanced disease through promotion of EMT, immune evasion, and ECM remodeling ^35,36^.

Cross-talk between Hippo and TGF-β pathways creates complex regulatory networks governing EMT and metastasis. YAP/TAZ can both amplify and be amplified by TGF-β signaling: YAP/TAZ enhance TGF-β pathway output through transcriptional regulation of pathway components, while TGF-β signaling stabilizes YAP/TAZ protein and promotes nuclear localization ^37^. This feed-forward architecture enables robust induction of EMT programs and persistent mesenchymal states. Matrix metalloproteases (MMPs) represent critical downstream effectors of both pathways, catalyzing ECM degradation and remodeling that facilitates cellular invasion. MMP2 and MMP9 (gelatinases) target type IV collagen and gelatin, while MMP3 (stromelysin-1) exhibits broader substrate specificity ^38^. Beyond direct proteolysis, MMPs regulate inflammatory signaling, growth factor availability, and immune cell recruitment through liberation of sequestered mediators and generation of bioactive fragments ^39^.

Metastasis, the dissemination and growth of tumor cells at distant sites, represents the primary cause of cancer mortality. This complex process requires cancer cells to successfully navigate multiple barriers: detachment from primary tumors, intravasation into circulation, survival in transit, extravasation into target organs, and ultimately colonization with outgrowth into macroscopic lesions ^40^. The lung constitutes a predominant metastatic site for diverse malignancies, reflecting both its extensive vasculature serving as a mechanical trap for circulating tumor cells and its unique microenvironmental properties that can either support or suppress metastatic colonization ^41^. The balance between tumor-promoting and tumor-suppressing factors within the pulmonary microenvironment, including immune populations, ECM composition, and inflammatory mediators, critically determines metastatic success.

Despite accumulating evidence linking circadian disruption to cancer incidence and progression ^42–48^, the mechanisms through which temporal misalignment specifically facilitates metastatic colonization remain incompletely understood. We hypothesized that circadian disruption creates permissive microenvironmental conditions in target organs by dysregulating ECM homeostasis, inflammatory signaling, and mechanotransduction pathways. Here, we demonstrate that chronic circadian desynchronization enhances pulmonary metastatic colonization in concert with coordinated upregulation of matrix metalloproteases, temporal reorganization of TGF-β and Hippo signaling pathways, and sustained activation of EMT programs. These findings are consistent with a model in which circadian clocks serve as temporal gatekeepers of tissue-level metastatic permissiveness, with broad implications for understanding how modern circadian disruption may accelerate cancer progression.

## RESULTS

### Loss of temporal organization eliminates circadian gates restricting fibroblast migration

Circadian clocks coordinate cellular migration through rhythmic regulation of cytoskeletal dynamics and adhesion machinery, establishing temporal gates that restrict migratory competence to defined circadian windows. This temporal gating is evident in multiple physiological contexts. For example, skin wounds inflicted during active phase heal faster than rest-phase injuries, reflecting time-of-day variation in fibroblast invasion capacity ^49^. This temporal control extends beyond mechanical responses to encompass signaling pathway regulation, as BMAL1 serves as an essential mediator of TGF-β-driven fibroblast differentiation in lung tissue ^50^. These observations position the circadian system as a critical determinant of fibroblast migratory competence in the lung microenvironment. To interrogate how loss of circadian synchronization influences this capacity in the context of metastatic colonization, we employed mouse lung fibroblast-like (MLg) cells maintained under synchronized or unsynchronized conditions, assessing migratory responses to growth factors (EGF, FGF2) and cytokines (TNF-α, TGF-β) at defined circadian times (Figure 1A).

**Figure 1.**
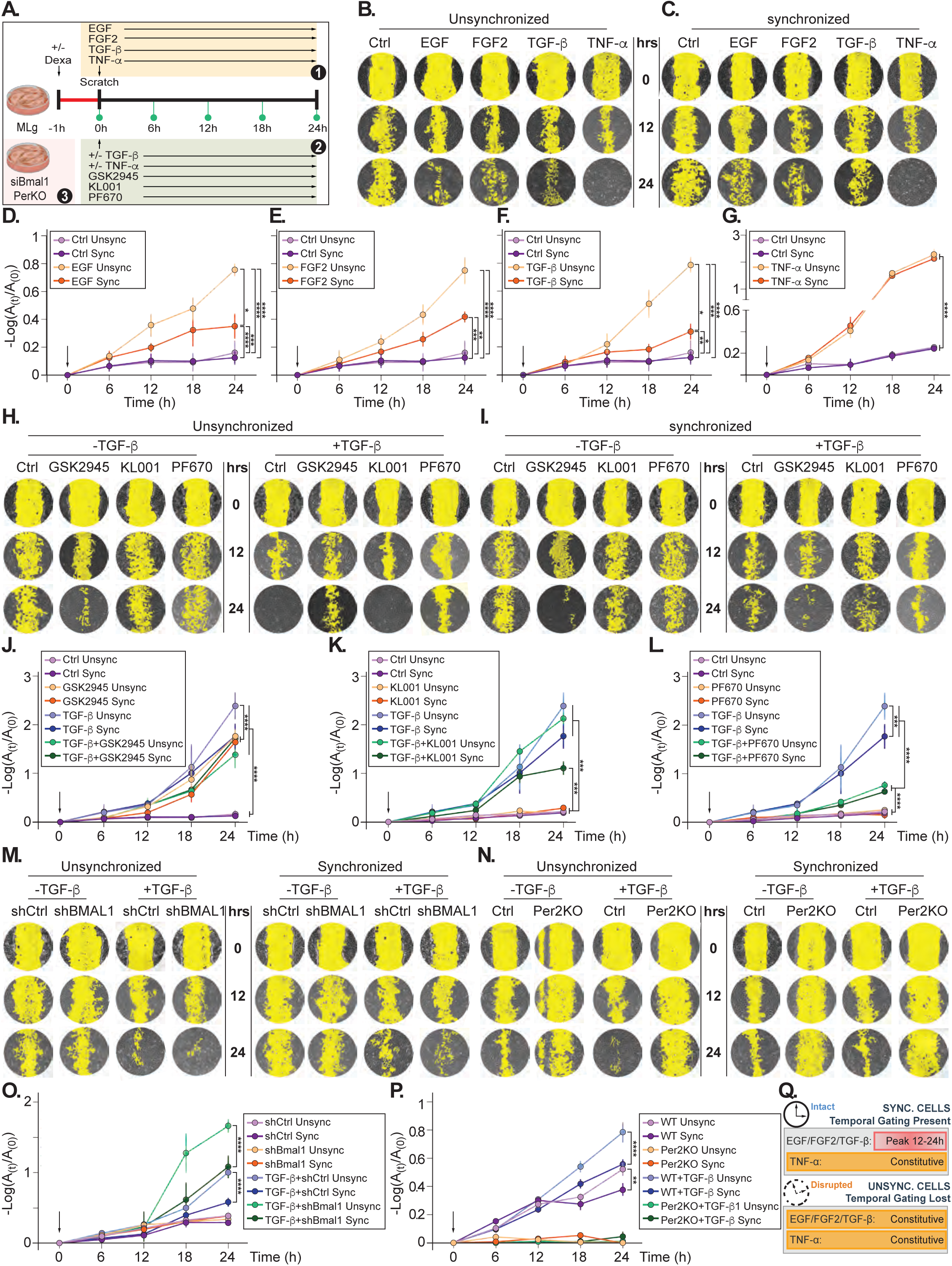
Lack of synchrony increased migration in fibroblast-like lung cells. Schematic representation of the experimental design used to test MLg cells migration under different treatments (**A**). Scenario 1 represents (**B-G**), scenario 2 represents (**H-L**) and scenario 3 represents (**M-P**). Representative images (**B-C**) and quantification (**D-G**) of MLg cell migration in response to growth factors (EGF, FGF2) or cytokines (TGF-β, TNF-α) under synchronized (sync) or unsynchronized (unsync) conditions. (**D-G**) Time-course quantification for EGF (**D**), FGF2 (**E**), TGF-β (**F**), and TNF-α (**G**) treatments, showing differential responses in synchronized (orange) versus unsynchronized (yellow) conditions. Control (Ctrl) indicates cells were either synchronized or unsynchronized, and no stimulus was applied. **Pharmacological and genetic perturbation of core clock components reveals differential regulation of basal and TGF-β-stimulated migration.** Representative images (**H-I**) and quantification (**J-L**) of MLg cell migration following treatment with clock-modulating compounds GSK2945 (REV-ERB antagonist), KL001 (CRY stabilizer), or PF670462 (CK1δ/ε inhibitor) in the absence or presence of TGF-β under unsynchronized (**H**) or synchronized (**I**) conditions. (**J-L**) Time-course analysis for GSK2945 (**J**), KL001 (**K**), and PF670462 (**L**) showing effects on basal and TGF-β-induced migration. Representative images (**M-N**) and quantification (**O-P**) of migration in Bmal1-depleted MLg cells (shBmal1, **M**, **O**) or Per2-knockout fibroblasts (Per2 KO) or wild-type (control, **N-P**) stimulated with TGF-β under synchronized or unsynchronized conditions. Temporal gating in migration is lost in unsynchronized cells that exhibit constitutive migration responses across all timepoints, which is further exacerbated in response to stimuli such as EGF, FGF2, TGF-b, and TNF-a. (**Q**) Migration is expressed as -Log(A₁/A₀), where A₁ represents the wound area at indicated timepoints and A₀ represents the initial wound area. (**B-C**, **H-I**, **M-N**) Cell-free wound areas are pseudocolored in yellow for visualization. (**D-G**, **J-L**, **O-P**). Data represent mean ± SD from n=2 independent experiments, with three scratches per well and four images per scratch at each timepoint. Two-way ANOVA: p<0.0001 for treatment factor (**D-G**, **J-L**, **O-P**). (**J-L**) Post-hoc comparisons in Supplementary Table 1.

Desynchronization eliminated temporal gating of migratory responses. Under unsynchronized conditions, all tested factors, EGF, FGF2, TGF-β, and TNF-α, significantly potentiated migration compared to vehicle controls, with responses appearing largely equivalent in magnitude across the 24-h observation period (Figures 1B, 1D-1G, yellow vs. light purple lines). Importantly, following circadian synchronization, MLg cells exhibited pronounced temporal restriction of migratory responses to EGF, FGF2, and TGF-β, with peak migration confined to the 12-24 h window post-synchronization (Figures 1C, 1D-1F, orange vs. purple lines). The temporal gating manifested as sustained attenuation throughout the observation period despite equivalent stimulus exposure (Figures 1D-1F, yellow vs. orange lines). TNF-α defied this pattern, eliciting comparable migratory responses regardless of synchronization state (Figure. 1G, yellow vs. orange lines), suggesting distinct regulatory mechanisms.

Constitutively elevated responses in desynchronized cells reveal that loss of temporal organization eliminates endogenous gating mechanisms. This temporal restriction normally reflects coordinated oscillations in cytoskeletal regulators and signaling components (Figure 1Q). Importantly, the ablation of this temporal architecture creates sustained migratory permissiveness that may facilitate metastatic colonization by eliminating temporal barriers to cell motility.

### Pharmacological and genetic perturbation of core clock components differentially regulates basal and stimulus-induced migration

External signals initiate migration, yet circadian clocks fundamentally shape when and how robustly cells respond. Across diverse contexts, dendritic cell trafficking, leukocyte homing, and neutrophil recruitment, circadian clocks temporally gate migratory responses rather than permitting constitutive responsiveness ^51–54^. To determine whether individual clock components exhibit differential control over basal migratory machinery versus cytokine-responsive pathways, we employed pharmacological and genetic perturbations targeting distinct oscillator nodes.

We treated MLg cells with three clock-modulating compounds: GSK2945, a selective REV-ERB antagonist that maintains BMAL1 transcription levels independently of RORα activity [a mechanism distinct from RORα silencing, which itself promotes migration ^55,56^]; KL001, a CRY stabilizer that lengthens circadian period by inhibiting FBXL3-dependent CRY degradation ^57^; and PF670462, a CK1δ/ε inhibitor that modulates PER phosphorylation and alters circadian period ^58^.

Remarkably, these clock perturbations revealed divergent mechanisms of migration regulation. Under unsynchronized conditions, GSK2945 treatment alone significantly enhanced basal migration even without TGF-β stimulation, with this effect persisting throughout the timepoints analyzed (Figures 1H, 1J, yellow versus light purple lines; Supplementary Table 1). This constitutive elevation of motility suggests that REV-ERB antagonism, and steady BMAL1 levels, directly influences the basal migratory machinery independent of external stimuli. In marked contrast, KL001 and PF670462 showed minimal effects on unstimulated migration (Figures 1H, 1K-1L, yellow versus light purple lines; Supplementary Table 1) but selectively potentiated TGF-β-induced migration, amplifying the cytokine response without altering basal rates (Figures 1H, 1K-1L; Supplementary Table 1). Following synchronization, these patterns persisted: GSK2945 maintained its capacity to promote both basal and TGF-β-stimulated migration, while KL001 and PF670462 again amplified TGF-β responses without affecting unstimulated motility (Figures 1I-1L; Supplementary Table 1).

To complement these pharmacological approaches, we examined genetic perturbations of core clock components. BMAL1 knockdown (sh*Bmal1*; Figure S1) dramatically enhanced TGF-β-induced migration even in dexamethasone-synchronized cells, with sh*Bmal1* cells displaying robust migratory responses that exceeded control levels across the observation period (Figures 1M-1O; Supplementary Table 2). Notably, BMAL1 depletion abolishes core clock function due to its non-redundant role as the essential positive transcriptional activator in the molecular feedback loop rendering cells constitutively arrhythmic ^59,60^. In marked contrast, Per2-knockout fibroblasts completely abrogated TGF-β-induced migration (Figures 1N, 1P, dark and light green lines), whereas sh*Bmal1* cells showed robust enhancement (Figure 1O, dark and light green lines; Supplementary Table 2).

Collectively, these findings establish that core clock components exhibit functionally distinct roles in regulating cellular migration. Clock perturbations bifurcate into two classes: those that modulate basal migratory capacity independent of external stimuli (GSK2945/REV-ERB antagonism, potentially reflecting direct BMAL1 effects on cytoskeletal machinery), and those that primarily amplify responsiveness to pro-migratory cues without affecting unstimulated motility (KL001/CRY stabilization, PF670462/CK1δ/ε inhibition). The genetic experiments further reveal that loss of positive clock elements (BMAL1) versus negative regulators (PER2) produces overlapping yet mechanistically distinct effects on migration, with complete clock ablation (sh*Bmal1*) generating constitutive enhancement while partial disruption (PER2KO with intact PER1/3 compensation) produces more nuanced alterations. These observations position the molecular clock not as a monolithic regulator but as a modular system wherein distinct components govern separate aspects of migratory behavior, basal motility machinery versus stimulus-response coupling, with both contributing to temporal gating of cell movement.

### Circadian desynchronization transforms temporally restricted metalloprotease induction into sustained proteolytic competence

Matrix metalloproteases (MMPs) represent critical effectors of cellular migration and ECM remodeling, catalyzing proteolytic degradation that facilitates invasion. Given that circadian synchronization temporally gates migratory responses to TGF-β (Figure 1F), and that this gating depends on core clock component function (Figures 1H-1P), we hypothesized that MMP expression would similarly exhibit circadian regulation of both basal levels and stimulus-induced responses. We quantified *Bmal1*, *Mmp2*, *Mmp3*, and *Mmp9* mRNA levels across 24 h in synchronized versus unsynchronized MLg cells treated with vehicle, TGF-β, or GSK2945 (Figure 2).

**Figure 2.**
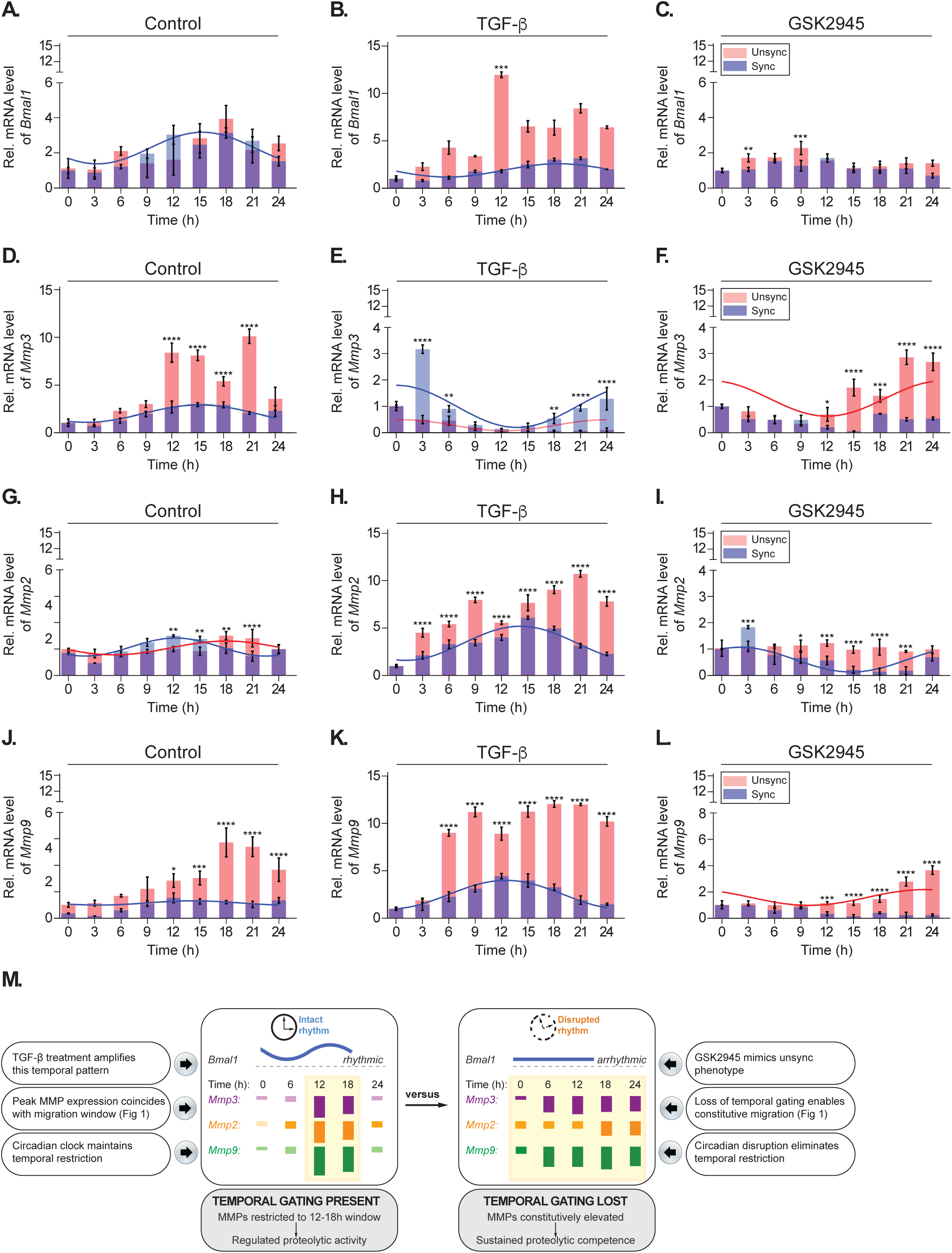
Circadian synchronization governs metalloprotease expression magnitude and temporal dynamics. Relative mRNA levels of *Bmal1* (**A-C**), *Mmp3* (**D-F**), *Mmp2* (**G-I**), and *Mmp9* (**J-L**) quantified by RT-qPCR across 24 h in synchronized (blue) versus unsynchronized (orange) MLg cells treated with vehicle control (**A**, **D**, **G**, **J**), TGF-β (**B**, **E**, **H**, **K**), or GSK2945 (**C**, **F**, **I**, **L**). Purple bars indicate overlapping synchronized and unsynchronized values. Curves represent statistically significant circadian oscillations detected by MetaCycle analysis (p<0.05; Supplementary Table 3). Data represent mean ± SD from n=3 biological replicates per timepoint. (**M**) Schematic summary illustrating temporal MMP gating in synchronized versus unsynchronized MLg cells. Left: Synchronized cells exhibit rhythmic *Bmal1* expression and temporally restricted MMP expression (12-18h window). Right: Unsynchronized cells show arrhythmic *Bmal1* and constitutive MMP elevation at all timepoints, eliminating temporal gating.

Under control conditions, *Bmal1* exhibited robust circadian oscillation with peak expression during 12-18 h post-synchronization, validating successful entrainment (Figure 2A; Supplementary Table 3). Remarkably, TGF-β treatment preserved this rhythmic pattern in synchronized cells while simultaneously inducing a pronounced elevation (∼2-fold peak at 12 h) in unsynchronized cells that lacked oscillatory behavior (Figure 2B; Supplementary Table 3). This differential response establishes that circadian state determines whether TGF-β signaling maintains or disrupts core clock gene rhythmicity. As expected, GSK2945 treatment abolished *Bmal1* rhythmicity and significantly maintains basal expression levels in both synchronized and unsynchronized conditions (Figure 2C; Supplementary Table 3).

MMP expression revealed circadian state-dependent regulation with gene-specific patterns. *Mmp3* displayed rhythmic expression peaking at 12-18 h under control conditions in synchronized cells, with significantly elevated constitutive levels in unsynchronized cells (Figure 2D; Supplementary Table 3). Remarkably, TGF-β treatment produced diametrically opposite effects contingent on synchronization status: in synchronized cells, TGF-β further reduced *Mmp3* expression while shifting the peak to 24-3 h, whereas in unsynchronized cells, TGF-β markedly suppressed *Mmp3* levels across the entire time course (Figure 2E; Supplementary Table 3). This inversion of TGF-β responsiveness represents an example of how circadian status can fundamentally reverse signaling outcomes. GSK2945 treatment abolished synchronization-dependent patterns, maintaining *Mmp3* at constitutive low levels resembling the unsynchronized state regardless of actual synchronization status (Figure 2F; Supplementary Table 3), corroborating that REV-ERB antagonism disrupts temporal gating by maintaining basal BMAL1 levels and eliminating rhythmic regulation.

*Mmp2* displayed more subtle circadian regulation, showing modest rhythmicity under basal conditions. TGF-β enhanced *Mmp2* expression primarily during 12-18 h in synchronized and across all timepoints in unsynchronized cells, though unsynchronized cells exhibited markedly higher absolute levels (Figures 2G-2H; Supplementary Table 3). This relative insensitivity to synchronization status under control conditions suggests that *Mmp2* regulation may be governed by circadian-independent mechanisms or subject to distinct regulatory nodes compared to *Mmp3* and *Mmp9*. Notably, analysis of additional stimuli revealed that EGF and FGF2 suppressed *Mmp2* while enhancing *Mmp9* at 12 and 21 h, whereas TNF-α dramatically elevated *Mmp2*, *Mmp3*, and *Mmp9* at both timepoints (Figures S2B-S2D), indicating stimulus-specific regulation of MMP expression patterns.

*Mmp9* exhibited the most dramatic circadian amplification. Under control conditions, *Mmp9* showed modest rhythmicity with gradual elevation during late timepoints in unsynchronized cells (Figure 2J; Supplementary Table 3). TGF-β treatment produced profound circadian-dependent effects: in synchronized cells, TGF-β generated robust circadian amplification with expression peaking sharply at 9-15 h post-stimulation, coinciding with the temporal window of peak migration observed in Figure 1F (Figure 2K; Supplementary Table 3). In marked contrast, unsynchronized cells exhibited sustained elevation of *Mmp9* throughout the observation period without temporal restriction, creating a state of constitutive proteolytic competence (Figure 2K). GSK2945 again normalized these patterns toward the unsynchronized phenotype (Figure 2L).

These temporal profiling results establish several critical principles. First, circadian synchronization governs not merely the magnitude but the temporal architecture of *MMP* induction. Synchronized cells confine peak *MMP* expression to discrete circadian windows, while desynchronized cells exhibit temporal deregulation producing sustained elevation (Figure 2M). Second, circadian state can invert TGF-β responsiveness, as exemplified by *Mmp3*, suggesting that the molecular clock actively determines signaling polarity rather than passively modulating amplitude. Third, different MMPs show distinct degrees of circadian coupling, *Mmp9* exhibits profound temporal gating, *Mmp3* shows moderate regulation with signal inversion, while *Mmp2* displays relative independence, positioning these proteases as differentially regulated effectors within the ECM remodeling machinery. Finally, GSK2945-mediated clock disruption recapitulates the unsynchronized phenotype, establishing that loss of temporal organization through REV-ERB antagonism produces constitutive *MMP* expression analogous to the sustained migratory permissiveness observed in Figure 1J. These findings extend the temporal gating framework established in Figure 1 to the level of ECM remodeling machinery, wherein desynchronization converts episodic MMP induction into constitutive proteolytic output (Figure 2M).

### Chronic circadian disruption generates nocturnal elevation of inflammatory and proteolytic programs in lung tissue

Having established that circadian desynchronization eliminates temporal gating of migratory responses (Figure 1Q), disrupts core clock component regulation (Figures 1H-1P), and produces constitutive metalloprotease expression in cultured MLg fibroblasts (Figure 2), we interrogated whether these cellular phenotypes manifest in pulmonary tissue subjected to chronic circadian disruption *in vivo*. We employed a well-established chronic jet lag (CJL) paradigm wherein mice experienced repeated 6h phase advances every 48 h for 6 weeks, compared to control animals maintained under standard LD 12:12 cycles ^61^. Lung tissue was harvested at 3h intervals during a complete circadian cycle to capture temporal dynamics of inflammatory mediators and matrix remodeling enzymes.

Initial validation confirmed circadian disruption at the molecular level: core clock genes *Bmal1*, *Cry1*, and *Cry2* exhibited robust oscillations in LD lung tissue but showed markedly altered or abolished rhythmicity under CJL (Figures S3S-S3C; Supplementary Table 4), establishing effective clock disruption in the pulmonary compartment. Against this backdrop of compromised circadian architecture, we examined inflammatory and proteolytic gene expression.

*Ccl2* (encoding MCP-1, a key monocyte chemoattractant) showed circadian rhythmicity under CJL condition but with differing phase relationships and elevated expression compared to LD at specific zeitgeber times, particularly at ZT9-15 (Figure 3A; Supplementary Table 5). Importantly, *Tnf-α* expression exhibited a profound transformation under chronic disruption: whereas LD control lungs showed minimal *Tnf-α* expression with no discernible circadian pattern, CJL lungs displayed dramatic nocturnal elevation on expression (ZT12-21) with nearly 4-fold amplification relative to daytime nadirs (Figure 3B; Supplementary Table 5). This emergence of robust *Tnf-α* enhancement under circadian disruption, rather than simple amplitude enhancement, suggests that chronic temporal misalignment actively induces inflammatory signaling programs during the night phase. In contrast, *Tgf-β* expression remained relatively stable across both zeitgeber time and lighting conditions (Figure 3C; Supplementary Table 5), indicating that not all pro-fibrotic pathways respond uniformly to circadian disruption.

**Figure 3.**
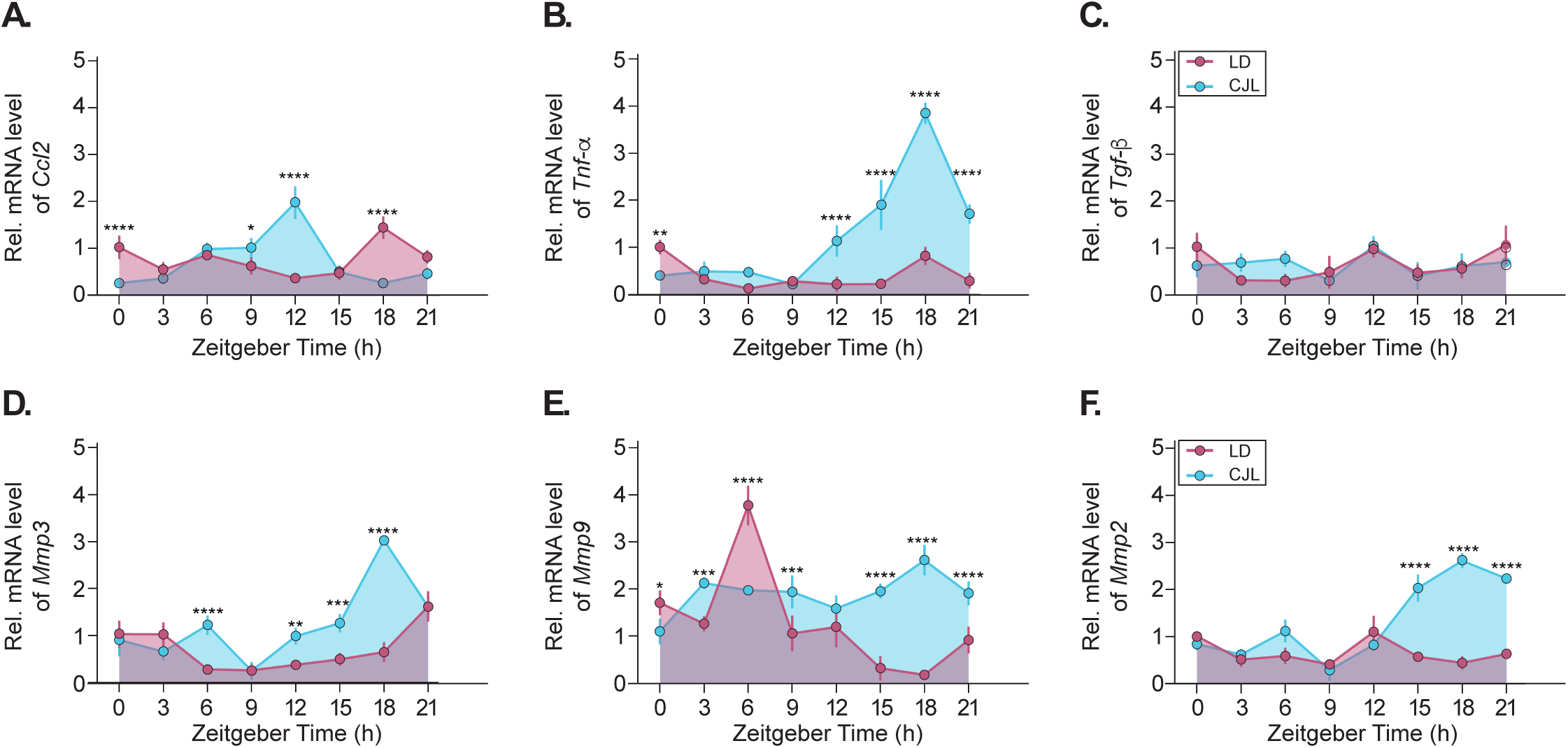
Chronic circadian disruption elevates nocturnal expression of inflammatory mediators and matrix metalloproteases in lung tissue. Relative mRNA levels of *Ccl2* (**A**), *Tnf-α* (**B**), *Tgf-β* (**C**), *Mmp3* (**D**), *Mmp9* (**E**), and *Mmp2* (**F**) quantified by RT-qPCR in lung tissue from mice maintained under standard LD 12:12 (pink) or subjected to chronic jet lag (CJL, cyan) for 6 weeks. Tissues were harvested at 3h intervals across a complete 24h cycle. Note the emergence of robust *Tnf-α* rhythmicity with dramatic nocturnal peak exclusively under CJL (**B**), and enhanced nocturnal expression of *Mmp3*, *Mmp9*, and *Mmp2* under circadian disruption (**D-F**). Purple shading indicates overlapping LD and CJL values. Data represent mean ± SD from n=4 animals per timepoint. Two-way ANOVA: (**A**, **B**, **D-F**) p<0.0001 for lighting condition; (**A-F**) p<0.0001 for zeitgeber time. Post-hoc comparisons: *p<0.05, **p<0.01, ***p<0.001, ****p<0.0001.

Examining matrix metalloproteases revealed patterns concordant with our *in vitro* observations and mechanistically linked to TNF-α dynamics. *Mmp3* displayed modest rhythmicity under LD control but showed elevated nocturnal expression under CJL, particularly at ZT12-21 (Figure 3D; Supplementary Table 5). *Mmp9* exhibited daily variation under LD conditions with a daytime peak at ZT6 (Figure 3E). In contrast, CJL lungs displayed a sustained elevation throughout the circadian cycle, particularly during the night (ZT12-21, Figure 3E; Supplementary Table 5). *Mmp2* showed the clearest CJL-specific pattern, with pronounced nocturnal elevation at ZT15-21 that was largely absent under LD conditions (Figure 3F; Supplementary Table 5). Critically, the temporal window of peak MMP expression (ZT12-21) directly overlaps with the dramatic nocturnal TNF-α surge (ZT12-21), suggesting a causal relationship. This inference is strongly supported by our *in vitro* studies demonstrating that TNF-α represents the most potent inducer of all three MMPs in fibroblasts, elevating *Mmp2*, *Mmp3*, and particularly *Mmp9* expression at both 12h and 21h post-treatment (Figures S2B-S2D), a temporal pattern recapitulated *in vivo*. The concordance between cellular TNF-α responsiveness and tissue-level temporal coincidence establishes TNF-α as a likely driver of nocturnal MMP elevation under CJL.

These *in vivo* findings validate our cellular model while suggesting a mechanistic basis for these associations. Chronic circadian disruption correlates with nocturnal upregulation of inflammatory mediators (*Tnf-α*, *Ccl2*) and matrix metalloproteases (*Mmp2*, *Mmp3*, *Mmp9*). The temporal convergence is mechanistically coherent: elevated TNF-α signaling, which promoted constitutive migration *in vitro* (Figure 1G) and potently induced MMP expression in cultured fibroblasts (Figure S2), peaks at ZT12-21 under CJL, directly preceding and overlapping with maximal MMP expression (ZT12-21). This temporal sequence suggests that the nocturnal TNF-α surge likely drives MMP upregulation, creating windows wherein the lung microenvironment exhibits both heightened inflammatory tone and elevated proteolytic capacity. Given that nocturnal mice normally exhibit enhanced physiological repair processes during their rest phase (day), the inversion of this temporal architecture represents a fundamental disruption of tissue homeostatic rhythms ^62–66^. This multi-scale concordance between cellular and tissue phenotypes suggests that cellular circadian disruption aligns with organismal circadian misalignment.

### YAP/TEAD activity represents an obligate mediator of cytokine-induced migration independent of circadian state

Having established sustained metalloprotease expression under circadian desynchronization (Figures 2-3), we examined mechanotransduction pathways linking ECM remodeling to migration. MMP-mediated matrix degradation alters tissue stiffness, which regulates Hippo pathway activity and YAP/TAZ nuclear localization. We hypothesized that YAP/TEAD transcriptional complexes integrate ECM dysregulation with migratory competence. To test this, we treated MLg cells with verteporfin, a YAP/TEAD inhibitor ^67^. To probe temporal requirements, verteporfin was added at three different timepoints: simultaneously with cytokine (0h), during the early migratory phase (6h), or at mid-observation (12h), with experiments performed under both synchronized and unsynchronized conditions (Figure 4A).

**Figure 4.**
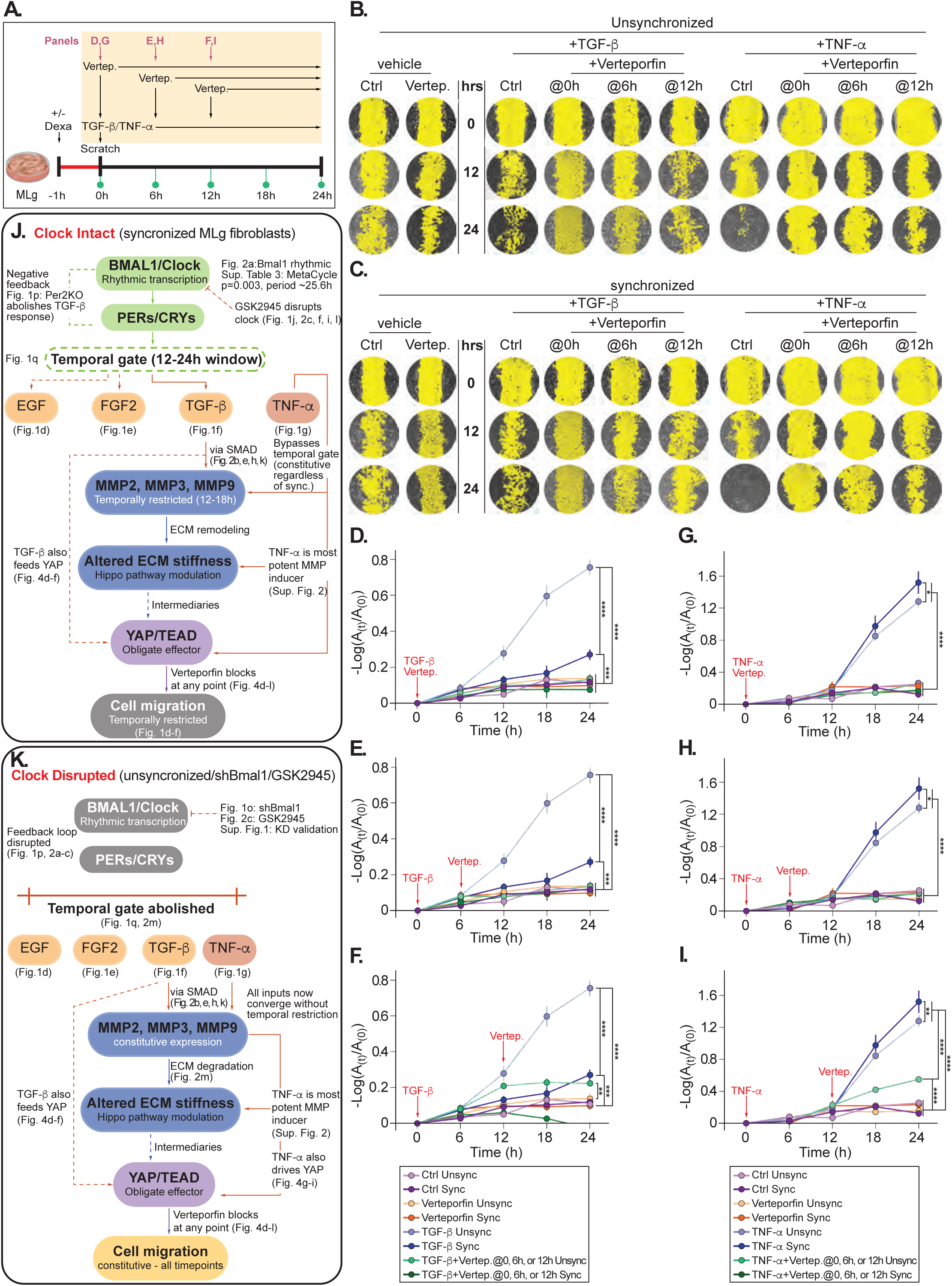
YAP/TEAD activity is obligate for cytokine-induced migration regardless of circadian state or inhibitor timing. (**A**) Schematic representation of experimental design. MLg cells were synchronized with dexamethasone (Dexa, +/-) at - 1h, scratched at 0h, and treated with TGF-β or TNF-α. Verteporfin (YAP/TEAD inhibitor) was added at 0h, 6h, or 12h post-scratch (indicated by arrows), with wound closure monitored at 6h intervals for 24h. Representative images (**B-C**) and quantification (**D-I**) of MLg cell migration in response to TGF-β (**D-F**) or TNF-α (**G-I**) with or without verteporfin (YAP/TEAD complex inhibitor) under synchronized (**B**) or unsynchronized (**C**). Verteporfin was added at 0h (simultaneous with cytokine, **D**, **G**), 6h post-cytokine (**E**, **H**), or 12h post-cytokine (**F**, **I**), indicated by red arrows. Note that both TGF-β (light blue and dark blue lines) and TNF-α (light blue and dark blue lines) robustly induce migration in synchronized and unsynchronized cells, but verteporfin treatment (orange and green lines) completely abolishes migration regardless of addition timing or synchronization status. (**B-C**) Cell-free wound areas are pseudocolored in yellow for visualization. Migration is expressed as -Log(A₁/A₀). Data represent mean ± SD from n=2 independent experiments, with three scratches per well and four images per scratch at each timepoint. Two-way ANOVA: (**D-I**) p<0.0001 for treatment and timepoint factors. Post-hoc comparisons in Supplementary Table 6: *p<0.05, **p<0.01, ***p<0.001, ****p<0.0001. (j–k) Mechanistic schematics summarizing the circadian clock-dependent regulation of cytokine-induced migration and YAP/TEAD signaling identified in Figs. 1–4. (**J**) In synchronized MLg fibroblasts with an intact molecular oscillator, BMAL1/CLOCK-driven rhythmic transcription, validated by Bmal1 circadian oscillation (Figure 2A; MetaCycle p = 0.003, period ∼25.6h; Supplementary Table 3) and PER2-dependent feedback (Figure 1P), imposes a temporal gate (12–24h window) on migratory responses to EGF, FGF2, TGF-β, and TNF-α (Figures 1D–1G). Within this gated window, *Mmp2*, *Mmp3*, and *Mmp9* expression is temporally restricted to the 12–18h phase (Figure 2), driving ECM remodeling that modulates tissue stiffness and suppresses Hippo pathway activity, ultimately enabling nuclear YAP/TEAD complex formation. YAP/TEAD functions as an obligate downstream effector of both TGF-β (via SMAD; Figures 2B, 2E, 2H, 2K) and TNF-α signaling (the most potent MMP inducer; Figure S2), with verteporfin-mediated disruption of YAP/TEAD complexes abolishing migration at any point of intervention (Figures 4D-4L). (**K**) Upon clock disruption, through desynchronization, shBmal1-mediated BMAL1 depletion (Figure 1O and S1), or GSK2945 treatment (Figures 1J, 2C, 2F, 2I, 2L), the feedback loop between BMAL1/CLOCK and PER/CRY components is dismantled, eliminating the temporal gate (Figures 1Q, 2M). All cytokine inputs (EGF, FGF2, TGF-β, TNF-α) now converge without temporal restriction, driving constitutive *Mmp2*, *Mmp3*, and *Mmp9* expression (Figure 2M) and sustained ECM degradation that maintains continuous Hippo pathway suppression and nuclear YAP accumulation. The result is constitutive YAP/TEAD-dependent cell migration across all timepoints, with verteporfin retaining its capacity to abolish migration regardless of intervention timing (Figures 4D–4L), establishing YAP/TEAD as the obligate convergence node integrating clock-dependent and clock-independent pro-migratory signals.

Our results revealed YAP/TEAD as an absolute, non-redundant requirement for cytokine-induced migration across all conditions tested. TGF-β robustly induced migration in both unsynchronized and synchronized cells across the 24h observation period, recapitulating the differential responses observed in Figure 1 (Figures 4D-4F, light and dark blue lines). However, verteporfin co-treatment completely abolished this response regardless of addition timing. When added simultaneously with TGF-β (0h), migration was prevented entirely (Figures 4B-4D, orange and green lines remaining flat). Importantly, even when verteporfin was added after TGF-β at 6h or 12h, timepoints when cytokine-induced migration was actively progressing, the migratory response was immediately arrested and cells remained essentially immotile for the remainder of the observation period (Figures 4B-4C, 4E-4F, lines flattening upon verteporfin addition indicated by red arrows). This pattern was identical in both synchronized and unsynchronized cells, demonstrating that YAP/TEAD requirement is independent of circadian state. TNF-α treatment revealed an identical YAP/TEAD requirement. Consistent with Figure 1G, TNF-α promoted robust migration independent of the synchronization status of the cells (Figures 4G-4L, light blue and dark blue lines). The treatment with verteporfin completely abolished this response regardless of addition timing, whether applied simultaneously with TNF-α (0h) or during active migration (6h, 12h; red arrows), reducing migratory responses to baseline levels indistinguishable from unstimulated controls in both synchronized and unsynchronized conditions (Figures 4B-4C, 4G-4L, orange and green lines; Supplementary Table 6).

These findings establish several critical principles. First, YAP/TEAD activity represents an <u>obligate mediator</u> of both TGF-β and TNF-α signaling in the context of fibroblast migration: there are no compensatory pathways capable of driving motility when YAP/TEAD is disrupted. Second, this requirement is both <u>temporally invariant</u> and <u>immediately essential</u>: YAP/TEAD inhibition blocks migration regardless of circadian phase, synchronization status, or when the inhibitor is introduced, with cells halting migration immediately upon verteporfin treatment even when cytokine stimulation preceded inhibitor addition by hours. Third, the uniformity of verteporfin’s effects across both cytokines suggests <u>a common</u> <u>mechanotransduction node</u>, despite TGF-β showing circadian-gated responses while TNF-α does not (Figure 1), both signals converge on YAP/TEAD as an obligate effector.

The mechanistic integration of these findings is summarized schematically in Figures 4J-4K. Under intact circadian conditions (Figure 4J), BMAL1/CLOCK-driven rhythmic transcription maintains temporal gating through PER/CRY negative feedback, restricting EGF-, FGF2-, and TGF-β-induced MMP expression and downstream migration to discrete 12-24h windows, while TNF-α bypasses this gate entirely. Regardless of temporal restriction, all cytokine-driven migration converges on YAP/TEAD as the obligate downstream effector via ECM remodeling and Hippo pathway modulation. Under disrupted circadian conditions (Figure 4K), loss of BMAL1 function (shBmal1, GSK2945, or desynchronization) abolishes the temporal gate, producing constitutive MMP expression and sustained ECM degradation. All cytokine inputs now converge simultaneously on YAP/TEAD without temporal restriction, creating constitutive migratory competence at all timepoints. This model predicts that circadian disruption should alter YAP signaling dynamics in vivo, a hypothesis we examined in subsequent experiments.

### Chronic circadian disruption reorganizes Hippo pathway temporal dynamics in lung tissue

Figures 1–4 establish a causal mechanistic framework through direct pharmacological and genetic perturbation: circadian desynchronization eliminates temporal gating of fibroblast migration, produces constitutive metalloprotease expression, and positions YAP/TEAD as an obligate, non-redundant effector of cytokine-induced motility. Having established this framework in vitro, we next asked whether the same signaling architecture is temporally reorganized in lung tissue under chronic circadian disruption *in vivo*, examining YAP expression and phosphorylation status alongside upstream and downstream regulatory components *Mob1a* and *Tead4* throughout complete circadian cycles in LD versus CJL mice (Figure 5).

**Figure 5.**
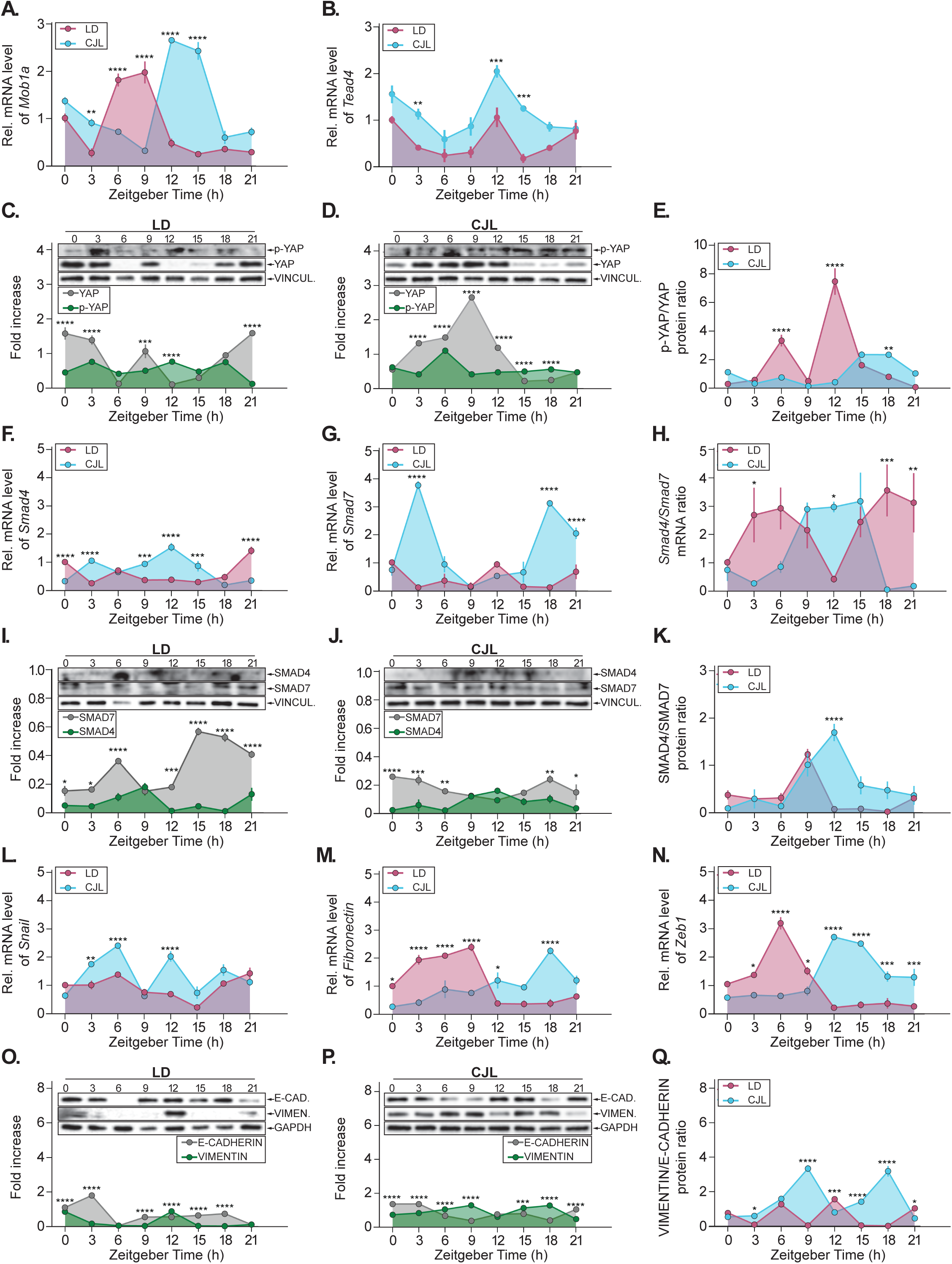
Chronic circadian disruption reorganizes temporal dynamics of Hippo and TGF-β pathways and induces mesenchymal marker expression in lung tissue. Relative mRNA levels of *Mob1a* (**A**) and *Tead4* (**B**) quantified by RT-qPCR in lung tissue from mice maintained under LD 12:12 (pink) or chronic jet lag (CJL, cyan) at various zeitgeber times. Purple shading indicates overlapping values. Representative Western blots (top) and densitometric quantification (bottom) of phospho-YAP (Ser127, inactive cytoplasmic form) and total YAP proteins in lung tissue from LD (**C**) or CJL (**D**), and p-YAP/YAP ratio (**E**) in mice harvested at 3h intervals. Quantification shows fold-increase relative to VINCULIN for total YAP (gray) and phospho-YAP (green). Note that under LD, phospho-YAP peaks at ZT12, indicating maximal Hippo activity and YAP inactivation during the day, while CJL alters this temporal relationship with elevated phospho-YAP during night (ZT15-18). Relative mRNA levels of *Smad4* (**F**) and *Smad7* (**G**) quantified by RT-qPCR, and SMAD4/SMAD7 mRNA ratio (**H**) in lung tissue from mice maintained under LD 12:12 (pink) or chronic jet lag (CJL, cyan) at and collected at different zeitgeber times. Purple shading indicates overlapping values. Representative Western blots (top) and densitometric quantification (bottom) of SMAD4 and SMAD7 proteins in lung tissue from LD (**I**) or CJL (**J**) mice harvested at 3h intervals. Vinculin serves as loading control. (**K**) SMAD4/SMAD7 protein ratio comparing LD (pink) and CJL (cyan) conditions. Note that CJL produces elevated SMAD4/SMAD7 ratios during the day (ZT9-12), indicating enhanced TGF-β signaling capacity during this temporal window. Relative mRNA levels of EMT transcription factors *Snail* (**L**), *Fibronectin* (**M**), and *Zeb1* (**N**) quantified by RT-qPCR in lung tissue from mice maintained under LD 12:12 (pink) or chronic jet lag (CJL, cyan) at zeitgeber times. Purple shading indicates overlapping values. Representative Western blots (top) and densitometric quantification (bottom) of E-CADHERIN (epithelial marker, **O-P**, green line) and VIMENTIN (mesenchymal marker, **O-P** gray line) proteins in lung tissue from LD (**O**) or CJL (**P**) and VIMENTIN/E-CADHERIN ratio (**Q**) in mice harvested at 3-hour intervals. GAPDH serves as loading control. Note that CJL produces dramatic elevation of *Zeb1* during day (ZT9-18) and sustained VIMENTIN expression, while reducing E-CADHERIN amplitude. Data represent mean ± SEM from n=4 animals per timepoint. Two-way ANOVA: (**A-B**, **G**, **L**, **O-Q**) p<0.0001, (**F**) p=0.011, (**H**) p=0.0028 (**M**) p=0.001 for lighting condition; (**C-D, I-J**) p<0.0001 for protein type and (**A-D**, **F-I**, **K-Q**) p<0.0001, (**J**) p=0.0449 for zeitgeber time. Post-hoc comparisons: *p<0.05, **p<0.01, ***p<0.001, ****p<0.0001.

We first examined transcriptional regulation of Hippo pathway components. *Mob1a* and *Tead4* exhibited robust daily variation with altered phase relationships under CJL. *Mob1a* peaked at ZT6-9 under LD but shifted to early night (ZT12-15) under CJL (Figure 5A, pink vs. cyan lines), while *Tead4* showed elevated expression with higher amplitude under CJL compared to LD at all times (Figure 5B, cyan vs. pink lines), demonstrating that circadian disruption remodels temporal availability of Hippo signaling components.

Protein-level analysis of YAP revealed temporal regulation of both total expression and phosphorylation status. Under LD conditions, phospho-YAP (Ser127), the inactive, cytoplasmic form generated by LATS1/2 kinase activity that is subsequently targeted for proteasomal degradation, exhibited a peak at ZT12 (Figure 5C, green line). Total YAP protein showed a daily variation with elevation during the late night and early day transition (ZT18-ZT3) (Figure 5C, gray line). The antiphasic relationship between phospho-YAP (day peak) and total YAP (night peak) generates a temporal window during the night/day transition (ZT18-ZT3) permitting maximal YAP accumulation and transcriptional output under LD conditions (Figure 5E, pink line).

Chronic jet lag fundamentally inverted these temporal dynamics. Under CJL, phospho-YAP exhibited a peak at ZT6 and sustained levels during the night (ZT15-21, Figure 5D, green line), shifting to the LD pattern where phospho-YAP peaks during the end of the day, while total YAP showed elevated levels predominantly during day (ZT3-12, Figure 5D, gray line). This reorganization inverts the temporal window of YAP transcriptional activity: whereas LD permits YAP-mediated transcription during the end of the night (ZT18-3) when phospho-YAP is low, CJL shifts the permissive window to the day (ZT3-12) when total YAP is elevated and phospho-YAP levels are relatively lower (Figure 5E, cyan line), fundamentally reversing the temporal architecture of mechanotransduction signaling. Mechanistically, this reorganization integrates with the TNF-α-driven MMP cascade (Figures 3-4 and S2). This is, nocturnal TNF-α elevation (ZT12-21) induces sustained proteolytic activity that remodels ECM stiffness, suppressing Hippo signaling and promoting the daytime YAP permissive window, thereby enabling constitutive rather than temporally restricted mechanotransduction.

### Chronic circadian disruption reorganizes TGF-β pathway temporal dynamics and signaling capacity

Having established that circadian disruption reorganizes Hippo pathway dynamics (Figures 5A-5E), we examined TGF-β signaling components given the extensive cross-talk between Hippo/YAP-TAZ and TGF-β/SMAD signaling in controlling EMT programs and migration ^22,68,69^.

We first examined *Smad4* and *Smad7* mRNA expression throughout circadian times. *Smad4* exhibited daily variations under both LD and CJL, though with altered peaks of expression (Figure 5F). Under LD, *Smad4* showed modest rhythmicity with slightly elevated expression during the late night and early day transition (ZT18-ZT3; Figure 5F, pink line), while CJL produced more pronounced oscillation with peaks at ZT3 and ZT9-ZT15 (Figure 5F, cyan line). *Smad7* displayed more dramatic regulation with distinct patterns of variations between lighting conditions (Figure 5G). LD lungs showed relatively stable *Smad7* levels with modest elevation at ZT12 (Figure 5G, pink line), whereas CJL lungs exhibited pronounced peaks at ZT3 and ZT18 with significantly elevated expression compared to LD (ZT15-ZT21; Figure 5G, cyan line).

The *Smad4*/*Smad7* mRNA ratio revealed temporal reorganization of TGF-β signaling capacity. Under LD, the ratio showed an elevation during late night (ZT18-ZT6), indicating a temporal window of enhanced pathway responsiveness during this phase (Figure 5H, pink line). CJL fundamentally altered this pattern, producing elevated ratios during day to early night (ZT9-15) rather than late night to early day in LD (Figure 5H, cyan line), suggesting a shift in the temporal window of maximal TGF-β signaling capacity from night to day under chronic circadian disruption.

Protein-level analysis confirmed these transcriptional patterns while revealing additional complexity. Western blot analysis showed that SMAD4 and SMAD7 proteins both exhibited temporal variation in lung tissue (Figures 5I-5J). Under LD conditions, SMAD4 protein displayed pronounced changes in expression with a peak at ZT9 (Figure 5I, green line), while SMAD7 showed a peak at ZT6 and sustained high expression levels during night, ZT12-ZT0 (Figure 5I, gray line). The resulting SMAD4/SMAD7 protein ratio peaked at ZT9 under LD (Figure 5K, pink line). CJL conditions altered these dynamics: SMAD4 protein showed reduced amplitude of expression throughout the time course analyzed with a modest peak at ZT12 (Figure 5J, green line), while SMAD7 maintained relatively stable levels (Figure 5J, gray line). Critically, the SMAD4/SMAD7 protein ratio under CJL exhibited a pronounced peak at ZT12 (Figure 5K, cyan line), confirming the mRNA-level observation that circadian disruption shifts maximal TGF-β signaling capacity to daytime hours.

These findings establish that chronic circadian disruption reorganizes TGF-β pathway temporal architecture, shifting peak SMAD4/SMAD7 ratios from ZT9 (LD) to ZT12 (CJL). This reorganization creates temporal convergence with the daytime YAP permissive window established in Figure 5E, wherein the feed-forward architecture of YAP/TAZ and TGF-β mutual amplification enables synergistic co-activation that would remain segregated under normal circadian conditions. Combined with nocturnal TNF-α elevation (ZT12-21, Figure 3B) and sustained MMP expression (Figures 2-3), this pathway convergence would eliminate the temporal segregation that normally constrains pro-metastatic signaling to discrete windows.

### Chronic circadian disruption activates EMT programs in lung tissue

The temporal convergence of YAP transcriptional activity and TGF-β signaling capacity during day under CJL (Figure 5A-5K), combined with nocturnal TNF-α elevation (Figure 3), predicts downstream activation of epithelial-to-mesenchymal transition (EMT) programs that enable migratory and invasive phenotypes essential for metastatic colonization. Given that both YAP/TEAD and TGF-β/SMAD pathways directly activate EMT transcription factors (SNAIL, ZEB1, TWIST), we examined whether circadian disruption influences EMT marker expression in lung tissue.

Analysis of EMT transcription factors revealed circadian regulation with CJL-dependent alterations. *Snail* exhibited time-of-day-dependent expression under both conditions, with CJL producing elevated levels at ZT3-6 and a peak at ZT12 compared to LD (Figure 5L, cyan line). *Fibronectin*, encoding a mesenchymal ECM component, showed temporal variations under LD with daytime accumulation (ZT0-ZT9; Figure 5M, pink line), while CJL conditions generated pronounced elevation during night (ZT12-21) with significantly higher expression than LD (Figure 5M, cyan line). Most dramatically, *Zeb1* displayed a profound CJL-induced elevation during night (ZT12-ZT21; Figure 5N, cyan line). This nocturnal *Zeb1* peak under CJL temporally follows the daytime YAP/TGF-β pathway convergence (ZT3-12, Figures 5A-5K) and directly overlaps with the TNF-α surge (ZT12-ZT21, Figure 3B), suggesting that coordinated activation of these upstream pathways drives *Zeb1* transcription. The temporal sequence is mechanistically coherent: daytime YAP and TGF-β co-activation initiates EMT transcriptional programs, while the subsequent nocturnal TNF-α surge, which potently induces MMPs (Figure S2) and promoted constitutive migration (Figure 1G), sustains and amplifies *Zeb1* expression, positioning this EMT transcription factor as a molecular integrator of temporally converging pro-metastatic signals.

Protein-level analysis of canonical EMT markers revealed a fundamental reorganization of the epithelial-mesenchymal balance. Under LD conditions, E-CADHERIN (epithelial marker) exhibited a peak expression during early day (ZT3) followed by sustained levels from ZT9-ZT18 (Figure 5O, gray line), while VIMENTIN (mesenchymal marker) showed a discrete peak at ZT12 with basal levels maintained from ZT21 through early morning (ZT0) (Figure 5O, green line). This temporal organization restricts VIMENTIN expression to a narrow window at the day-night transition under normal circadian conditions (Figure 5Q, pink line). CJL fundamentally disrupted this temporal architecture: E-CADHERIN maintained sustained expression throughout the 24-h cycle (Figure 5P, gray line), while VIMENTIN exhibited dramatic and sustained elevation throughout day and night (Figure 5P, green line). The net effect is a shift toward constitutive mesenchymal marker expression, with VIMENTIN levels under CJL exceeding LD values at nearly all zeitgeber times and eliminating the temporal restriction that normally confines mesenchymal marker expression to discrete circadian phases (Figure 5Q, pink line for LD vs. cyan line for CJL).

These findings establish coordinated EMT activation at transcriptional and protein levels under circadian disruption. Importantly, *Zeb1* exhibits pronounced nocturnal elevation (ZT12-21) specifically under CJL, while VIMENTIN protein shows sustained elevation throughout the circadian cycle. This elimination of temporal restriction in mesenchymal marker expression represents the downstream consequence of the pathway convergence characterized in Figures 3-5; this is, coordinated YAP and TGF-β signaling drives constitutive rather than temporally gated EMT programs.

### Chronic circadian disruption enhances pulmonary metastatic colonization

The preceding results established that chronic circadian disruption creates a coordinated pro-metastatic lung microenvironment characterized by temporally converging signaling pathways. We hypothesized that these coordinated molecular changes would translate to enhanced metastatic colonization *in vivo*. To test this directly, we subjected mice to either standard LD 12:12 or CJL conditions for 3 weeks, then inoculated B16F10 melanoma cells intravenously at day 21 (ZT6, daytime) (Figure S4). Animals remained under their respective lighting conditions for an additional 21 days before the lung was harvested and the metastatic burden quantified (Figure 6A).

**Figure 6.**
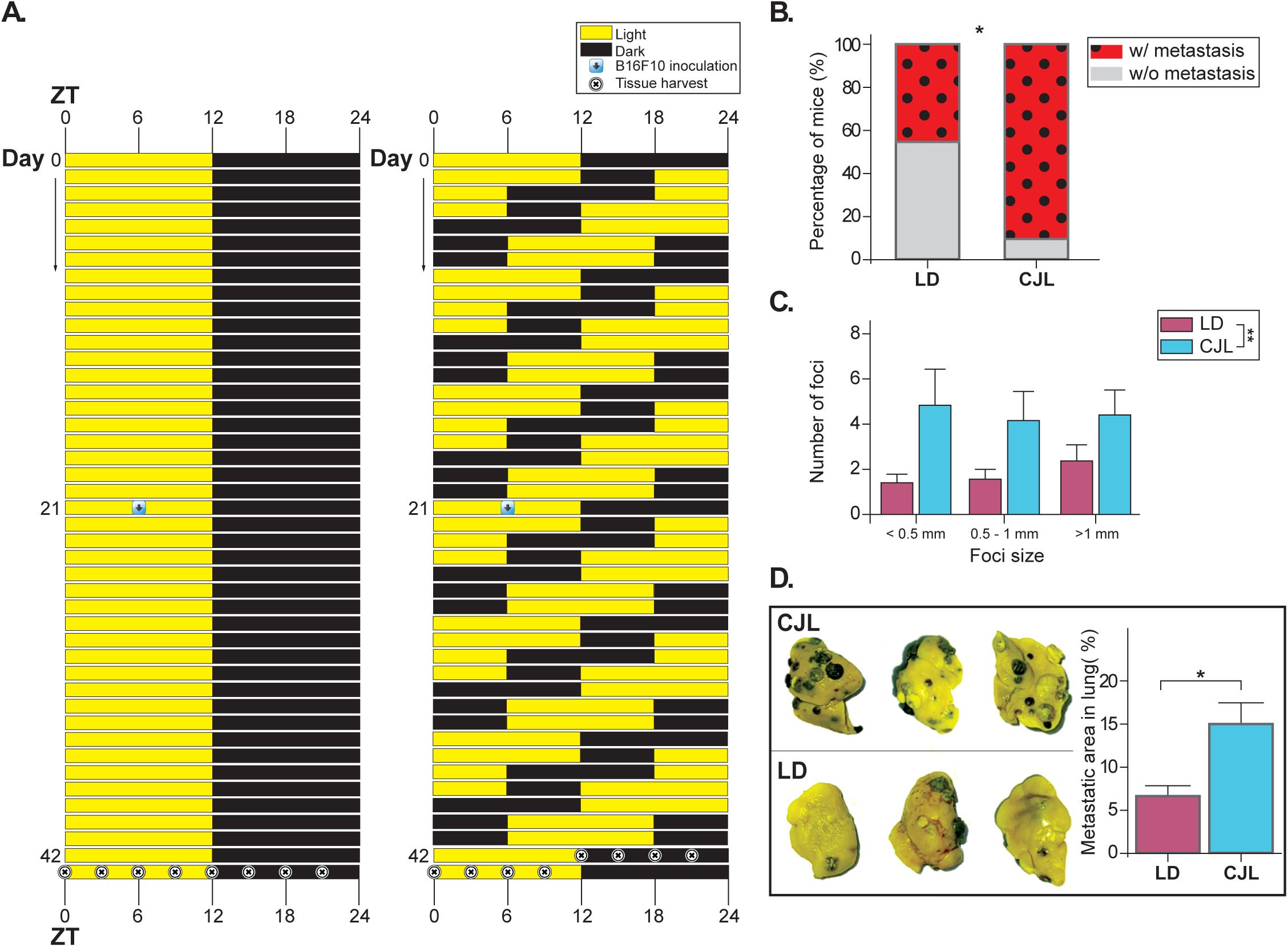
Chronic circadian disruption enhances pulmonary metastatic colonization. (**A**) Experimental design: Mice maintained under LD 12:12 or chronic jet lag (CJL) for 3 weeks received intravenous B16F10 melanoma cells at day 21 (ZT6), then remained under respective lighting conditions until lung harvest at day 42. Yellow = light phase, black = dark phase. Experimental timeline: Blue arrows indicate B16F10 melanoma cell inoculation (Day 21, ZT6), and crossed circles indicate lung tissue harvest (Day 42, various ZTs). (**B**) Percentage of mice developing detectable lung metastases under LD (∼40%) versus CJL (∼90%) (black dots in red background). Chi-square test: p=0.045. (**C**) Quantification of metastatic foci per lung grouped by size (<0.5 mm, 0.5-1 mm, >1mm diameter) showing increased large metastases under CJL. Two-way ANOVA: p=0.0093 per condition. Data represent mean ± SEM from n=9-10 animals per group. (**D**) Representative metastasis-bearing lungs from LD and CJL mice showing enhanced metastatic burden (dark-stained areas) under circadian disruption. Right: Quantification of the percentage of lung area occupied by metastatic lesions. Data represent mean ± SEM from n=9-10 animals per group. Unpaired t-test: *p=0.0309.

Our results show that CJL dramatically increased metastatic incidence and burden. While approximately 40% of LD control mice developed detectable lung metastases, nearly 90% of CJL-exposed mice showed metastatic lesions (Figure 6B, p=0.045), representing more than a doubling of metastatic incidence. Among metastasis-bearing animals, CJL significantly increased the number and size of metastatic foci (Figure 6C, p=0.0093). This corresponded to a significant increase in the percentage of total metastatic lung area (Figure 6D), indicating that circadian disruption facilitates both initial colonization and subsequent metastatic outgrowth. Representative lung images demonstrate the dramatic difference in metastatic burden, with CJL lungs exhibiting numerous large surface metastases (dark-stained areas) compared to the sparse lesions in LD controls (Figure 6D). These data validate that the molecular changes induced by chronic circadian disruption translate to enhanced metastatic colonization.

### Chronic circadian disruption reorganizes temporal dynamics of pulmonary macrophage populations

Metastatic success depends not only on stromal and matrix properties deregulated under CJL (Figures 3, 5), but also on immune cell populations. Macrophages exhibit functional plasticity from M1 (anti-tumor) to M2 (pro-tumor) phenotypes, with M2 macrophages promoting metastatic colonization through immunosuppression, angiogenesis, and ECM remodeling ^70^. Given that circadian clocks regulate immune cell trafficking and function, we hypothesized that macrophage temporal organization represents an additional component of the pro-metastatic microenvironment generated by circadian disruption.

Flow cytometric analysis of control (non-tumor-bearing) lungs revealed basal temporal patterns under LD conditions. Although total macrophage numbers did not differ throughout the day (Figure S5), M1 and M2 subpopulations exhibited time-of-day variations: M1 macrophages showed sustained levels throughout all circadian times (Figure 7A, red line), while M2 macrophages showed a modest elevation during late night (ZT21, Figure 7A, blue line). These temporal fluctuations generated corresponding variations in the M1:M2 ratio, with anti-tumor predominance at ZT3 that diminished by ZT9-21 under LD conditions (Figure 7C, pink line). CJL abolished these temporal patterns in control lungs: M1 macrophages dropped dramatically at ZT21 (Figure 7B, red line) while M2 macrophages remained elevated (Figure 7B, blue line), producing a constitutively low M1:M2 ratio throughout the day (Figure 7C, cyan line). This indicates that circadian disruption eliminates temporal organization of immune surveillance even in non-pathological tissue.

**Figure 7.**
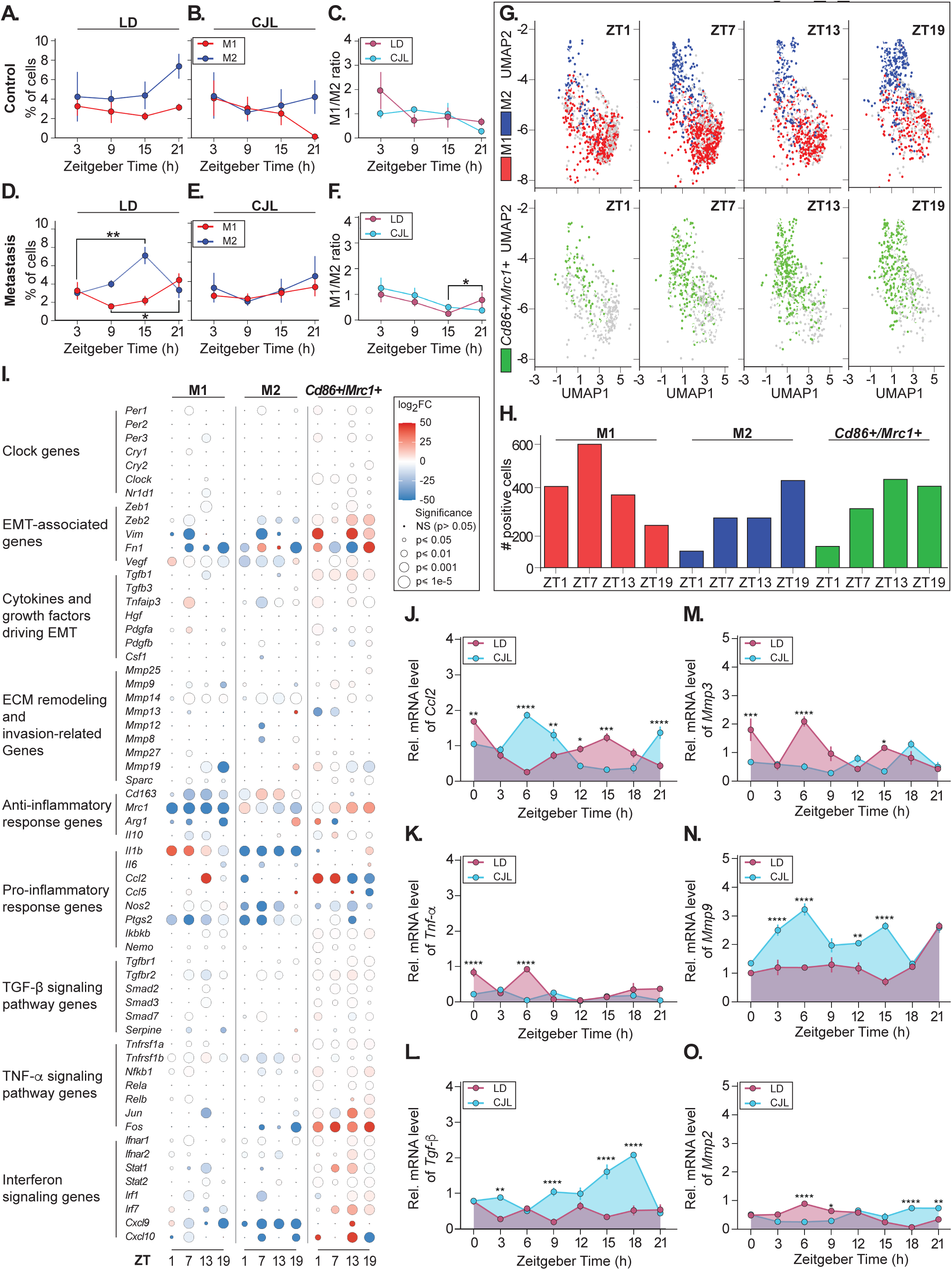
Chronic circadian disruption reorganizes macrophage dynamics and dysregulates inflammatory and proteolytic pathways in lung tissue. Flow cytometric quantification of M1 (CD86^+^, anti-tumor) and M2 (*Mrc1+*/CD206^+^, pro-tumor) macrophage populations in control (**A-C**) and metastasis-bearing (**D-F**) lungs from mice maintained under LD 12:12 or CJL. Samples were harvested at ZT3, 9, 15, and 21. (**A-B**) M1 (red) and M2 (blue) populations in control lungs under LD (**A**) or CJL (**B**). (**C**) M1/M2 ratio in control lungs under LD (pink) or CJL (cyan). (**D-E**) M1 and M2 populations in metastatic lungs under LD (**D**) or CJL (**E**). (**F**) M1/M2 ratio in metastatic lungs under LD (pink) or CJL (cyan). Note temporal organization under LD that is lost under CJL. Data represent mean ± SEM from n=4-6 animals per condition and timepoint. Two-way ANOVA: (**A**) p=0.0434, (**D**) p=0.0005 for lighting condition; (**D**) p=0.0071, (**F**) p=0.05 for zeitgeber time. Post-hoc: *p<0.05, **p<0.01. (**G**) UMAP plots from single-cell RNA-seq (GSE260641) showing macrophage populations at ZT01, 07, 13, and 19. Upper panels: M1 (*Cd86*⁺, red) and M2 (*Mrc1*⁺, blue) populations. Lower panels: Double-positive (*Cd86*⁺/*Mrc1*⁺, green) population. Background cells (gray/white) represent remaining macrophages and are preserved for spatial context. **(H)** Quantification of positive cells for each population across zeitgeber times. (**I**) Dot plot showing differential gene expression from single-cell RNA-seq dataset [GSE260641; ^71^] of tumor-associated macrophages isolated from B16F10 subcutaneous tumor-bearing mice maintained under standard LD conditions and harvested at ZT01, ZT07, ZT13, and ZT19. Macrophages were classified into three populations based on marker expression: *Cd86⁺* (M1), *Mrc1⁺* (M2), and *Cd86⁺*/*Mrc1⁺* double-positive macrophages. Genes are grouped by functional category. Dot size represents statistical significance (p-value); dot color indicates log₂ fold change (red = upregulated, blue = downregulated). Relative mRNA levels of *Ccl2* (**J**), *Tnf-α* (**K**), *Tgf-β* (**L**), *Mmp3* (**M**), *Mmp9* (**N**), and *Mmp2* (**O**) quantified by RT-qPCR in metastasis-bearing lung tissue from mice maintained under LD 12:12 (pink) or CJL (cyan) and harvested 21 days post-B16F10 inoculation at zeitgeber times. Purple shading indicates overlapping values. Note dramatic elevation of *Tgf-β* under CJL during night (ZT15-21) and sustained *Mmp9* elevation under CJL throughout the cycle. Data represent mean ± SEM from n=4 animals per timepoint. Two-way ANOVA: (**K-L**, **M-N**) p<0.0001 for lighting condition, (**J-O**) p<0.0001 for zeitgeber time. Post-hoc: *p<0.05, **p<0.01, ***p<0.001, ****p<0.0001.

Metastatic burden amplified and reorganized these dynamics. Under LD conditions, metastasis-bearing lungs showed pronounced temporal reorganization: M2 macrophages peaked at ZT15 (Figure 7D, blue line) while M1 macrophages were elevated at ZT21-3 (Figure 7D, red line), with statistically significant temporal differences between these populations (Figure 7D, **p<0.01, blue line and *p<0.05, red line). The M1:M2 ratio in metastatic LD lungs remained relatively stable, suggesting that while both populations expand in response to metastatic challenge, their temporal relationship is maintained (Figure 7F, *p<0.05, pink line). In marked contrast, CJL-exposed metastatic lungs exhibited loss of temporal structure: both M1 and M2 populations showed decreased and arrhythmic levels (Figure 7E, red and blue lines), and the M1:M2 ratio displayed variability without organized temporal patterning (Figure 7F, cyan line). This temporal disorganization under CJL suggests elimination of coordinated immune responses that would normally restrict metastatic growth during specific circadian phases.

To characterize macrophage heterogeneity and functional states at higher resolution, we analyzed publicly available single-cell RNA sequencing data from B16F10 subcutaneous tumor-bearing mice maintained under LD conditions and harvested at ZT01, 07, 13, and 19 [GSE260641; ^71^]. UMAP dimensional reduction revealed distinct M1 (*Cd86⁺*, red) and M2 (*Mrc1⁺*, blue) populations maintaining stable cell-type identity throughout the day (Figure 7G). Despite this stable linage commitment, both populations exhibited temporal variation in abundance (Figure 7H), recapitulating the antiphasic M1/M2 dynamics observed in our flow cytometry analysis (Figure 7D). A double-positive (*Cd86⁺*/*Mrc1⁺*, green) population was also evident (Figures 7G-7H), potentially representing an intermediate activation state or an M2-like subpopulation with pro-inflammatory signatures ^72^.

Differential gene expression analysis revealed functionally distinct transcriptional programs providing mechanistic context for immune dysregulation under CJL. M1 macrophages exhibited robust expression of pro-inflammatory markers including cytokines (*Il1b*), chemokines (*Ccl2*, *Cxcl9*), and interferon-stimulated genes (*Irf7*), consistent with anti-tumor surveillance functions (Figure 7I). *Tnfaip3*, a negative regulator of M1 polarization, showed upregulation at ZT07, supporting the presence of regulatory mechanisms that may limit classical M1 activation in the tumor microenvironment. Clock gene expression in M1 populations was modest yet significant, with expression of *Per1*, *Per3*, *Cry1*, and *Nr1d1*, but minimal *Per2* and *Cry2*, indicating these cells possess circadian regulatory capacity, though perhaps less robust than other tissue-resident cell types.

M2 macrophages displayed a transcriptional signature with direct mechanistic relevance to the pro-metastatic microenvironment characterized in Figs. 3 and 5 (Figure 7I). These cells exhibited strong expression of classical M2 markers (*Cd163*, *Arg1*) and elevated expression of specific matrix metalloproteases, particularly *Mmp13* and *Mmp19*, directly implicating these immune cells as contributors to the ECM remodeling programs identified in Figs. 2-3. The co-expression of MMPs and EMT-associated genes (*Vim*, *Fn1*, *Vegf*) in M2 populations indicates these cells contribute to both matrix remodeling and pro-angiogenic signaling within the metastatic niche. TGF-β signaling pathway gene expression was present but more restricted than initially anticipated.

The double-positive *Cd86⁺*/*Mrc1⁺* population displayed characteristics of both M1 and M2 phenotypes, representing a functionally hybrid state (Figure 7I). This population showed strong M2-like oriented features including high *Arg1* expression, anti-inflammatory cytokines (*Il10*, *Tgfb1*), and platelet-derived growth factor (*Pdgfa*), alongside transcriptional regulators (*Tnfaip3*, *Fos*, *Jun*). However, these cells preserved M1 characteristics through expression of chemokines (*Ccl2*, *Cxcl10*), interferon-stimulated genes (*Irf7*, *Irf1*, *Ifnar1*, *Ifnar2*), and TNF-α signaling components (*Nfkb1*, *Rela*). Most notably, the double-positive population exhibited enhanced clock gene expression (*Clock*, *Nr1d1*, *Per2*, *Per3*), elevated matrix metalloprotease activity (*Mmp9*, *Mmp19*), and strong EMT-associated gene patterns (*Vim*, *Fn1*, *Zeb2*), positioning these cells as highly activated hybrid states contributing to both inflammatory signaling and matrix remodeling.

Zeitgeber time-dependent variation in pro-inflammatory (*Il1b*, *Ccl2*), interferon pathway (*Irf1*, *Cxcl9*), and transcriptional regulation (*Fos*) genes was observed in all three populations under LD tumor-bearing conditions (Figure 7I), with the double-positive population displaying the most pronounced temporal oscillations. This demonstrates that macrophage functional states oscillate throughout the day even when absolute cell numbers remain relatively stable. The loss of this temporal regulation under CJL, as demonstrated by flow cytometric analysis (Figures 7B-7C, 7E-7F), would likely eliminate these time-of-day-dependent immune programs, creating sustained rather than temporally gated pro-tumorigenic conditions.

These findings suggest that circadian disruption reorganizes pulmonary immune surveillance through two complementary mechanisms. First, CJL eliminates temporal patterning of M1 and M2 populations in both control and metastatic lungs, creating constitutive rather than rhythmic immune activity. Second, the double-positive population contributes to pro-metastatic signaling through simultaneous expression of inflammatory mediators, matrix remodeling enzymes, and EMT-promoting factors, potentially amplifying the stromal pathway convergence characterized in Figures 3 and 5. The loss of temporally organized anti-tumor surveillance, combined with sustained activation of hybrid macrophage states, provides an additional immune-mediated mechanism through which circadian disruption facilitates the enhanced metastatic colonization observed in Figure 6.

### Metastatic burden and circadian disruption synergistically dysregulate inflammatory and proteolytic pathways

Having established that circadian disruption enhances metastatic colonization (Figure 6), at least in part through elimination of temporal immune organization (Figures 7A-7I), we examined whether established metastatic burden amplifies the inflammatory and matrix remodeling pathway dysregulation characterized in non-tumor-bearing CJL lungs (Figures 3 and 5).

Inflammatory mediators showed complex regulation by the interaction of metastatic burden and circadian state. *Ccl2* exhibited elevated expression under CJL with prominent peaks at ZT6 and ZT21 and sustained levels in the middle of the day (ZT9-ZT18; Figure 7J, cyan line), contrasting with the more focused ZT9-12 elevation observed in non-tumor-bearing CJL lungs (Figure 3A, cyan line). This temporal expansion suggests metastatic burden broadens the window of chemokine production. *Tnf-α* displayed patterns distinct from non-tumor-bearing conditions: whereas non-tumor LD lungs showed minimal *Tnf-α* expression (Figure 3B, pink line), metastatic LD lungs exhibited two expression peaks during early day (ZT0 and ZT6, Figure 7K, pink line). In contrast to the dramatic nocturnal elevation observed in non-tumor CJL lungs (ZT12-21, Figure 3B, cyan line), metastatic CJL lungs maintained sustained low-level expression throughout the circadian cycle (Figure 7K, cyan line). This indicates that metastatic burden fundamentally alters TNF-α regulation, potentially through feedback mechanisms initiated by established tumor-immune interactions.

Most significantly, *Tgf-β* exhibited dramatic and selective elevation in metastatic lungs under CJL, with expression being elevated from ZT9-ZT21 including a pronounced nocturnal peak at ZT15-ZT21 (Figure 7L, cyan line), a pattern absent in LD-exposed metastatic lungs (Figure 7L, pink line) and markedly different from the stable expression observed in non-tumor-bearing animals (Figure 3C, cyan line). This selective amplification of TGF-β under CJL conditions in the metastatic context directly validates the temporal reorganization of TGF-β signaling capacity characterized in Figure 5 and positions established metastases as active drivers of further TGF-β pathway activation. The temporal alignment of peak *Tgf-β* expression (ZT15-ZT21) with the period of M2 macrophage abundance under CJL (Figure 7E) suggests coordinated activation of pro-tumorigenic signaling networks.

Matrix metalloproteases displayed synergistic regulation by metastatic burden and circadian disruption. *Mmp3* showed reduced, yet constant, expression in metastatic compared to control lungs, particularly under CJL conditions (Figure 7M, cyan line), potentially reflecting tissue-specific responses to established tumor burden. In contrast, *Mmp9* expression exhibited a sustained high elevation under CJL compared to LD throughout all timepoints (ZT0-121, Figure 7N, cyan vs. pink line), representing amplification of the nocturnal *Mmp9* elevation observed in non-tumor-bearing CJL lungs (Figure 3E). This sustained rather than temporally restricted *Mmp9* expression in metastatic CJL lungs indicates that established tumors maintain the proteolytic activity initially induced by circadian disruption. *Mmp2* showed an enhanced nocturnal expression under CJL (ZT15-21, Figure 7O, cyan line), recapitulating and extending the pattern observed in control CJL lungs (Figure 3F) and temporally aligning with peak *Tgf-β* expression.

The temporal convergence of *Tgf-β* and *Mmp2* expression (ZT15-ZT21) with constitutive *Mmp9* elevation suggest the emergence of a self-reinforcing cycle. This is, circadian disruption enables initial colonization (Figure 6), established metastases amplify pro-metastatic signaling, and this amplification perpetuates permissive microenvironmental conditions. This mechanistic framework establishes circadian disruption as an active driver, not merely a facilitator, of progressive metastatic burden through continuous pathway amplification.

### Metastatic burden alters Hippo pathway temporal dynamics and YAP activity windows

To determine whether metastatic burden alters the Hippo pathway reorganization characterized in non-tumor-bearing lungs (Figures 5A-5E), we examined *Mob1a* and *Tead4* mRNA and YAP protein dynamics in metastasis-bearing lungs harvested 21 days post-B16F10 inoculation. This analysis reveals how established metastases modulate the mechanotransduction pathways obligate for migration (Figure 4).

*Mob1a* mRNA expression in metastatic lungs showed temporal regulation under both LD and CJL conditions, but with patterns distinct from non-tumor-bearing tissue (compare Figure 5A to Figure 8A). Under LD, metastatic lungs exhibited elevated *Mob1a* at ZT0-ZT6 (Figure 8A, pink line), contrasting with the ZT6-ZT9 peak observed in non-tumor-bearing LD lungs (Figure 5A, pink line). Under CJL, *Mob1a* showed sustained elevation at ZT12-21 (Figure 8A, cyan line), representing a temporal expansion compared to the ZT12-15 peak in non-tumor-bearing CJL lung (Figure 5A, cyan line). This suggests that metastatic burden broadens the temporal window of Hippo pathway regulatory component availability.

**Figure 8.**
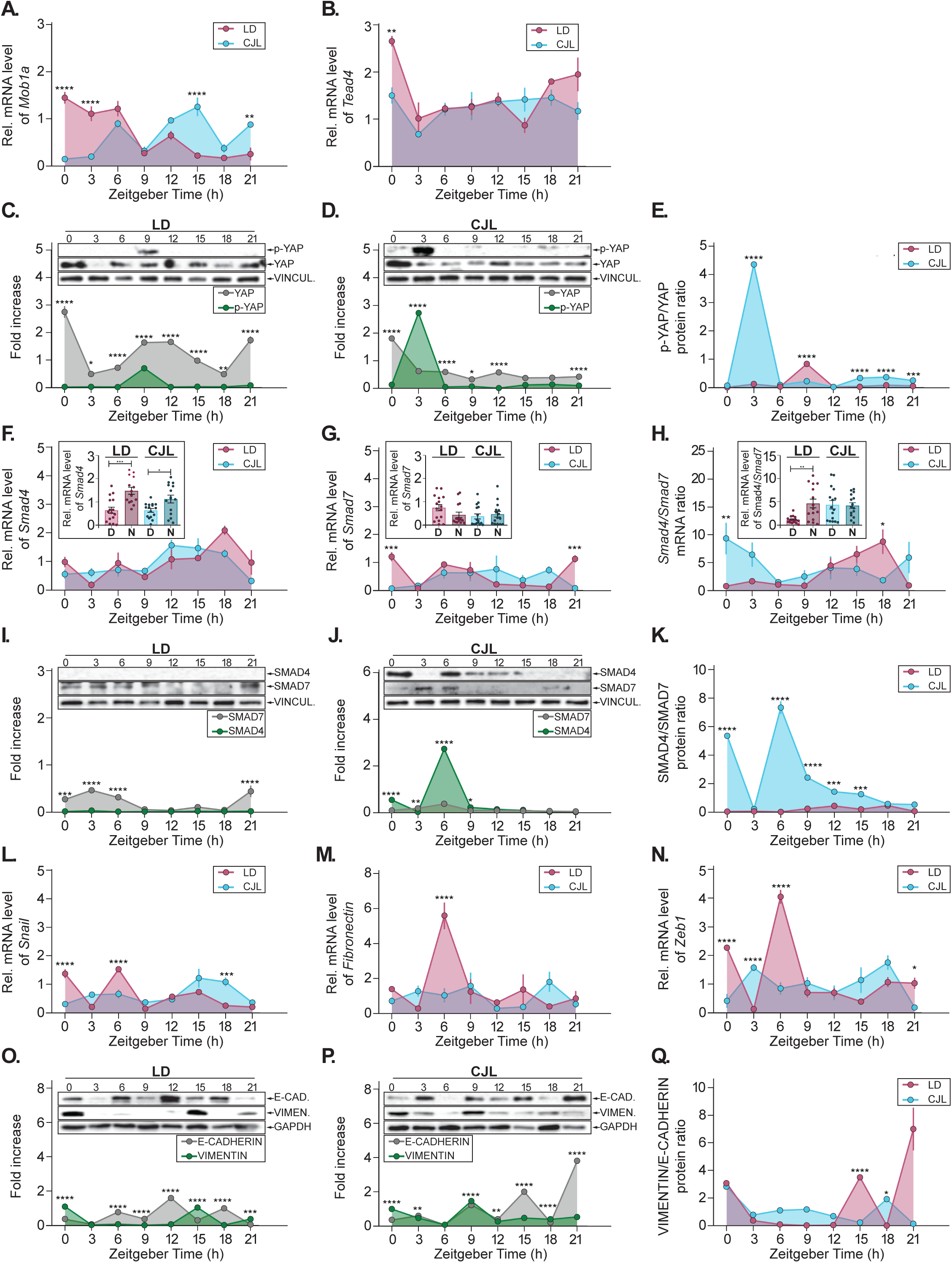
Metastatic burden amplifies Hippo and TGF-β pathway reorganization and EMT marker expression under circadian disruption. Relative mRNA levels of *Mob1a* (**A**) and *Tead4* (**B**) quantified by RT-qPCR in metastasis-bearing lung tissue from mice maintained under LD 12:12 (pink) or CJL (cyan) and harvested 21 days post-B16F10 inoculation at various zeitgeber times. Purple shading indicates overlapping values. Representative Western blots (top) and densitometric quantification (bottom) of phospho-YAP (Ser127, inactive cytoplasmic form) and total YAP proteins in metastatic lung tissue from LD (**C**) or CJL (**D**) mice, and p-YAP/YAP ratio (**E**) in mice harvested at 3h intervals. Quantification shows fold-increase relative to VINCULIN for total YAP (gray) and phospho-YAP (green). Note altered temporal patterns compared to non-tumor-bearing lungs (Figure 5). Relative mRNA levels of *Smad4* (**F**) and *Smad7* (**G**) quantified by RT-qPCR, and *Smad4*/*Smad7* mRNA ratio (**H**; Bar plots inside each panel indicate day-night analysis) in metastasis-bearing lung tissue from mice maintained under LD 12:12 (pink) or CJL (cyan) and harvested 21 days post-B16F10 inoculation. Representative Western blots (top) and densitometric quantification (bottom) of SMAD4 and SMAD7 proteins under LD (**I**) or CJL (**J**), with SMAD4/SMAD7 protein ratio comparison (**K**). VINCULIN serves as loading control. Note amplified CJL effects compared to non-tumor-bearing lungs (Figure 5). Relative mRNA levels of EMT transcription factors *Snail* (**L**), *Fibronectin* (**M**), and *Zeb1* (**N**) quantified by RT-qPCR in metastasis-bearing lung tissue from mice maintained under LD 12:12 (pink) or CJL (cyan) and harvested 21 days post-B16F10 inoculation at zeitgeber times. Purple shading indicates overlapping values. Representative Western blots (top) and densitometric quantification (bottom) of E-CADHERIN (epithelial marker, **O-P**, gray line) and VIMENTIN (mesenchymal marker, **O-P**, green line) proteins in metastatic lung tissue from LD (**O**) or CJL (**P**) mice and VIMENTIN/E-CADHERIN ratio (**Q**) in mice harvested at 3-hour intervals. GAPDH serves as loading control. Note amplified mesenchymal marker expression compared to non-tumor-bearing lungs (Figure 5). Data represent mean ± SEM from n=4 animals per timepoint. Two-way ANOVA for lighting condition: (**M**) p=0.0024, (**N**) p=0.0313; for protein type: (**D**) p=0.0127, (**C**, **I-K**, **O-P**) p<0.0001; for zeitgeber time: (**A-D**, **I-K**, **L-N**) p<0.0001. Mann-Whitney test (day vs. night bar graphs): (**B**) ***p=0.0004, (**C**) *p=0.033, (**H**) **p=0.0036. Post-hoc: *p<0.05, **p<0.01, ***p<0.001, ****p<0.0001.

*Tead4* mRNA expression patterns revealed more dramatic alterations (Figure 8B). Whereas non-tumor-bearing lungs showed modest expression under LD with CJL-induced elevation (Figure 5B), metastatic lungs exhibited pronounced elevation under both conditions, with LD showing higher level at ZT18-ZT0 (Figure 8B, pink line) and CJL producing sustained expression throughout most timepoints (ZT0-21; Figure 8B, cyan line). This constitutive elevation of YAP’s primary transcriptional partner suggests that metastatic burden creates conditions favoring enhanced YAP/TEAD transcriptional capacity.

Protein-level analysis of YAP revealed fundamental reorganization of temporal dynamics in metastatic tissue. In non-tumor-bearing LD lungs, phospho-YAP peaked at the end of the day (ZT12) with total YAP elevated during the night/day transition (ZT18-ZT3), creating nighttime YAP transcriptional activity (Figure 5C). Metastatic LD lungs showed altered patterns: phospho-YAP exhibited elevation only at ZT9 followed by near-undetectable levels throughout the remaining timepoints (Figure 8C, green line), while total YAP showed pronounced and sustained expression with particularly high levels at ZT18-0 and ZT6-15 (Figure 8C; gray line). The phospho-YAP/YAP ratio indicates an extended window of potential YAP activity compared to non-tumor-bearing tissue (Figure 8E vs. 5E, pink lines).

Under CJL conditions, the reorganization was even more pronounced. Non-tumor-bearing CJL lungs showed inverted YAP activity windows (day rather than night, Figure 5E), but metastatic CJL lungs exhibited early phospho-YAP peak (ZT3) followed by sustained low levels throughout day and night (Figure 8D, green line), while total YAP peaked at ZT0 and remained elevated particularly at ZT6-21 (Figure 8D, gray line). This pattern suggests that metastatic burden under CJL creates prolonged windows of YAP transcriptional competence extending through both day and night phases (Figure 8E, cyan line).

This constitutive YAP/TEAD activity amplifies the temporal reorganization observed in non-tumor-bearing tissue (Figure 5), providing mechanotransduction support for the feed-forward cycles characterized in Figure 7.

### Metastatic burden amplifies TGF-β pathway temporal reorganization under circadian disruption

Having characterized TGF-β pathway temporal reorganization in non-tumor-bearing lungs (Figures 5F-5K), we examined whether metastatic burden alters or amplifies these circadian disruption effects.

*Smad4* mRNA expression in metastatic lungs showed temporal patterns under both LD and CJL conditions (Figure 8F), but with enhanced amplitude compared to non-tumor-bearing tissue (compare to Figure 8F). Day-night analysis revealed significantly elevated *Smad4* during night under both LD and CJL conditions (Figure 8F, light vs. dark pink bars ***p=0.0004, light cyan vs. dark cyan bars *p=0.033, respectively), representing amplification of the temporal expansion observed in non-tumor-bearing CJL lungs. This suggests that metastatic burden enhances the availability of TGF-β pathway activator components.

*Smad7* mRNA displayed circadian oscillation with patterns distinct from non-tumor-bearing tissue (Figure 8G vs. Figure 5G). Whereas non-tumor-bearing CJL lungs showed pronounced *Smad7* peaks (Figure 5G), metastatic lungs exhibited more variable expression with reduced amplitude oscillations under both conditions. Day-night analysis confirmed relatively stable *Smad7* levels under both LD and CJL in metastatic lungs (Figure 8G, light vs. dark pink and light vs. dark cyan, n.s.), contrasting with the dramatic CJL-induced elevation observed in non-tumor-bearing tissue. This suggests that metastatic burden suppresses the negative regulatory component of TGF-β signaling.

The *Smad4*/*Smad7* mRNA ratio revealed an amplified temporal reorganization in metastatic tissue (Figure 8H). Whereas non-tumor-bearing lungs showed modest ratio shifts under CJL (Figure 5H), metastatic lungs exhibited pronounced elevation during night under LD conditions with dramatically altered temporal patterns under CJL. Day-night comparison confirmed significantly elevated ratios during night in LD metastatic lungs (Figure 8H, light pink vs. dark pink **p=0.0036), while CJL eliminated this temporal restriction (Figure 8H, light vs. dark cyan bars, n.s.), creating constitutive rather than temporally gated TGF-β signaling potential.

Protein-level analysis demonstrated even more pronounced reorganization in metastatic tissue. SMAD4 protein in metastatic LD lungs showed no expression throughout the day (Figure 8I, green line), while CJL conditions produced pronounced peaks at ZT0 and ZT6 (Figure 8J, green line). This contrasts with the more gradual changes observed in non-tumor-bearing tissue (Figures 5I-5J). SMAD7 protein displayed temporal variations with a peak at ZT6 under CJL (Figure 8P, gray line), while LD levels remained higher during night and early morning (ZT21-6, Figure 8O, gray line), representing amplification of the protein-level changes characterized in non-tumor-bearing lungs (Figures 5I-5J). Most significantly, the SMAD4/SMAD7 protein ratio in metastatic lungs exhibited dramatic temporal reorganization (Figure 8K). Under CJL, metastatic lungs showed elevated ratios with peaks at ZT0 and ZT6 and sustained elevation compared to LD throughout the time course, representing amplification of the temporal reorganization observed in non-tumor-bearing CJL lungs (Figure 5K). The absolute ratio values in metastatic CJL lungs exceeded those in non-tumor-bearing tissue, indicating that established tumors create conditions of enhanced TGF-β signaling capacity.

These findings demonstrate that metastatic burden amplifies TGF-β pathway temporal reorganization induced by circadian disruption, creating conditions maximally permissive for TGF-β signaling through enhanced *Smad4* availability, suppressed *Smad7* regulation, and elevated SMAD4/SMAD7 ratios. This amplified signaling capacity provides the molecular foundation for the *Tgf-β* elevation observed in metastatic CJL lungs (Figure 7L) and, combined with constitutive YAP activity (Figures 8A-8E), establishes feed-forward cycles wherein established metastases perpetuate the permissive conditions facilitating progressive metastatic burden.

### Metastatic burden amplifies EMT marker expression and eliminates temporal restriction under circadian disruption

Lastly, we examined how metastatic burden and circadian disruption affect EMT markers, the downstream effectors of the YAP (Figures. 5A-5E, 8A-8E) and TGF-β (Figures 5F-5K, 8F-8K) pathway reorganization characterized in this study. This analysis aimed to establish how established tumors maintain and amplify pro-metastatic conditions that facilitate progressive metastatic burden.

EMT transcription factors in metastatic lungs showed patterns distinct from non-tumor-bearing tissue (compare Figures 8L-8Q to Figures. 5L-5Q). *Snail* exhibited temporal variations under both LD and CJL conditions, but with reduced amplitude compared to non-tumor-bearing lungs (Figure 8L vs. Figure 5L). Most significantly, *Fibronectin* displayed a conspicuous peak under LD at ZT6, contrasting with the modest daytime levels observed in non-tumor-bearing LD lungs (Figure 8M vs. Figure 5M, pink lines). Under CJL, *Fibronectin* maintained modest steady level throughout the circadian cycle (Figure 8M, cyan line), contrasting with the nigh time elevation observed in non-tumor-bearing CJL tissue (ZT12-21); Figure 5M, cyan line). *Zeb1* showed peaks at ZT0 and ZT6 under LD in tumor-bearing animals (Figure 8N, pink line), within the same time window of expression observed in non-tumor-bearing animals under LD (ZT0-9, Figure 5N). Interestingly, sustained *Zeb1* expression was detected throughout the time course in tumor-bearing CJL animals (Figure 8N, cyan line), contrasting with the temporal window of expression observed in non-tumor-bearing CJL lungs (Figure 5N, cyan line). This temporal redistribution suggests that metastatic burden shifts EMT transcription factor expression toward constitutive rather than temporally restricted patterns.

Protein-level analysis revealed fundamental reorganization of epithelial-mesenchymal balance in metastatic tissue. E-CADHERIN expression in metastatic LD lungs showed jagged elevation with peaks and troughs from ZT0-21 every 6h (Figure 8O, gray line), contrasting with the early-day elevation (ZT0-ZT3) observed in non-tumor-bearing LD lungs (Figure 5O, gray line). Under CJL, E-CADHERIN displayed even more dramatic jagged elevation with progressive increase from ZT3-21 Figure 8P, gray line) that seems out of phase with the pattern observed in metastatic LD lungs (Figure 8O, gray line), representing a fundamental shift from the sustained expression pattern observed in non-tumor-bearing CJL tissue. Most significantly, VIMENTIN protein exhibited an amplification in metastatic lungs. Under LD conditions, VIMENTIN showed sustained low levels with modest elevation at ZT0 and ZT15 (Figure 8O, green line), while CJL produced pronounced elevation at ZT9 followed by sustained levels throughout day and night (Figure 8P, green line). The VIMENTIN/E-CADHERIN ratio confirmed a mesenchymal phenotype prominent at night in LD metastatic tissue, while CJL metastatic lungs showed sustained elevation throughout the cycle (Figure 8Q, pink vs. cyan lines). This represents amplification of the patterns observed in non-tumor-bearing lungs, which showed discrete peaks under LD and a shifted day-night elevation under CJL (Figure 5Q). Importantly, absolute VIMENTIN levels in metastatic CJL lungs exceeded all other conditions, indicating maximal mesenchymal marker expression.

The elimination of temporal restriction in both EMT transcription factor expression and mesenchymal marker accumulation demonstrates that established metastases under CJL do not merely exploit a permissive microenvironment but actively perpetuate it, sustaining the conditions that enabled their own colonization.

### Pathway convergence architecture is recapitulated in human metastatic melanoma

The coordinated temporal reorganization of pro-metastatic signaling pathways characterized in CJL-exposed mouse lungs raises the fundamental question of whether circadian clock integrity similarly shapes inter-pathway coupling architecture in human cancer. Although transcriptomic data cannot resolve circadian oscillatory dynamics directly, the ratio of positive to negative clock regulators within a tumor sample provides a tractable proxy for transcriptional clock architecture integrity: loss of BMAL1-driven transcription disrupts the coordinated expression balance that defines functional circadian oscillation, producing measurable disorganization of clock gene co-expression relationships ^73^. To interrogate this question and establish that the pathway convergence we characterize experimentally is not an epiphenomenon of the chronic jet lag paradigm, we performed a bioinformatic analysis of RNA-seq data from the TCGA-SKCM (The Cancer Genome Atlas Skin Cutaneous Melanoma) PanCancer Atlas.

Of 443 melanoma datasets retrieved, 76 primary tumor samples were excluded, yielding 367 metastatic specimens for downstream analysis (Figure 9A). Composite pathway activity scores were computed for MMP signaling, YAP/TAZ activity, TGF-β activators, EMT markers, and inflammatory mediators, alongside individual circadian clock gene expression. Samples were stratified by clock integrity on the basis of co-expression relationships among core clock components: functioning clock status required concordant ARNTL/NPAS2 expression inversely related to NR1D1, reflecting the canonical transcriptional architecture of an active molecular oscillator ^73^. This classification yielded 78 Metastatic/Functioning clock and 289 Metastatic/Disrupted clock samples (Figure 9A). Classification validity was confirmed using the Clock Correlation Distance (CCD) algorithm, which quantifies the regularity of circadian clock progression from clock gene co-expression structure, with higher scores indicating greater disorganization ^74^. The Metastatic/Functioning clock group exhibited a CCD of 3.925, reflecting relative integrity of the canonical clock gene network, whereas the Metastatic/Disrupted clock group displayed a markedly elevated CCD of 4.703, indicative of pervasive disruption of coordinated clock gene expression (ΔCCD = 0.778, p = 0.002; Figure 9B).

**Figure 9.**
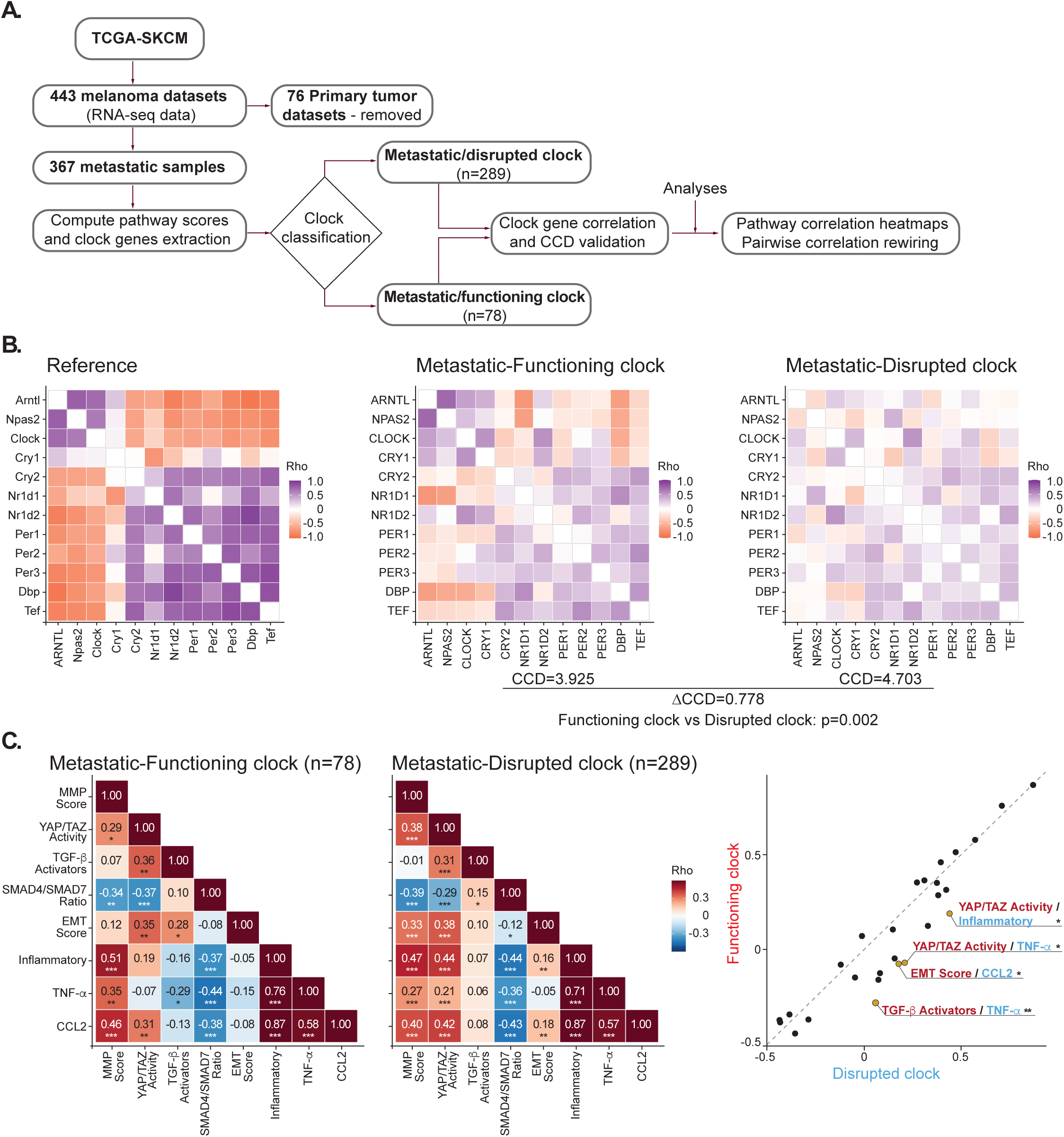
Circadian clock integrity determines inter-pathway coupling architecture in human metastatic melanoma. (**A**) Analytical pipeline applied to RNA-seq data retrieved from the TCGA-SKCM PanCancer Atlas. Of 443 melanoma datasets, 76 primary tumor samples were excluded, yielding 367 metastatic specimens subjected to downstream analysis. Composite pathway activity scores were computed and core circadian clock gene expression extracted; samples were subsequently stratified by clock integrity into two groups: Metastatic/Disrupted clock (n=289) and Metastatic/Functioning clock (n=78). Downstream analyses encompassed clock gene pairwise correlation matrices, Circadian Clock Disruption (CCD) score validation, pathway correlation heatmaps, and pairwise correlation rewiring. (**B**) Pairwise Spearman correlations among 12 core circadian clock genes (ARNTL, NPAS2, CLOCK, CRY1, CRY2, NR1D1, NR1D2, PER1, PER2, PER3, DBP, TEF) displayed as correlation heatmaps for a reference dataset (left), the Metastatic-Functioning clock group (center), and the Metastatic-Disrupted clock group (right). Tumors with functional clock coordination exhibited a lower CCD score (3.925), reflecting closer concordance with the reference clock gene correlation architecture, whereas clock-disrupted tumors displayed a markedly elevated CCD (4.703), indicative of pervasive disruption of circadian gene co-regulation (ΔCCD = 0.778; p = 0.002). Color scale: Spearman ρ from −1.0 (blue) to +1.0 (red). (**C**) Spearman correlation heatmaps of composite pathway activity scores, encompassing MMP signaling, YAP/TAZ activity, TGF-β activators, SMAD4/SMAD7 ratio, EMT markers, inflammatory signaling, TNF-α, and CCL2, computed for Metastatic-Functioning clock (left, n=78) and Metastatic-Disrupted clock (center, n=289) tumors. Full gene sets used to compute each score are provided in the Materials and Methods section. Asterisks denote statistical significance (*p<0.05, **p<0.01, ***p<0.001). The scatter plot (right) directly compares pairwise Spearman correlation coefficients between clock-disrupted (x-axis) and clock-functioning (y-axis) groups; each point represents a single pathway pair. Labeled pairs denote pathway combinations with statistically significant differences between clock states, as determined by Fisher z-transformation of correlation coefficients. Notably, all labeled significant pairs, YAP/TAZ Activity/Inflammatory, YAP/TAZ Activity/TNF-α, EMT Score/CCL2, and TGF-β Activators/TNF-α, fall below the identity diagonal, indicating that these pathway couplings are selectively strengthened in clock-disrupted tumors, consistent with the pathway convergence model characterized experimentally in Figures 3–8.

Having confirmed the biological validity of the classification, we examined whether clock status was associated with differences in inter-pathway coupling by computing pairwise Spearman correlation matrices across the pathway variables found to be temporally reorganized in CJL-exposed mouse lungs (Figure 9C, left and middle panels). In the Metastatic/Functioning clock group, several pathway pairs exhibited significant positive correlations, including MMP Score with Inflammatory signaling (ρ = 0.51, p < 0.001) and CCL2 (ρ = 0.46, p < 0.001), and YAP/TAZ Activity with both TGF-β Activators (ρ = 0.36, p < 0.01) and EMT Score (ρ = 0.35, p < 0.01). Of note, the SMAD4/SMAD7 ratio showed significant negative correlations with YAP/TAZ Activity (ρ = −0.37, p < 0.01) and Inflammatory signaling (ρ = −0.37, p < 0.01) in the functioning clock group, a pattern substantially attenuated in clock-disrupted tumors, suggesting that intact clock architecture maintains inhibitory constraints on TGF-β pathway output relative to mechanotransduction and inflammatory programs, mirroring the temporal segregation characterized in LD mouse lungs. As anticipated from the dominant role of inflammatory axis coupling in the metastatic niche, TNF-α and CCL2 converged strongly in both groups (Functioning: ρ = 0.76, p < 0.001; Disrupted: ρ = 0.71, p < 0.001), indicating that this core inflammatory relationship is preserved independently of clock status.

To systematically identify selectively rewired pathway relationships, all 28 pairwise pathway correlations were compared between groups using Fisher z-transformed correlation coefficients. The resulting scatter plot revealed that the majority of pathway pairs clustered near the identity diagonal, indicating broad conservation of overall coupling architecture across clock groups (Figure 9C, right panel). This conservation is consistent with the notion that large-scale pathway rewiring may precede the established metastatic stage sampled in the TCGA cohort. However, given the cross-sectional design nature of these data, the precise timing of such rewiring cannot be formally determined from this analysis alone. Nonetheless, four pathway pairs were significantly and selectively rewired in clock-disrupted tumors: YAP/TAZ Activity exhibited stronger positive coupling with both Inflammatory signaling and TNF-α (p < 0.05 for each), while EMT Score/CCL2 (p < 0.05) and TGF-β Activators/TNF-α (p < 0.01) correlations were similarly strengthened in the Metastatic/Disrupted clock group relative to functioning clock tumors. All four significantly rewired pairs fell below the identity diagonal, confirming directional strengthening of these couplings specifically in the clock-disrupted context. Together, these findings provide direct translational validation of the pathway convergence model established in CJL-exposed mouse lungs, demonstrating that circadian clock integrity shapes pro-metastatic signaling architecture in human disease.

## DISCUSSION

The findings presented in this study suggest that the temporal architecture of the pulmonary microenvironment, rather than individual pathway activity, is a primary determinant of metastatic permissiveness, with circadian disruption acting as the mechanism through which this organizational framework is systematically dismantled. Our comprehensive analysis, summarized in Figure 10, reveals a two-phase progression consisting of foundational pathway reorganization that creates permissive conditions for enhanced colonization, followed by progressive amplification, wherein established metastatic burden perpetuates and extends these pro-metastatic programs through self-reinforcing cycles. Together, these findings position circadian clock integrity as a critical determinant of pulmonary metastatic susceptibility and, as validated in human TCGA-SKCM data, a conserved feature of pro-metastatic signaling architecture in human melanoma.

**Figure 10.**
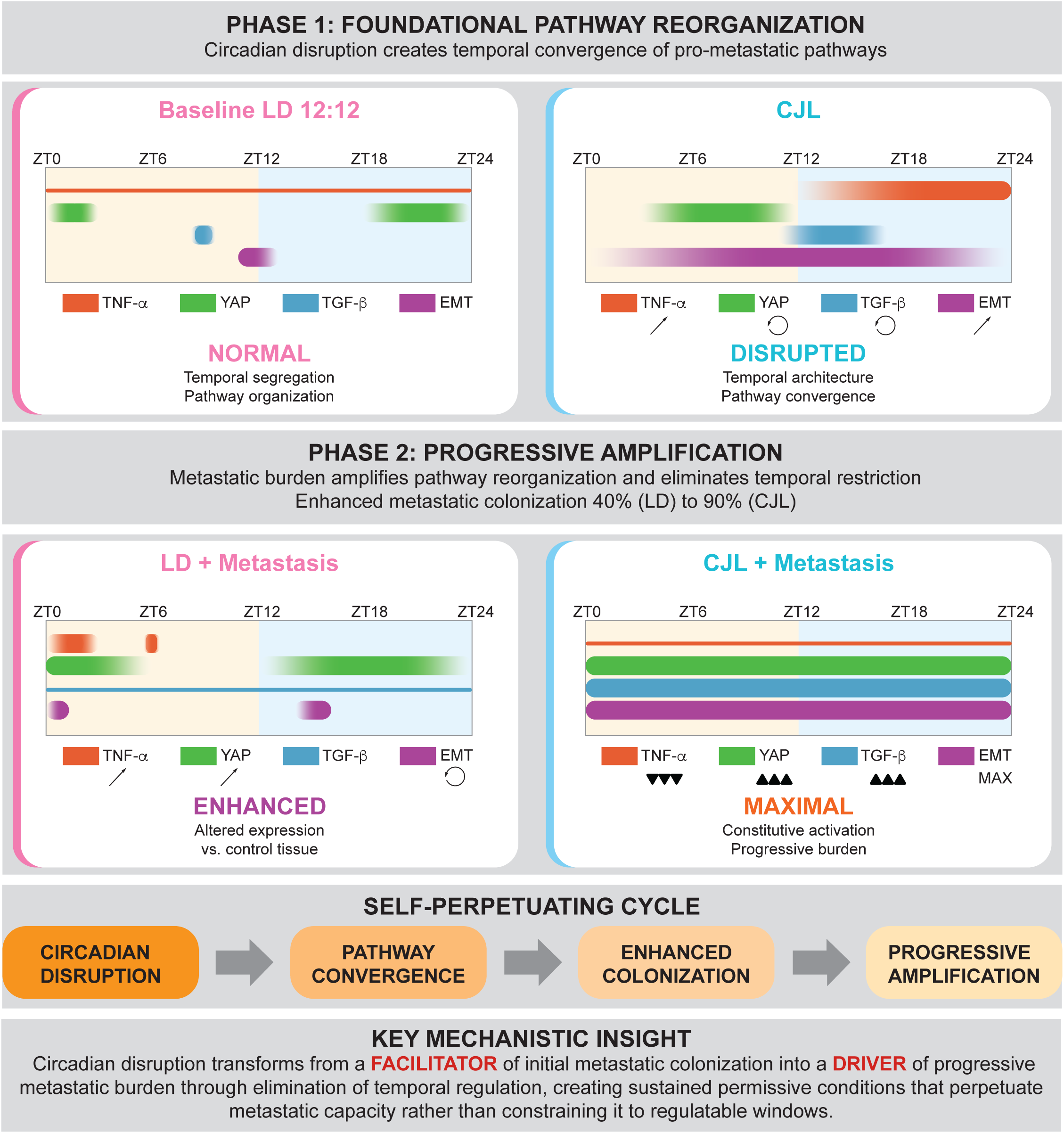
Circadian disruption transforms from facilitator to driver of progressive metastatic burden through elimination of temporal regulatory architecture. Phase 1: Foundational Pathway Reorganization. Temporal pathway activity patterns in non-tumor-bearing lung tissue showing baseline LD 12:12 conditions with normal temporal segregation of TNF-α (red), YAP (green), TGF-β (blue), and EMT (purple) pathways versus CJL-induced disruption creating pathway convergence and temporal reorganization. Yellow background indicates light phase (ZT0-12); blue background indicates dark phase (ZT12-24). Enhanced metastatic colonization (Figure 10) demonstrates translation of molecular pathway convergence to increased metastatic success: 40% incidence under LD conditions versus 90% incidence under CJL conditions following B16F10 melanoma inoculation. Phase 2: Progressive Amplification. Temporal pathway activity in established metastatic tissue harvested 21 days post-tumor inoculation, showing LD + metastasis conditions with enhanced expression versus control tissue, and CJL + metastasis conditions with maximal constitutive pathway activation eliminating temporal restriction. Pathway symbols indicate expression levels: normal (baseline), ↗ (elevated), ⟲ (temporally reorganized), ▴▴▴ (maximally amplified). This progression establishes that circadian disruption eliminates temporal barriers to create self-perpetuating cycles wherein pathway convergence facilitates initial colonization and established metastases amplify pro-metastatic signaling to maintain sustained permissive microenvironmental conditions, transforming circadian disruption from a facilitator of initial metastatic events into a driver of progressive metastatic burden.

The present results are consonant with, and mechanistically extend, a body of evidence establishing circadian clocks as temporal gatekeepers of metastatic competence. Chronic jet lag promotes breast cancer dissemination and lung metastasis through immunosuppressive TME remodeling and enhanced stemness ^44^, and experimental disruption accelerates metastasis of transplanted and carcinogen-induced tumors through immune microenvironment remodeling ^75,76^. The generation of circulating tumor cells peaks during the rest phase, establishing that metastatic capacity is itself time-of-day-restricted ^77^. At the molecular level, BMAL1 loss activates a PAI-1–plasmin–TGF-β cascade that drives myofibroblastic CAF differentiation and metastatic niche formation ^78^, and, reciprocally, TGF-β coordinates circadian phase synchronization between neighboring cells through SMAD-dependent clock gene regulation ^79^, establishing a bidirectional link between clock integrity and TGF-β pathway activity. This bidirectional relationship provides a mechanistic basis for understanding why TGF-β dysregulation in CJL-exposed lung tissue is simultaneously a consequence and a perpetuator of circadian disorganization, and illustrates a broader principle wherein clock disruption and pro-metastatic pathway activation are mutually reinforcing rather than causally sequential. Consistent with this principle, altered clock organization in tumors correlates with EMT cycling and metastatic potential ^80^, and circadian clocks rhythmically regulate EMT, ECM remodeling, and tumor-microenvironment interactions across the metastatic cascade ^81–83^.

Prior studies have identified circadian regulation of individual metastasis-associated pathways, including matrix metalloproteases, inflammatory mediators, and growth factors ^48,84–89^. The present findings reframe this picture by indicating that individual pathway regulation is not sufficient to explain metastatic susceptibility; rather, what matters is the temporal segregation that prevents these programs from converging simultaneously. YAP/TEAD emerges not merely as a downstream effector but as an obligate integration node where clock-regulated ECM mechanical signals and cytokine-driven transcriptional programs converge, thereby representing the point at which temporal disorganization translates into metastatic competence. This convergence architecture is conserved in clock-disrupted human metastatic melanoma (Figure 9), suggesting that it may generalize beyond the experimental model. The amplified SMAD4/SMAD7 ratios and constitutive VIMENTIN elevation in metastatic CJL lungs further raise the possibility that circadian pathway signatures, rather than tumor burden alone, could serve as prognostic indicators of metastatic progression rate, a hypothesis warranting prospective evaluation.

The temporal specificity of the pathways characterized here has direct therapeutic implications, converging with clinical evidence for chronotherapy as an actionable intervention. The first prospective randomized phase 3 trial of immunotherapy chronotherapy demonstrated that anti-PD-1 plus chemotherapy administered in the morning more than doubled progression-free and overall survival in NSCLC ^90^. This effect is mechanistically grounded in the circadian regulation of CD8+ tumor-infiltrating lymphocyte recruitment driven by rhythmic ICAM-1 expression on tumor endothelium ^71^, a principle directly reflected in the M1/M2 macrophage temporal disorganization we characterize under CJL, wherein the elimination of time-of-day-dependent immune programs would be expected to abrogate the circadian window of maximal anti-tumor immune competence. In melanoma, retrospective analyses demonstrate that late-afternoon ICI infusions are associated with significantly worse overall survival ^91^. The temporal concentration of TGF-β elevation and YAP activity in metastatic CJL lungs at defined zeitgeber windows suggests that timing TGF-β or YAP/TEAD inhibitors to coincide with peak pathway activity could improve therapeutic index, a chronotherapeutic opportunity currently absent from the literature for both drug classes, despite active clinical development of TEAD inhibitors ^92,93^. Clock restoration strategies, including REV-ERB agonists ^94^, CRY stabilizers that suppress fibrocyte-to-CAF differentiation and overcome immune exclusion ^95^, and time-restricted feeding that reduces lung metastasis through restoration of diurnal gene expression rhythms ^96^, offer complementary approaches targeting the temporal architecture whose loss drives the amplification cycles characterized here. These considerations are clinically relevant to the estimated 20% of the industrialized workforce engaged in shift work, classified as Group 2A probable carcinogens by the IARC ^97,98^, and to the broader population experiencing social jet lag, for whom circadian disruption may be actively driving progressive metastatic burden rather than merely predisposing to cancer initiation ^99^.

Our analysis presents both strengths and opportunities for future investigation. Whole-lung tissue analysis enabled comprehensive assessment of coordinated pathway dynamics across the pulmonary microenvironment; the temporal concordance across independent pathway readouts (TNF-α/MMPs, YAP/SMAD ratios, EMT markers) and stable total macrophage counts (Figure S5) support microenvironmental rather than tumor-derived origins. Future studies integrating systemic endocrine and autonomic measurements will determine whether the pathway convergence we observe reflects tissue-autonomous coupling or coordinated responses to common upstream drivers altered by chronic jet lag. Examination of additional cancer types, metastatic sites, and clinically relevant circadian disruption patterns, including shift work, social jet lag, and age-related circadian decline, will establish the generalizability of the temporal architecture framework across the metastatic cascade. Longitudinal cohort studies with objective circadian monitoring will be needed to determine whether clock disruption precedes metastatic progression or is amplified by established tumor burden, a distinction with direct implications for preventive versus therapeutic intervention windows. Together, these findings position circadian rhythm restoration and chronotherapeutic approaches as rational interventions for metastatic disease, grounded in the mechanistic principle that temporal architecture is an active constraint on metastatic capacity whose elimination is measurable and potentially reversible.

## METHODS

### Cell culture

The lung fibroblast-like MLg cells and melanoma B16F10 cells were purchased from ATCC (CCL-206, CRL-6475, respectively) and cultured in complete Essential Modified Eagle’s Medium (EMEM; ATCC) supplemented with 10% (v/v) fetal bovine serum (FBS) and maintained at 37°C in a humidified incubator with 5% CO2. NIH 3T3 and NIH PER2KO cell lines were cultured in complete Dulbecco Modified Eagle’s Medium (DMEM; ATCC) supplemented with 10% (v/v) fetal bovine serum (FBS) and maintained at 37°C in a humidified incubator with 5% CO2.

### Bmal1 shRNA-mediated knockdown

For Bmal1 knockdown, 80% confluent MLg cells were incubated with a suspension containing 10 pmol of ON-TARGETplus mouse Arntl (1165) siRNA-SMART pool (Dharmacon) or ON-TARGET-plus non-targeting pool (Dharmacon), 150 µL of OPTI-MEM medium and 9 µL of Lipofectamine RNAiMAX (Thermo Fisher) according to manufacturer’s instructions. After 48 h of incubation, samples were collected and knockdown of Bmal1 was confirmed by qPCR and Western blot (following the protocols described below for each technique).

### Wound healing assays

MLg and NIH 3T3 migration was detected by wound healing assay. MLg cells were inoculated in 6-well plates at 7×10^5^ cells/well. Cells were synchronized in the presence of dexamethasone (100 nM) for 1 h when needed, and Mitomycin C (20 µg/ml) was added to prevent proliferation. Thereafter, three parallel scratches were made in each well using the tip of a sterile plastic microtube and EGF (10 ng/ml), FGF2 (20 µM; SinoBiological), TGF-β (10 ng/ml; R&D Systems), TNF-α (2 ng/ml; R&D Systems), GSK2945 (20 µM; MedChemExpress), KL001 (3 µM, Cayman Chemicals), PF670462 (1 µM, Cayman Chemicals) or verteporfin (20 µM; Aobious) was added at the beginning of the assay (Time = 0 h post synchronization), unless specified in the figure legend. Four images per scratch were captured every 6 h for 24 h and analyzed using ImageJ to quantify the wound area. Experiments were performed in duplicate, and each experiment was repeated twice independently.

### Animals

Young adult 2-month-old C57bl/6J wild type (WT) male mice were raised in our colony. All mice were housed in groups under a 12:12-h light-dark (LD) photoperiod (with lights on at 7 AM and lights off at 7 PM) with food and water ad libitum for 2 weeks before entering experimental conditions. This study was carried out in accordance with international ethical standards for the care and use of laboratory animals. The protocol was approved by the Institutional Animal Care and Use Committee of the University of Quilmes, Argentina.

### Experimental lighting schedules

Individually caged animals were maintained for 21 days under LD12:12 or Chronic Jet Lag [CJL, 6 h advance of the LD12:12 cycle every 2 days, shortening of every second dark phase ^61^] schedules with food and water ad libitum. Zeitgeber time (ZT) was used as a temporal reference: ZT0 is the moment of lights on, and ZT12 the moment of lights off. For the CJL schedule, samples were taken during the 12-h dark phase and during the light phase. Effective circadian desynchronization was confirmed by individually monitoring locomotor activity rhythms using infrared sensors. As expected, CJL-maintained animals showed two components of activity rhythms with periods of ∼21h and 24.7h as previously described ^61^.

### *In vivo* experimental design

As previously mentioned, mice were housed under either an LD or CJL schedule for 3 weeks prior to being inoculated with B16F10 tumor cells, or vehicle (DMEM), and maintained under the corresponding light schedule for another 3 weeks (end point of the tumorigenic protocol). To induce metastasis, 200,000 cells were inoculated intravenously in the lateral tail vein. Injections of all animals were carried out during a 2h window, ZT6 to ZT8, a timeframe at which the light phase (which followed a 12h-night) in CJL overlapped with that of the LD. After 21 days, mice were sacrificed at various ZTs (ZT0, ZT3, ZT6, ZT9, ZT12, ZT15, ZT18, and ZT21) and lung tissues were carefully dissected and washed with distilled water. The same lung lobule was used in each experimental technique (RT-qPCR, Western blot and Flow Cytometry). Metastatic foci size and the percentage of total lung area occupied by metastases were quantified using ImageJ software.

### RNA extraction and Real-time PCR

For MLg cells, total RNA was isolated using 500 μl of TRIzol reagent (Life Technologies). Next, an equal ethanol (100%) volume was added and RNA purification was performed using Direct-zol^TM^ RNA mini prep (Zymo Research) according to manufacturer’s instructions. For lung tissue, total RNA was isolated using 300 μl of TRIzol reagent (Life Technologies) according to the manufacturer’s instructions until the aqueous phase was obtained. Next, an equal ethanol (96%) volume was added to the aqueous phase (1:1) and RNA purification was performed using Direct-zol^TM^-96 RNA (Zymo Research) according to the manufacturer’s instructions. RNA samples were quantified using NanoDrop1000 (Thermo Fisher Scientific) and their integrity was evaluated by electrophoresis. One μg of total RNA was treated with RQ1 RNAse-Free DNAse (Promega), and cDNA was synthesized using LunaScript® RT SuperMix Kit (New England BioLabs^TM^ Inc.) for MLg cells RNA and oligo(dT) primers and the SuperScript™ First-Strand Synthesis System (Invitrogen) for lung RNA samples.

Gene amplification was performed on a CFX Opus 384 instrument (Applied Biosystems), using 10 μl of final reaction volume containing 2 μl of cDNA as the template, 1× of the PowerUp^TM^ SYBR^TM^ Green Master Mix (Applied Biosystems), and the corresponding primers (Supplemental Table 7) at a final concentration of 400 nM. The cDNA template was amplified in triplicate, with the following conditions: 95°C for 2 min, followed by 45 cycles of 95°C for 15 s and 60°C for 1 min. Relative gene expression was calculated using the 2^−ΔΔCt^ method, and *Hprt* was used as the reference housekeeping gene.

### Protein extraction

Lung tissues were mechanically homogenized in protein extraction buffer [10 mM Tris-HCl, 137 mM NaCl, 1 mM EDTA, 10% glycerol, 80 mM β-glycerophosphate, 1 mM Na_3_VO_4_, 10 mM NaF and 1× protease inhibitors cocktail (Sigma Aldrich)]. Next, 0.5% NP-40 was added, incubated for 15 min at 4°C, and centrifuged at 14,000 g for 15 min. Supernatants were collected and protein concentration was measured by Bio-Rad Protein Assay (Bio-Rad). Briefly, 200 μl of 1:5 diluted dye reagent was mixed with 10 μl of each sample and absorbance was measured at 595 nm. Bovine Serum Albumin (BSA) was used as a protein standard.

### Western blotting

Fifty μg of protein samples were loaded onto 12% polyacrylamide gel. After electrophoretic separation, proteins were transferred to a PVDF membrane. The membranes were blocked with 5% milk powder in Tris-buffered saline with 0.1% Tween-20 detergent (TBST). The following primary antibodies were used: YAP-1 (Cell Signaling #8418), p-YAP (Cell Signaling #4911), SMAD7 (Abcam #ab216428), SMAD4 (Cell Signaling #38454), E-CADHERIN (Cell Signaling #3195T), VIMENTIN (Cell Signaling #5741T), VINCULIN (Cell signaling #13901T) and GAPDH (AbClonal #AC002), in combination with the following secondary antibodies: Goat anti-rabbit HRP (Invitrogen #A16104) and Goat anti-mouse (Invitrogen #A16072). Membranes were digitally imaged using ChemiDOC MP (Bio-Rad) and analyzed with ImageLab Software (Bio-Rad).

### Flow cytometry

Lung tissues were collected in culture medium containing 1 mg/ml Collagenase IV (Thermo Fisher Scientific). Following mechanical disruption, tissues were maintained at 37°C for 20 min before the collagenase activity was inhibited by adding 15% fetal bovine serum. Next, cells were filtered with a 70 μm filter and incubated for 5 min with lysis buffer to eliminate red blood cells (NH4Cl 8290 μg/ml, KHCO3 1 μg/ml and EDTA 1mM) in a 1:9 ratio, and then centrifuged at 400 g for 10 min at 4°C. About 1×10^6^ cells were blocked with mouse normal serum diluted 1:5 for 15 min and incubated with the indicated fluorescent antibody (F4/80^+^, CD86, and CD206; Biolegend), or their corresponding isotype control (PE rat IgG2a, PerCP7cy5.5 rat IgG2a, APC rat IgG2a; Biolegend), for 30 min at room temperature. Of note, *Mrc1* is the gene encoding CD206, a surface protein marker used to identify M2 macrophages. Cells were then washed with 3% PBS-FBS, centrifuged at 400 g for 10 min, and fixed. Fluorescence detection was carried out using a flow cytometer FACSCalibur^TM^ (Becton Dickinson). Data were analyzed using the FlowJo 7.6 software. For M1 and M2 macrophage analysis, the F4/80^+^ (macrophage marker) population was plotted for CD86 and CD206 markers, and CD86^+^CD206^-^ cells were considered M1 and CD86^-^CD206^+^ were considered M2 (similar percentage of CD86^+^CD206^+^ population was observed in all group of samples]. Additionally, the M1/M2 ratio was calculated in each sample.

### Single cell RNAseq analysis

All data were sourced from the publicly available dataset GSE260641 ^71^. Macrophages were identified from the single-cell RNA-seq dataset (a Seurat object in R v4.x) based on prior cell-type annotation and subset for downstream analysis. Cells were classified into four categories according to the expression of *Cd86* and *Mrc1* (encodes for CD206): *Cd86⁺*, *Mrc1⁺*, *Cd86⁺*/Mrc1⁺ (double-positive), and *Cd86⁻*/*Mrc1⁻* (negative). Positivity was defined using SCTransform-normalized expression values, with a threshold of >1 for each gene. For visualization, Uniform Manifold Approximation and Projection (UMAP) coordinates previously computed for the full dataset were reused without recomputation. For each time point (ZT01, ZT07, ZT13, ZT19), UMAPs were generated independently to display macrophage distributions. Two panels were shown per time point: one highlighting *Cd86⁺* and *Mrc1⁺* cells against the remaining macrophage population, and a second highlighting *Cd86⁺*/*Mrc1⁺* cells against the remaining macrophages. Cells not belonging to the highlighted category were retained as background to preserve spatial context. All UMAPs were displayed using identical axis limits, point size, and color schemes to allow direct visual comparison across time points. Differential gene expression (DGE) analysis was performed to characterize transcriptional differences between macrophage subpopulations defined as M1 or M2 macrophages by *Cd86* and *Mrc1* expression, respectively. Within each time point (ZT01, ZT07, ZT13, ZT19), macrophages were grouped into *Cd86⁺*, *Mrc1⁺*, and *Cd86⁺*/*Mrc1⁺* populations, and each group was compared against all remaining macrophages from the same time point. DGE was conducted using the Seurat FindMarkers function with the Wilcoxon rank-sum test, using log-normalized RNA expression values. Genes were ranked by average log2 fold change, and statistical significance was assessed using Bonferroni-adjusted p-values. For each comparison and time point, results were compiled into a unified table including p-values, adjusted p-values, average log2 fold change, and the percentage of cells expressing each gene in the target and reference populations. Differential gene expression results were visualized using dot plots generated in R (v4.x) for each macrophage population (M1: *Cd86⁺*, M2: *Mrc1⁺*, and *Cd86^+^*/*Mrc1^+^*). Dot size represents statistical significance (p-value), while dot color encodes magnitude of expression change (log₂ fold change).

### Bioinformatic analysis

RNA-seq expression data, sample metadata, and clinical annotations were retrieved from cBioPortal using the TCGA-SKCM PanCancer Atlas dataset (skcm_tcga_pan_can_atlas_2018), encompassing z-score-normalized and RSEM expression values at both the sample and patient levels. The processed expression dataset comprised 443 samples in total; downstream analyses were restricted to metastatic specimens based on clinical annotation, yielding 367 samples for subsequent classification and pathway analysis.

Composite pathway activity scores were computed from z-score expression matrices as the mean expression of predefined gene sets, defined as follows: MMP signaling (*MMP2*, *MMP3*, *MMP9*, *MMP13*, *MMP19*); YAP/TAZ activity [*CTGF*, *CYR61*, *AMOTL2*, *ANKRD1*, *AXL*, *BIRC5*, *CCND1*, *CDH2*, *CRIM1*, *EDN1*, *FOXF2*, *GADD45A*, *IGFBP3*, *INHBA*, *ITGB2*, *LOXL2*, *MYOF*, *NT5E*, *NUAK2*, *SERPINE1*, *TGFB2*, *THBS1*; ^100^]; TGF-β activators (*TGFB1*, *TGFB2*, *TGFB3*, *SMAD2*, *SMAD3*, *SMAD4*, *SMAD7*, *TGFBR1*, *TGFBR2*); EMT markers (*CDH1*, *CDH2*, *VIM*, *FN1*, *SNAI1*, *SNAI2*, *ZEB1*, *ZEB2*, *TWIST1*, *OCLN*, *TJP1*); and inflammatory signaling (*TNF*, *CCL2*, *IL1B*, *IL6*, *IL10*, *CXCL9*, *CXCL10*, *IFNG*). The SMAD4/SMAD7 ratio was calculated as a proxy for TGF-β signaling balance, capturing the relative contributions of pathway activation and inhibitory feedback.

Distinct signaling states were defined by the relative expression of these components: high SMAD4 with low SMAD7 indicates active signaling; high SMAD4 with high SMAD7 reflects feedback-constrained activation; and low SMAD4 with high SMAD7 indicates pathway suppression. This ratio was therefore analyzed as an independent pathway score rather than incorporated into the TGF-β activator composite, as it captures directional signaling capacity not resolved by individual gene expression levels. Individual core clock gene expression values (*ARNTL*, *NPAS2*, *CLOCK*, *CRY1*, *CRY2*, *NR1D1*, *NR1D2*, *PER1*, *PER2*, *PER3*, *DBP*, *TEF*) were extracted to generate clock network correlation matrices, visualized as previously described ^74^.

Circadian clock integrity was defined on the basis of expression relationships among core clock components: samples were classified as clock-functioning when *ARNTL* and *NPAS2* were concordantly expressed and inversely related to *NR1D1* (i.e., *ARNTL* > 0, *NPAS2* > 0, *NR1D1* < 0, or *ARNTL* < 0, *NPAS2* < 0, *NR1D1* > 0 in z-score space), and as clock-disrupted otherwise ^73^. This classification yielded 78 samples in the Metastatic/Functioning clock group and 289 in the Metastatic/Disrupted clock group. Disruption of clock gene coordination was quantified using the Clock Correlation Distance (CCD) metric and the delta CCD (ΔCCD) method, as previously described ^74^.

Within each clock group, pairwise Spearman correlation matrices were computed across all pathway-level variables (MMP signaling, YAP/TAZ activity, TGF-β activators, SMAD4/SMAD7 ratio, EMT score, inflammatory signaling, TNF-α, CCL2) and visualized as heatmaps. Inter-group differences in pairwise pathway correlations were evaluated across all 28 possible pathway edges (derived from 8 pathway variables); for each edge, the difference in Spearman correlation between the Metastatic/Functioning clock and Metastatic/Disrupted clock groups was tested using Fisher z-transformation, with correction for group sample size.

Mutational annotations for canonical melanoma driver genes (BRAF, NRAS, NF1) were available within the TCGA-SKCM dataset but were deliberately excluded from the primary analysis. As the central argument of this study concerns circadian disruption as a microenvironmental modifier of metastatic permissiveness, rather than the oncogenic drivers that initiate melanoma, stratification by driver genotype was not warranted in this context. The extent to which clock disruption-associated pathway rewiring operates independently of driver mutation status represents an important question that may be addressed in future sensitivity analyses.

### Statistical analysis

Data are presented as the mean ± standard deviation (SD) or as the mean ± standard error of the mean (SEM). For rhythmicity analysis, Meta2D function of MetaCycle was used with a minimum period of 20 h and maximum period of 28 h ^101^. Differences between two groups were analyzed using the unpaired Student’s t test or Mann-Whitney test. Differences between more than two groups were analyzed by one or two-way analysis of variance (ANOVA) or the non-parametric Kruskal Wallis test. Chi square test was used to compare the percentage of tumor-carrying mice by contingency tables. P values of 0.05 or less were considered to be statistically significant. All statistical analyses were performed with GraphPad Prism7 Software Inc.

## ACKNOWLEDGEMENTS

We thank Dr. J. Webster for comments and proofreading of the manuscript, all members from the Integrated Cellular Responses Laboratory for feedback on the manuscript. This study was funded by the National Agency for Scientific and Technological Promotion and National University of Quilmes to N.P. and the Fralin Biomedical Research Institute to C.V.F.

## AUTHOR CONTRIBUTIONS

I.A. designed and conducted experiments for Figures 1-8, performed statistical analyses for all panels, and contributed to the intellectual development of the study. G.H. and C.A.S. designed and conducted experiments for Figure 6 and Figure 7A-7F, and collaborated in tissue sample collection for Figures 3 and 5-8. A.C. conducted the bioinformatics analysis for Figures 7G-7L and Figure 9. N.P., D.G., and C.V.F. contributed to experimental design and intellectual development of the study. C.V.F. and I.A. prepared the initial manuscript draft, and C.V.F. wrote the final manuscript with input from all authors. C.V.F. provided resources, secured funding, and administered the project. All authors reviewed and approved the final manuscript.

## DECLARATION OF GENERATIVE AI AND AI-ASSISTED TECHNOLOGIES IN THE MANUSCRIPT PREPARATION PROCESS

During the preparation of this work the author(s) used Claude in order to develop code for analysis and proofreading. After using this tool/service, the author(s) reviewed and edited the content as needed and take(s) full responsibility for the content of the published article.

## SUPPLEMENTARY MATERIAL

### SUPPLEMENTARY TABLES

**Supplementary Table 1. Statistical analysis of circadian clock inhibitor effects on MLg cell migration.** Post-hoc comparisons from two-way ANOVA testing the effects of clock pathway inhibitors GSK2945 (REV-ERB antagonist), KL001 (CRY stabilizer), or PF670462 (CK1δ/ε inhibitor) on synchronized versus unsynchronized MLg lung fibroblast migration, with and without TGF-β treatment. Bolded comparisons are statistically significant. These pharmacological interventions validate the role of specific clock components in temporal migration gating characterized in Figure 1.

**Supplementary Table 2. Statistical analysis of genetic circadian disruption effects on cell migration.** Post-hoc comparisons from two-way ANOVA examining migration responses in MLg cells with BMAL1 knockdown (sh*Bmal1*) and NIH fibroblasts from PER2-knockout mice (PER2KO), tested under synchronized versus unsynchronized conditions with and without TGF-β stimulation. Bolded comparisons are statistically significant. These genetic approaches complement the pharmacological studies in Supplemental Table 1, providing orthogonal validation that core clock component disruption eliminates temporal migration gating and enhances TGF-β responsiveness as demonstrated in Figure 1.

**Supplementary Table 3. MetaCycle analysis of clock and MMP gene rhythmicity in MLg cells.** Circadian rhythm parameters (p-value, period, phase, mesor, amplitude) for *Bmal1*, *Mmp2*, *Mmp3*, and *Mmp9* expression in synchronized MLg lung fibroblasts under control conditions, TGF-β treatment, or GSK2945 (REV-ERB antagonist) treatment. This analysis quantifies the temporal oscillation patterns that underlie the migration phenotypes characterized in Figure 1 and MMP temporal dysregulation shown in Figure 2.

**Supplementary Table 4. MetaCycle analysis of core clock gene oscillations in lung tissue.** Circadian rhythm parameters (p-value, period, phase, mesor, amplitude) for *Bmal1*, *Cry1*, and *Cry2* expression in non-tumor-bearing and metastasis-bearing lung tissue from mice maintained under LD 12:12 or chronic jet lag (CJL) conditions. This analysis quantifies the tissue-level clock disruption that provides the foundation for pathway reorganization characterized in the main figures.

**Supplementary Table 5. MetaCycle analysis of pathway component rhythmicity in control and metastatic lungs.** Circadian rhythm parameters for inflammatory (*Tnf-α*, *Ccl2*), matrix remodeling (*Mmp2*, *Mmp3*, *Mmp9*), mechanotransduction (*Mob1a*, *Tead4*), TGF-β signaling (*Smad4*, *Smad7*), and EMT (*Snail*, *Zeb1*, *Fibronectin*) genes in lung tissue from non-tumor-bearing (control) and metastasis-bearing mice under LD or CJL conditions. This analysis provides the temporal dynamics underlying the pathway reorganization and convergence characterized throughout Figures 3-8.

**Supplementary Table 6. Statistical analysis of YAP inhibitor temporal effects on cell migration.** Post-hoc comparisons from two-way ANOVA examining the effects of verteporfin (YAP/TEAD inhibitor) treatment at different time points (0h, 6h, 12h) on synchronized versus unsynchronized MLg cell migration, with and without TGF-β stimulation and dexamethasone treatment. Bolded comparisons are statistically significant. These time-course studies validate the temporal dependence of YAP/TEAD pathway activity in migration responses, providing mechanistic support for the obligate YAP requirement characterized in Figure 5 and the temporal YAP reorganization demonstrated in Figures 5 AND 8.

**Supplementary Table 7. Primer sequences for quantitative real-time PCR analysis.** Forward and reverse primer sequences with predicted product sizes (bp) for all genes analyzed by RT-qPCR throughout this study, including circadian clock components (*Bmal1*, *Cry1*, *Cry2*), matrix metalloproteases (*Mmp2*, *Mmp3*, *Mmp9*), inflammatory mediators (*Tnf-α*, *Ccl2*), growth factors (*Tgf-β*), mechanotransduction components (*Mob1a*, *Tead4*), TGF-β signaling elements (*Smad4*, *Smad7*), and EMT markers (*Zeb1*, *Snail*, *Fibronectin*). *Hprt* serves as the housekeeping gene for normalization.

### SUPPLEMENTARY FIGURES

**Figure S1. Effective knockdown of BMAL1 validates circadian clock disruption in MLg lung fibroblasts.** (**A**) Relative mRNA levels of *Bmal1* quantified by RT-qPCR in MLg cells transfected with control shRNA (shCtrl, black) or BMAL1-targeting shRNA (sh*Bmal1*, gray). (**B**) Representative Western blot (top) and densitometric quantification (bottom) of BMAL1 protein levels in transfected MLg cells. GAPDH serves as loading control. Knockdown efficiency demonstrates complete disruption of the core circadian transcriptional machinery required for temporal migration gating (Figure 1). Data represent mean ± SD from n=3 biological replicates per condition. Unpaired t-test: (**A**) ***p=0.0002, (**B**) ****p<0.0001.

**Figure S2. Growth factors and cytokines differentially regulate Bmal1 and MMP expression in synchronized MLg cells.** Relative mRNA levels of *Bmal1* (**A**), *Mmp2* (**B**), *Mmp3* (**C**), and *Mmp9* (**D**) quantified by RT-qPCR in synchronized MLg cells treated with EGF (diagonal lines), FGF2 (horizontal lines), TGF-β (black squares), or TNF-α (crosshatch) at 12 and 21 h post-synchronization (dark and light pink, respectively) compared to vehicle controls (solid). Note that TNF-α dramatically elevates *Bmal1* and all three MMPs, particularly *Mmp3* at 21h. TGF-β strongly induces *Bmal1* and *Mmp9*, while EGF and FGF2 selectively enhance *Mmp9* without affecting *Mmp2* or *Mmp3*. Data represent mean ± SD from n=3 biological replicates per condition. Two-way ANOVA: (**A**) p=0.0149, (**B**) p=0.010, (**C**) p=0.0432, (**D**) p=0.014. Post-hoc comparisons: *p<0.05, **p<0.01, ***p<0.001, ****p<0.0001. Note: TGF-β data shown here were obtained from an independent experiment performed concurrently with EGF, FGF2, and TNF-α treatments to enable direct comparison with the same control; TGF-β data in Figure 2 represent a separate replicate set.

**Figure S3. Chronic jet lag disrupts core circadian clock gene oscillation in lung tissue.** Relative mRNA levels of core clock components *Bmal1* (**A**), *Cry1* (**B**), and *Cry2* (**C**) quantified by RT-qPCR in non-tumor-bearing lung tissue from mice maintained under LD 12:12 (pink) or CJL (cyan) conditions and harvested at eight zeitgeber times. Fitted curves represent statistically significant circadian rhythmicity determined by MetaCycle analysis (Supplemental Table 4). Note disrupted oscillation patterns under CJL conditions, validating circadian clock disruption in the tissue microenvironment that underlies the pathway reorganization characterized in the main figures. Data represent mean ± SEM from n=4 animals per timepoint. MetaCycle analysis: (**A**) p=0.0149, (**B**) p=0.010, (**C**) p=0.0432.

**Figure S4. Metastatic burden further impairs circadian clock gene oscillation in lung tissue.** Relative mRNA levels of core clock components *Bmal1* (**A**), *Cry1* (**B**), and *Cry2* (**C**) quantified by RT-qPCR in metastasis-bearing lung tissue from mice maintained under LD 12:12 (pink) or CJL (cyan) conditions and harvested 21 days post-B16F10 inoculation at eight zeitgeber times. Fitted curve in (**C**) represents statistically significant circadian rhythmicity for *Cry2* under CJL conditions determined by MetaCycle analysis (Supplemental Table 4). Note severely dampened oscillations compared to non-tumor-bearing lungs (Figure S3), demonstrating that metastatic burden exacerbates circadian clock disruption in the tissue microenvironment. This provides the molecular foundation for the amplified pathway reorganization characterized in metastatic lungs (Figures 7-8). Data represent mean ± SEM from n=4 animals per timepoint. MetaCycle analysis: (**C**) p=0.014.

**Figure S5. Total macrophage populations remain stable across circadian and metastatic conditions.** Flow cytometric quantification of total macrophage percentages (F4/80+) in control non-tumor-bearing (**A-B**) and metastasis-bearing (**C-D**) lung tissue from mice maintained under LD 12:12 (**A**, **C**) or CJL (**B**, **D**) conditions and harvested at ZT3, 9, 15, and 21. Note stable total macrophage percentages across all conditions and timepoints, validating that the M1/M2 population changes and temporal reorganization characterized in Figure 7 reflect functional polarization shifts rather than alterations in total macrophage recruitment or retention. Data represent mean ± SEM from n=4-6 animals per condition and timepoint.

## REFERENCES

1. Chen, Z., Yoo, S.H., and Takahashi, J.S. (2018). Development and Therapeutic Potential of Small-Molecule Modulators of Circadian Systems. Annu Rev Pharmacol Toxicol 58, 231–252. 10.1146/annurev-pharmtox-010617-052645.

2. Golombek, D.A., Casiraghi, L.P., Agostino, P.V., Paladino, N., Duhart, J.M., Plano, S.A., and Chiesa, J.J. (2013). The times they’re a-changing: effects of circadian desynchronization on physiology and disease. J Physiol Paris 107, 310–322. 10.1016/j.jphysparis.2013.03.007.

3. Zhang, Z., Hunter, L., Wu, G., Maidstone, R., Mizoro, Y., Vonslow, R., Fife, M., Hopwood, T., Begley, N., Saer, B., et al. (2019). Genome-wide effect of pulmonary airway epithelial cell-specific Bmal1 deletion. Faseb J 33, 6226–6238. 10.1096/fj.201801682R.

4. Pezuk, P., Mohawk, J.A., Wang, L.A., and Menaker, M. (2012). Glucocorticoids as entraining signals for peripheral circadian oscillators. Endocrinology 153, 4775–4783. 10.1210/en.2012-1486.

5. Buhr, E.D., Yoo, S.H., and Takahashi, J.S. (2010). Temperature as a universal reseêng cue for mammalian circadian oscillators. Science 330, 379–385. 10.1126/science.1195262.

6. Abraham, U., Granada, A.E., Westermark, P.O., Heine, M., Kramer, A., and Herzel, H. (2010). Coupling governs entrainment range of circadian clocks. Mol Syst Biol 6, 438. 10.1038/msb.2010.92.

7. Pekovic-Vaughan, V., Gibbs, J., Yoshitane, H., Yang, N., Pathiranage, D., Guo, B., Sagami, A., Taguchi, K., Bechtold, D., Loudon, A., et al. (2014). The circadian clock regulates rhythmic activation of the NRF2/glutathione-mediated antioxidant defense pathway to modulate pulmonary fibrosis. Genes & development 28, 548–560. 10.1101/gad.237081.113.

8. Hadden, H., Soldin, S.J., and Massaro, D. (2012). Circadian disruption alters mouse lung clock gene expression and lung mechanics. J Appl Physiol (1985) 113, 385–392. 10.1152/japplphysiol.00244.2012.

9. Ehlers, A., Xie, W., Agapov, E., Brown, S., Steinberg, D., Tidwell, R., Sajol, G., Schutz, R., Weaver, R., Yu, H., et al. (2018). BMAL1 links the circadian clock to viral airway pathology and asthma phenotypes. Mucosal Immunol 11, 97–111. 10.1038/mi.2017.24.

10. Pariollaud, M., Gibbs, J.E., Hopwood, T.W., Brown, S., Begley, N., Vonslow, R., Poolman, T., Guo, B., Saer, B., Jones, D.H., et al. (2018). Circadian clock component REV-ERBalpha controls homeostatic regulation of pulmonary inflammation. J Clin Invest 128, 2281–2296. 10.1172/JCI93910.

11. Sengupta, S., Tang, S.Y., Devine, J.C., Anderson, S.T., Nayak, S., Zhang, S.L., Valenzuela, A., Fisher, D.G., Grant, G.R., Lopez, C.B., and FitzGerald, G.A. (2019). Circadian control of lung inflammation in influenza infection. Nature communications 10, 4107. 10.1038/s41467-019-11400-9.

12. Thenappan, T., Chan, S.Y., and Weir, E.K. (2018). Role of extracellular matrix in the pathogenesis of pulmonary arterial hypertension. Am J Physiol Heart Circ Physiol 315, H1322–H1331. 10.1152/ajpheart.00136.2018.

13. Karakioulaki, M., Papakonstantinou, E., and Stolz, D. (2020). Extracellular matrix remodelling in COPD. Eur Respir Rev 29. 10.1183/16000617.0124-2019.

14. Robert, S., Gicquel, T., Victoni, T., Valenca, S., Barreto, E., Bailly-Maitre, B., Boichot, E., and Lagente, V. (2016). Involvement of matrix metalloproteinases (MMPs) and inflammasome pathway in molecular mechanisms of fibrosis. Biosci Rep 36. 10.1042/BSR20160107.

15. Pain, M., Bermudez, O., Lacoste, P., Royer, P.J., Boíuri, K., Tissot, A., Brouard, S., Eickelberg, O., and Magnan, A. (2014). Tissue remodelling in chronic bronchial diseases: from the epithelial to mesenchymal phenotype. Eur Respir Rev 23, 118–130. 10.1183/09059180.00004413.

16. Bornes, L., Belthier, G., and van Rheenen, J. (2021). Epithelial-to-Mesenchymal Transition in the Light of Plasticity and Hybrid E/M States. J Clin Med 10. 10.3390/jcm10112403.

17. Martinez-Espinosa, I., Serrato, J.A., Cabello-Gutierrez, C., Carlos-Reyes, A., and Ortiz-Quintero, B. (2024). Mechanisms of microRNA Regulation of the Epithelial-Mesenchymal Transition (EMT) in Lung Cancer. Life (Basel) 14. 10.3390/life14111431.

18. Thiery, J.P., Acloque, H., Huang, R.Y., and Nieto, M.A. (2009). Epithelial-mesenchymal transitions in development and disease. Cell 139, 871–890. 10.1016/j.cell.2009.11.007.

19. Thomson, S., Peê, F., Sujka-Kwok, I., Mercado, P., Bean, J., Monaghan, M., Seymour, S.L., Argast, G.M., Epstein, D.M., and Haley, J.D. (2011). A systems view of epithelial-mesenchymal transition signaling states. Clin Exp Metastasis 28, 137–155. 10.1007/s10585-010-9367-3.

20. Jordan, N.V., Johnson, G.L., and Abell, A.N. (2011). Tracking the intermediate stages of epithelial-mesenchymal transition in epithelial stem cells and cancer. Cell cycle 10, 2865–2873. 10.4161/cc.10.17.17188.

21. Nieto, M.A., Huang, R.Y., Jackson, R.A., and Thiery, J.P. (2016). Emt: 2016. Cell 166, 21–45. 10.1016/j.cell.2016.06.028.

22. Derynck, R., Muthusamy, B.P., and Saeteurn, K.Y. (2014). Signaling pathway cooperation in TGF-beta-induced epithelial-mesenchymal transition. Curr Opin Cell Biol 31, 56–66. 10.1016/j.ceb.2014.09.001.

23. Zhong, Z., Jiao, Z., and Yu, F.X. (2024). The Hippo signaling pathway in development and regeneration. Cell Rep 43, 113926. 10.1016/j.celrep.2024.113926.

24. Ma, S., Meng, Z., Chen, R., and Guan, K.L. (2019). The Hippo Pathway: Biology and Pathophysiology. Annu Rev Biochem 88, 577–604. 10.1146/annurev-biochem-013118-111829.

25. Praskova, M., Xia, F., and Avruch, J. (2008). MOBKL1A/MOBKL1B phosphorylation by MST1 and MST2 inhibits cell proliferation. Curr Biol 18, 311–321. 10.1016/j.cub.2008.02.006.

26. Hergovich, A., Schmitz, D., and Hemmings, B.A. (2006). The human tumour suppressor LATS1 is activated by human MOB1 at the membrane. Biochem Biophys Res Commun 345, 50–58. 10.1016/j.bbrc.2006.03.244.

27. Liu, C.Y., Zha, Z.Y., Zhou, X., Zhang, H., Huang, W., Zhao, D., Li, T., Chan, S.W., Lim, C.J., Hong, W., et al. (2010). The hippo tumor pathway promotes TAZ degradation by phosphorylating a phosphodegron and recruiting the SCFbeta-TrCP E3 ligase. The Journal of biological chemistry 285, 37159–37169. 10.1074/jbc.M110.152942.

28. Zhao, B., Ye, X., Yu, J., Li, L., Li, W., Li, S., Yu, J., Lin, J.D., Wang, C.Y., Chinnaiyan, A.M., et al. (2008). TEAD mediates YAP-dependent gene induction and growth control. Genes & development 22, 1962–1971. 10.1101/gad.1664408.

29. Zhang, Y.E. (2009). Non-Smad pathways in TGF-beta signaling. Cell Res 19, 128–139. 10.1038/cr.2008.328.

30. Lee, K.P., Lee, J.H., Kim, T.S., Kim, T.H., Park, H.D., Byun, J.S., Kim, M.C., Jeong, W.I., Calvisi, D.F., Kim, J.M., and Lim, D.S. (2010). The Hippo-Salvador pathway restrains hepatic oval cell proliferation, liver size, and liver tumorigenesis. Proceedings of the National Academy of Sciences of the United States of America 107, 8248–8253. 10.1073/pnas.0912203107.

31. Lu, L., Li, Y., Kim, S.M., Bossuyt, W., Liu, P., Qiu, Q., Wang, Y., Halder, G., Finegold, M.J., Lee, J.S., and Johnson, R.L. (2010). Hippo signaling is a potent in vivo growth and tumor suppressor pathway in the mammalian liver. Proceedings of the National Academy of Sciences of the United States of America 107, 1437–1442. 10.1073/pnas.0911427107.

32. Kawabata, M., Inoue, H., Hanyu, A., Imamura, T., and Miyazono, K. (1998). Smad proteins exist as monomers in vivo and undergo homo- and hetero-oligomerization upon activation by serine/threonine kinase receptors. The EMBO journal 17, 4056–4065. 10.1093/emboj/17.14.4056.

33. Tzavlaki, K., and Moustakas, A. (2020). TGF-beta Signaling. Biomolecules 10. 10.3390/biom10030487.

34. Yan, X., Liao, H., Cheng, M., Shi, X., Lin, X., Feng, X.H., and Chen, Y.G. (2016). Smad7 Protein Interacts with Receptor-regulated Smads (R-Smads) to Inhibit Transforming Growth Factor-beta (TGF-beta)/Smad Signaling. The Journal of biological chemistry 291, 382–392. 10.1074/jbc.M115.694281.

35. Massague, J. (2008). TGFbeta in Cancer. Cell 134, 215–230. 10.1016/j.cell.2008.07.001.

36. Shi, Y., and Massague, J. (2003). Mechanisms of TGF-beta signaling from cell membrane to the nucleus. Cell 113, 685–700. 10.1016/s0092-8674(03)00432-x.

37. Varelas, X., and Wrana, J.L. (2012). Coordinating developmental signaling: novel roles for the Hippo pathway. Trends Cell Biol 22, 88–96. 10.1016/j.tcb.2011.10.002.

38. Page-McCaw, A., Ewald, A.J., and Werb, Z. (2007). Matrix metalloproteinases and the regulation of tissue remodelling. Nat Rev Mol Cell Biol 8, 221–233. 10.1038/nrm2125.

39. Egeblad, M., and Werb, Z. (2002). New functions for the matrix metalloproteinases in cancer progression. Nat Rev Cancer 2, 161–174. 10.1038/nrc745.

40. Li, Y., Liu, F., Cai, Q., Deng, L., Ouyang, Q., Zhang, X.H., and Zheng, J. (2025). Invasion and metastasis in cancer: molecular insights and therapeutic targets. Signal Transduct Target Ther 10, 57. 10.1038/s41392-025-02148-4.

41. Xiao, G., Wang, X., Xu, Z., Liu, Y., and Jing, J. (2025). Lung-specific metastasis: the coevolution of tumor cells and lung microenvironment. Mol Cancer 24, 118. 10.1186/s12943-025-02318-6.

42. Papagiannakopoulos, T., Bauer, M.R., Davidson, S.M., Heimann, M., Subbaraj, L., Bhutkar, A., Bartlebaugh, J., Vander Heiden, M.G., and Jacks, T. (2016). Circadian Rhythm Disruption Promotes Lung Tumorigenesis. Cell Metab 24, 324–331. 10.1016/j.cmet.2016.07.001.

43. Aiello, I., Fedele, M.L.M., Roman, F., Marpegan, L., Caldart, C., Chiesa, J.J., Golombek, D.A., Finkielstein, C.V., and Paladino, N. (2020). Circadian disruption promotes tumor-immune microenvironment remodeling favoring tumor cell proliferation. Sci Adv 6. 10.1126/sciadv.aaz4530.

44. Hadadi, E., Taylor, W., Li, X.M., Aslan, Y., Villote, M., Riviere, J., Duvallet, G., Auriau, C., Dulong, S., Raymond-Letron, I., et al. (2020). Chronic circadian disruption modulates breast cancer stemness and immune microenvironment to drive metastasis in mice. Nature communications 11, 3193. 10.1038/s41467-020-16890-6.

45. Houshyari, M., and Taghizadeh-Hesary, F. (2023). The Metastatic Spread of Breast Cancer Accelerates during Sleep: How the Study Design can Affect the Results. Asian Pac J Cancer Prev 24, 353–355. 10.31557/APJCP.2023.24.2.353.

46. Chun, S.K., Fortin, B.M., Fellows, R.C., Habowski, A.N., Verlande, A., Song, W.A., Mahieu, A.L., Lefebvre, A., Sterrenberg, J.N., Velez, L.M., et al. (2022). Disruption of the circadian clock drives Apc loss of heterozygosity to accelerate colorectal cancer. Sci Adv 8, eabo2389. 10.1126/sciadv.abo2389.

47. Wang, C., Barnoud, C., Cenerenti, M., Sun, M., Caffa, I., Kizil, B., Bill, R., Liu, Y., Pick, R., Garnier, L., et al. (2023). Dendritic cells direct circadian anti-tumour immune responses. Nature 614, 136–143. 10.1038/s41586-022-05605-0.

48. Pariollaud, M., Ibrahim, L.H., Irizarry, E., Mello, R.M., Chan, A.B., Altman, B.J., Shaw, R.J., Bollong, M.J., Wiseman, R.L., and Lamia, K.A. (2022). Circadian disruption enhances HSF1 signaling and tumorigenesis in Kras-driven lung cancer. Sci Adv 8, eabo1123. 10.1126/sciadv.abo1123.

49. Hoyle, N.P., Seinkmane, E., Putker, M., Feeney, K.A., Krogager, T.P., Chesham, J.E., Bray, L.K., Thomas, J.M., Dunn, K., Blaikley, J., and O’Neill, J.S. (2017). Circadian actin dynamics drive rhythmic fibroblast mobilization during wound healing. Sci Transl Med 9. 10.1126/scitranslmed.aal2774.

50. Dong, C., Gongora, R., Sosulski, M.L., Luo, F., and Sanchez, C.G. (2016). Regulation of transforming growth factor-beta1 (TGF-beta1)-induced pro-fibrotic activities by circadian clock gene BMAL1. Respir Res 17, 4. 10.1186/s12931-016-0320-0.

51. Brown, S.A. (2014). Circadian clock-mediated control of stem cell division and differentiation: beyond night and day. Development 141, 3105–3111. 10.1242/dev.104851.

52. He, W., Holtkamp, S., Hergenhan, S.M., Kraus, K., de Juan, A., Weber, J., Bradfield, P., Grenier, J.M.P., Pelletier, J., Druzd, D., et al. (2018). Circadian Expression of Migratory Factors Establishes Lineage-Specific Signatures that Guide the Homing of Leukocyte Subsets to Tissues. Immunity 49, 1175–1190 e1177. 10.1016/j.immuni.2018.10.007.

53. Ren, D.L., Li, Y.J., Hu, B.B., Wang, H., and Hu, B. (2015). Melatonin regulates the rhythmic migration of neutrophils in live zebrafish. J Pineal Res 58, 452–460. 10.1111/jpi.12230.

54. Lin, H.H., Robertson, K.L., Bisbee, H.A., and Farkas, M.E. (2020). Oncogenic and Circadian Effects of Small Molecules Directly and Indirectly Targeting the Core Circadian Clock. Integr Cancer Ther 19, 1534735420924094. 10.1177/1534735420924094.

55. Xiong, G., and Xu, R. (2022). Retinoid orphan nuclear receptor alpha (RORalpha) suppresses the epithelial-mesenchymal transition (EMT) by directly repressing Snail transcription. The Journal of biological chemistry 298, 102059. 10.1016/j.jbc.2022.102059.

56. Zhang, T., Zhao, M., Lu, D., Wang, S., Yu, F., Guo, L., Wen, S., and Wu, B. (2018). REV-ERBalpha Regulates CYP7A1 Through Repression of Liver Receptor Homolog-1. Drug Metab Dispos 46, 248–258. 10.1124/dmd.117.078105.

57. Hirota, T., Lee, J.W., St John, P.C., Sawa, M., Iwaisako, K., Noguchi, T., Pongsawakul, P.Y., Sonntag, T., Welsh, D.K., Brenner, D.A., et al. (2012). Identification of small molecule activators of cryptochrome. Science 337, 1094–1097. 10.1126/science.1223710.

58. Meng, Q.J., Maywood, E.S., Bechtold, D.A., Lu, W.Q., Li, J., Gibbs, J.E., Dupre, S.M., Chesham, J.E., Rajamohan, F., Knafels, J., et al. (2010). Entrainment of disrupted circadian behavior through inhibition of casein kinase 1 (CK1) enzymes. Proceedings of the National Academy of Sciences of the United States of America 107, 15240–15245. 10.1073/pnas.1005101107.

59. Liu, A.C., Tran, H.G., Zhang, E.E., Priest, A.A., Welsh, D.K., and Kay, S.A. (2008). Redundant function of REV-ERBalpha and beta and non-essential role for Bmal1 cycling in transcriptional regulation of intracellular circadian rhythms. PLoS Genet 4, e1000023. 10.1371/journal.pgen.1000023.

60. Yang, N., Smyllie, N.J., Morris, H., Goncalves, C.F., Dudek, M., Pathiranage, D.R.J., Chesham, J.E., Adamson, A., Spiller, D.G., Zindy, E., et al. (2020). Quantitative live imaging of Venus::BMAL1 in a mouse model reveals complex dynamics of the master circadian clock regulator. PLoS Genet 16, e1008729. 10.1371/journal.pgen.1008729.

61. Casiraghi, L.P., Oda, G.A., Chiesa, J.J., Friesen, W.O., and Golombek, D.A. (2012). Forced desynchronization of activity rhythms in a model of chronic jet lag in mice. Journal of biological rhythms 27, 59–69. 10.1177/0748730411429447.

62. Palombo, P., Moreno-Villanueva, M., and Mangerich, A. (2015). Day and night variations in the repair of ionizing-radiation-induced DNA damage in mouse splenocytes. DNA Repair (Amst) 28, 37–47. 10.1016/j.dnarep.2015.02.002.

63. Kang, T.H., Reardon, J.T., Kemp, M., and Sancar, A. (2009). Circadian oscillation of nucleotide excision repair in mammalian brain. Proceedings of the National Academy of Sciences of the United States of America 106, 2864–2867. 10.1073/pnas.0812638106.

64. Ryzhikov, M., Ehlers, A., Steinberg, D., Xie, W., Oberlander, E., Brown, S., Gilmore, P.E., Townsend, R.R., Lane, W.S., Dolinay, T., et al. (2019). Diurnal Rhythms Spatially and Temporally Organize Autophagy. Cell Rep 26, 1880–1892 e1886. 10.1016/j.celrep.2019.01.072.

65. Crespo, M.T., Trebucq, L.L., Senna, C.A., Hokama, G., Paladino, N., Agostino, P.V., and Chiesa, J.J. (2025). Circadian disruption of feeding-fasting rhythm and its consequences for metabolic, immune, cancer, and cognitive processes. Biomed J 48, 100827. 10.1016/j.bj.2025.100827.

66. Jerigova, V., Zeman, M., and Okuliarova, M. (2022). Circadian Disruption and Consequences on Innate Immunity and Inflammatory Response. Int J Mol Sci 23. 10.3390/ijms232213722.

67. Liu-Chiíenden, Y., Huang, B., Shim, J.S., Chen, Q., Lee, S.J., Anders, R.A., Liu, J.O., and Pan, D. (2012). Genetic and pharmacological disruption of the TEAD-YAP complex suppresses the oncogenic activity of YAP. Genes & development 26, 1300–1305. 10.1101/gad.192856.112.

68. Moustakas, A., and Heldin, C.H. (2016). Mechanisms of TGFbeta-Induced Epithelial-Mesenchymal Transition. J Clin Med 5. 10.3390/jcm5070063.

69. Lamouille, S., Xu, J., and Derynck, R. (2014). Molecular mechanisms of epithelial-mesenchymal transition. Nat Rev Mol Cell Biol 15, 178–196. 10.1038/nrm3758.

70. Jayasingam, S.D., Citartan, M., Thang, T.H., Mat Zin, A.A., Ang, K.C., and Ch’ng, E.S. (2019). Evaluating the Polarization of Tumor-Associated Macrophages Into M1 and M2 Phenotypes in Human Cancer Tissue: Technicalities and Challenges in Routine Clinical Practice. Front Oncol 9, 1512. 10.3389/fonc.2019.01512.

71. Wang, C., Zeng, Q., Gul, Z.M., Wang, S., Pick, R., Cheng, P., Bill, R., Wu, Y., Naulaerts, S., Barnoud, C., et al. (2024). Circadian tumor infiltration and function of CD8(+) T cells dictate immunotherapy efficacy. Cell 187, 2690–2702 e2617. 10.1016/j.cell.2024.04.015.

72. Strizova, Z., Benesova, I., Bartolini, R., Novysedlak, R., Cecrdlova, E., Foley, L.K., and Striz, I. (2023). M1/M2 macrophages and their overlaps - myth or reality? Clin Sci (Lond) 137, 1067–1093. 10.1042/CS20220531.

73. Wu, G., Ruben, M.D., Schmidt, R.E., Francey, L.J., Smith, D.F., Anafi, R.C., Hughey, J.J., Tasseff, R., Sherrill, J.D., Oblong, J.E., et al. (2018). Population-level rhythms in human skin with implications for circadian medicine. Proceedings of the National Academy of Sciences of the United States of America 115, 12313–12318. 10.1073/pnas.1809442115.

74. Shilts, J., Chen, G., and Hughey, J.J. (2018). Evidence for widespread dysregulation of circadian clock progression in human cancer. PeerJ 6, e4327. 10.7717/peerj.4327.

75. Aiello, I., Mul Fedele, M.L., Roman, F., Marpegan, L., Caldart, C., Chiesa, J.J., Golombek, D.A., Finkielstein, C.V., and Paladino, N. (2020). Circadian disruption promotes tumor-immune microenvironment remodeling favoring tumor cell proliferation. Sci Adv 6. 10.1126/sciadv.aaz4530.

76. Lee, Y. (2021). Roles of circadian clocks in cancer pathogenesis and treatment. Exp Mol Med 53, 1529–1538. 10.1038/s12276-021-00681-0.

77. Diamantopoulou, Z., Castro-Giner, F., Schwab, F.D., Foerster, C., Saini, M., Budinjas, S., Striímaíer, K., Krol, I., Seifert, B., Heinzelmann-Schwarz, V., et al. (2022). The metastatic spread of breast cancer accelerates during sleep. Nature 607, 156–162. 10.1038/s41586-022-04875-y.

78. Wu, J., Jing, X., Du, Q., Sun, X., Holgersson, K., Gao, J., He, X., Hosaka, K., Zhao, C., Tao, W., et al. (2023). Disruption of the Clock Component Bmal1 in Mice Promotes Cancer Metastasis through the PAI-1-TGF-beta-myoCAF-Dependent Mechanism. Adv Sci (Weinh) 10, e2301505. 10.1002/advs.202301505.

79. Finger, A.M., Jaschke, S., Del Olmo, M., Hurwitz, R., Granada, A.E., Herzel, H., and Kramer, A. (2021). Intercellular coupling between peripheral circadian oscillators by TGF-beta signaling. Sci Adv 7. 10.1126/sciadv.abg5174.

80. Li, S.Y., Hammarlund, J.A., Wu, G., Lian, J.W., Howell, S.J., Clarke, R.B., Adamson, A.D., Goncalves, C.F., Hogenesch, J.B., Anafi, R.C., and Meng, Q.J. (2024). Tumor circadian clock strength influences metastatic potential and predicts patient prognosis in luminal A breast cancer. Proceedings of the National Academy of Sciences of the United States of America 121, e2311854121. 10.1073/pnas.2311854121.

81. Wang, Y., Narasimamurthy, R., Qu, M., Shi, N., Guo, H., Xue, Y., and Barker, N. (2024). Circadian regulation of cancer stem cells and the tumor microenvironment during metastasis. Nat Cancer 5, 546–556. 10.1038/s43018-024-00759-4.

82. Zhou, Z., Zhang, R., Zhang, Y., Xu, Y., Wang, R., Chen, S., Lv, Y., Chen, Y., Ren, Y., Luo, P., et al. (2024). Circadian disruption in cancer hallmarks: Novel insight into the molecular mechanisms of tumorigenesis and cancer treatment. Cancer Leí 604, 217273. 10.1016/j.canlet.2024.217273.

83. Zhu, Y., Zheng, Y., Dai, R., and Gu, X. (2024). Crosstalk between Circadian Rhythm Dysregulation and Tumorigenesis, Tumor Metabolism and Tumor Immune Response. Aging Dis 16, 2073–2099. 10.14336/AD.2024.0533.

84. Blask, D.E., Hill, S.M., Dauchy, R.T., Xiang, S., Yuan, L., Duplessis, T., Mao, L., Dauchy, E., and Sauer, L.A. (2011). Circadian regulation of molecular, dietary, and metabolic signaling mechanisms of human breast cancer growth by the nocturnal melatonin signal and the consequences of its disruption by light at night. J Pineal Res 51, 259–269. 10.1111/j.1600-079X.2011.00888.x.

85. Hill, S.M., Blask, D.E., Xiang, S., Yuan, L., Mao, L., Dauchy, R.T., Dauchy, E.M., Frasch, T., and Duplesis, T. (2011). Melatonin and associated signaling pathways that control normal breast epithelium and breast cancer. J Mammary Gland Biol Neoplasia 16, 235–245. 10.1007/s10911-011-9222-4.

86. Wang, J., Li, S., Li, X., Li, B., Li, Y., Xia, K., Yang, Y., Aman, S., Wang, M., and Wu, H. (2019). Circadian protein BMAL1 promotes breast cancer cell invasion and metastasis by up-regulating matrix metalloproteinase9 expression. Cancer Cell Int 19, 182. 10.1186/s12935-019-0902-2.

87. Zhao, B., Nepovimova, E., and Wu, Q. (2025). The role of circadian rhythm regulator PERs in oxidative stress, immunity, and cancer development. Cell Commun Signal 23, 30. 10.1186/s12964-025-02040-2.

88. Zeng, Y., Guo, Z., Wu, M., Chen, F., and Chen, L. (2024). Circadian rhythm regulates the function of immune cells and participates in the development of tumors. Cell Death Discov 10, 199. 10.1038/s41420-024-01960-1.

89. Murgo, E., Falco, G., Serviddio, G., Mazzoccoli, G., and Colangelo, T. (2024). Circadian paíerns of growth factor receptor-dependent signaling and implications for carcinogenesis. Cell Commun Signal 22, 319. 10.1186/s12964-024-01676-w.

90. Huang, Z., Zeng, L., Ruan, Z., Zeng, Q., Yan, H., Jiang, W., Xiong, Y., Zhou, C., Yang, H., Liu, L., et al. (2026). Time-of-day immunochemotherapy in non-small cell lung cancer: a randomized phase 3 trial. Nat Med. 10.1038/s41591-025-04181-w.

91. Qian, D.C., Kleber, T., Brammer, B., Xu, K.M., Switchenko, J.M., Janopaul-Naylor, J.R., Zhong, J., Yushak, M.L., Harvey, R.D., Paulos, C.M., et al. (2021). Effect of immunotherapy time-of-day infusion on overall survival among patients with advanced melanoma in the USA (MEMOIR): a propensity score-matched analysis of a single-centre, longitudinal study. Lancet Oncol 22, 1777–1786. 10.1016/S1470-2045(21)00546-5.

92. Kim, B.G., Malek, E., Choi, S.H., Ignatz-Hoover, J.J., and Driscoll, J.J. (2021). Novel therapies emerging in oncology to target the TGF-beta pathway. J Hematol Oncol 14, 55. 10.1186/s13045-021-01053-x.

93. Pobbati, A.V., Kumar, R., Rubin, B.P., and Hong, W. (2023). Therapeutic targeting of TEAD transcription factors in cancer. Trends Biochem Sci 48, 450–462. 10.1016/j.tibs.2022.12.005.

94. Sulli, G., Manoogian, E.N.C., Taub, P.R., and Panda, S. (2018). Training the Circadian Clock, Clocking the Drugs, and Drugging the Clock to Prevent, Manage, and Treat Chronic Diseases. Trends Pharmacol Sci 39, 812–827. 10.1016/j.tips.2018.07.003.

95. Mitsuhashi, A., Koyama, K., Ogino, H., Matsuo, R., Thi Nguyen, N., Yabuki, Y., Ozaki, R., Tsukazaki, Y., Morita, Y., Yoshida, A., et al. (2025). Clock pathway inhibitor overcomes tumor immune-exclusion via regulation of fibrocyte differentiation. NPJ Precis Oncol 9, 274. 10.1038/s41698-025-01066-6.

96. Das, M., Ellies, L.G., Kumar, D., Sauceda, C., Oberg, A., Gross, E., Mandt, T., Newton, I.G., Kaur, M., Sears, D.D., and Webster, N.J.G. (2021). Time-restricted feeding normalizes hyperinsulinemia to inhibit breast cancer in obese postmenopausal mouse models. Nature communications 12, 565. 10.1038/s41467-020-20743-7.

97. Erren, T.C., Morfeld, P., Gross, J.V., Wild, U., and Lewis, P. (2019). IARC 2019: "Night shiñ work" is probably carcinogenic: What about disturbed chronobiology in all walks of life? J Occup Med Toxicol 14, 29. 10.1186/s12995-019-0249-6.

98. group, I.M.V. (2019). Carcinogenicity of night shiñ work. Lancet Oncol 20, 1058–1059. 10.1016/S1470-2045(19)30455-3.

99. Kisamore, C.O., Kisamore, C.A., and Walker, W.H., 2nd (2024). Circadian Rhythm Disruption in Cancer Survivors: From Oncogenesis to Quality of Life. Cancer Med 13, e70353. 10.1002/cam4.70353.

100. Wang, Y., Xu, X., Maglic, D., Dill, M.T., Mojumdar, K., Ng, P.K., Jeong, K.J., Tsang, Y.H., Moreno, D., Bhavana, V.H., et al. (2018). Comprehensive Molecular Characterization of the Hippo Signaling Pathway in Cancer. Cell Rep 25, 1304–1317 e1305. 10.1016/j.celrep.2018.10.001.

101. Wu, G., Anafi, R.C., Hughes, M.E., Kornacker, K., and Hogenesch, J.B. (2016). MetaCycle: an integrated R package to evaluate periodicity in large scale data. Bioinformatics 32, 3351–3353. 10.1093/bioinformatics/btw405.

