## Supplementary figures and tables for "Circadian Disruption Drives Extracellular Matrix Remodeling to Facilitate Pulmonary Metastatic Colonization"

Sup_Figure_1_Aiello et al.

A.

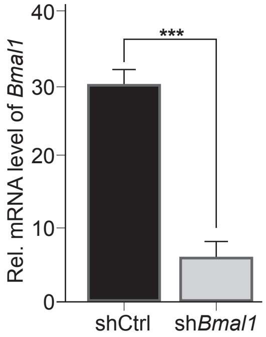


B.

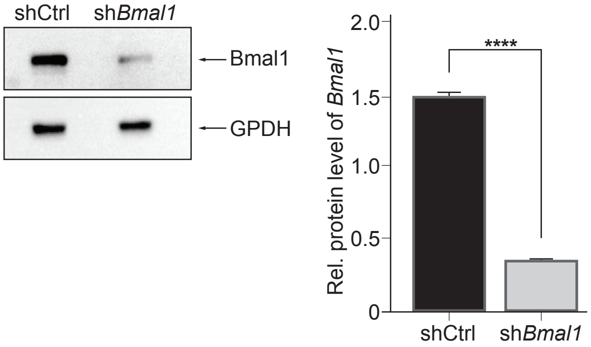

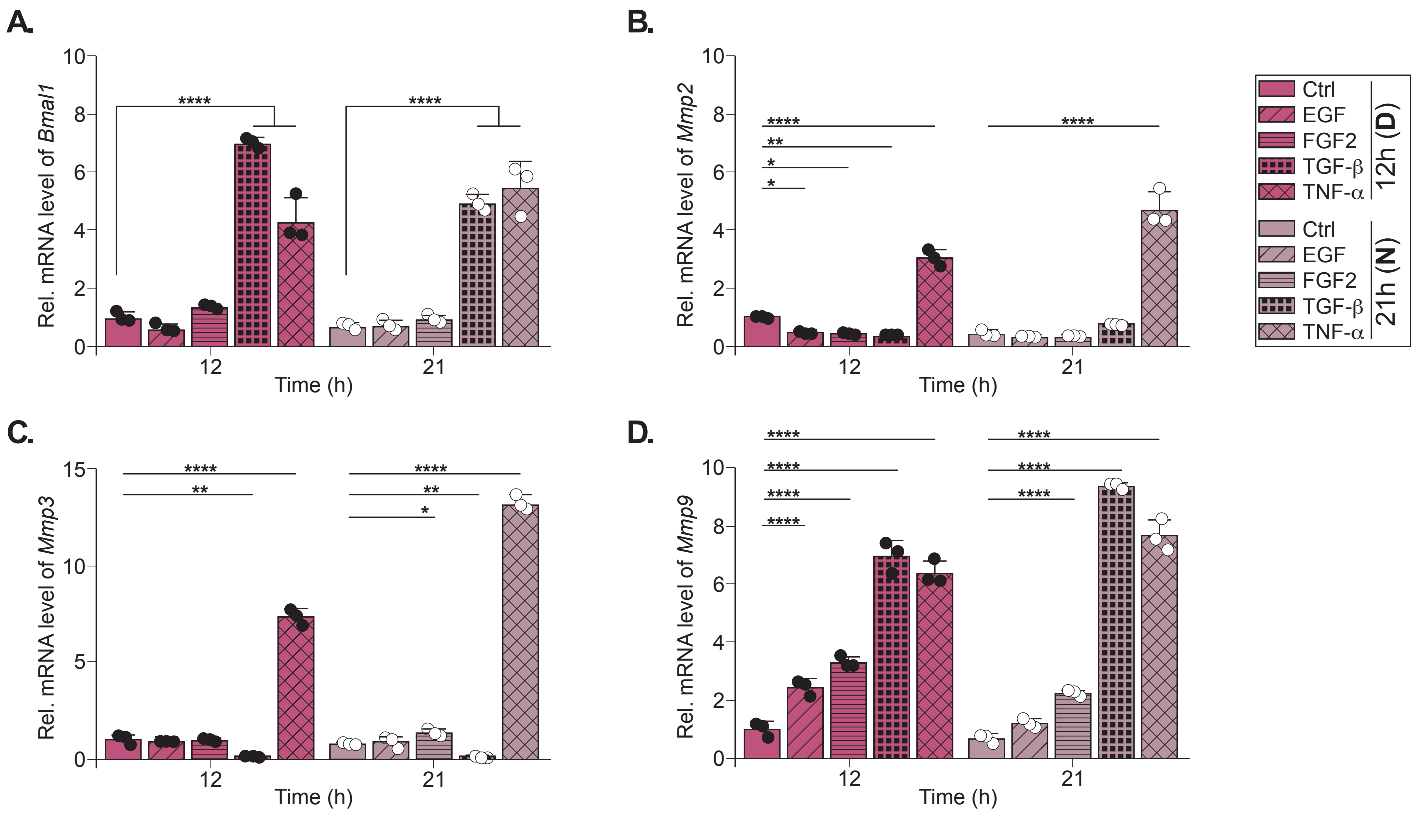
A.
Rel. mRNA level of Bmal1
10
8
6
4
2
0
****
****
12
21
Time (h)
B.
Rel. mRNA level of Mmp2
****
**
*
*
****
12
21
Time (h)
Ctrl
EGF
FGF2
TGF-β
TNF-α
12h (D)
Ctrl
EGF
FGF2
TGF-β
TNF-α
21h (N)
C.
Rel. mRNA level of Mmp3
15
10
5
0
****
**
****
**
*
12
21
Time (h)
D.
Rel. mRNA level of Mmp9
****
****
****
****
****
****
****
12
21
Time (h)

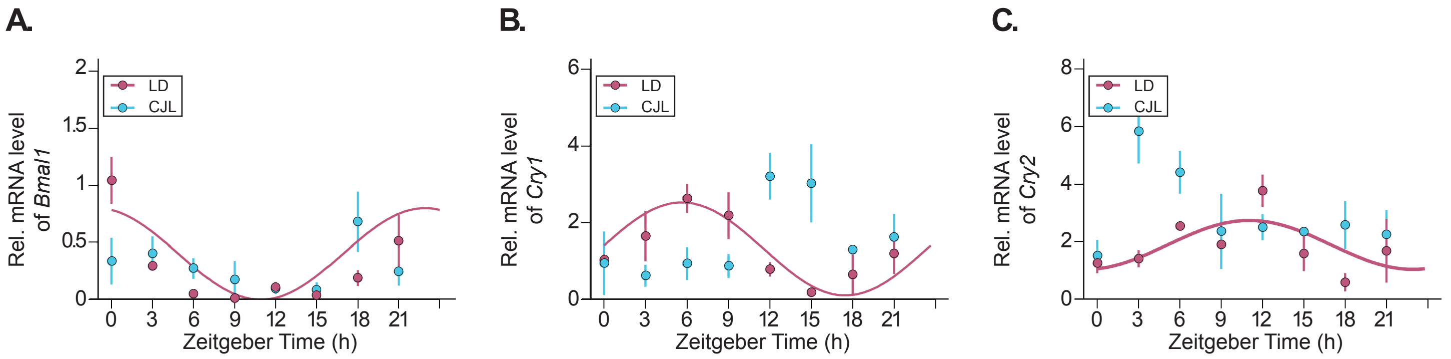
A.
B.
C.
LD
CJL
Rel. mRNA level of *Bmal1*
Rel. mRNA level of *Cry1*
Rel. mRNA level of *Cry2*
Zeitgeber Time (h)
0
3
6
9
12
15
18
21
0.5
1
1.5
2
4
6
8

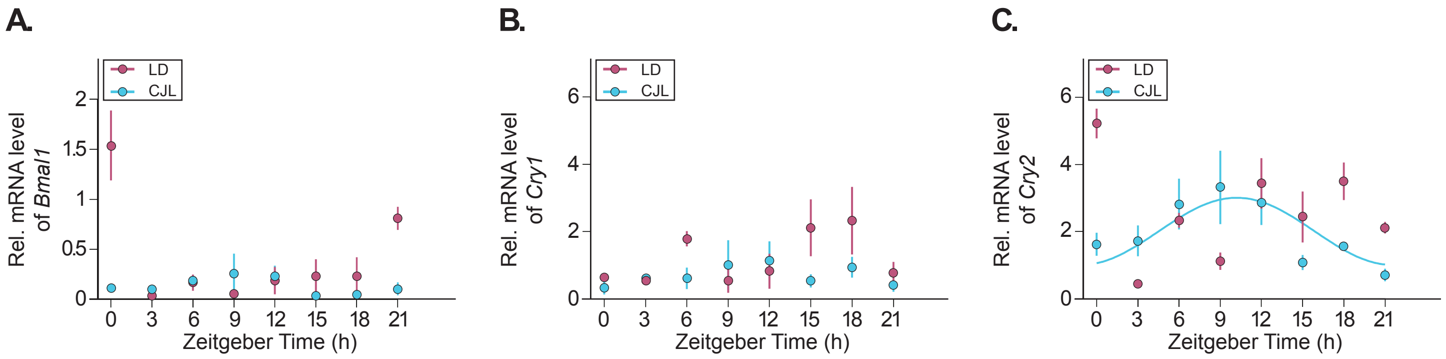
A.
LD
CJL
Rel. mRNA level of Bmal1
0
0.5
1
1.5
2
0 3 6 9 12 15 18 21
Zeitgeber Time (h)
B.
LD
CJL
Rel. mRNA level of Cry1
0
2
4
6
0 3 6 9 12 15 18 21
Zeitgeber Time (h)
C.
LD
CJL
Rel. mRNA level of Cry2
0
2
4
6
0 3 6 9 12 15 18 21
Zeitgeber Time (h)

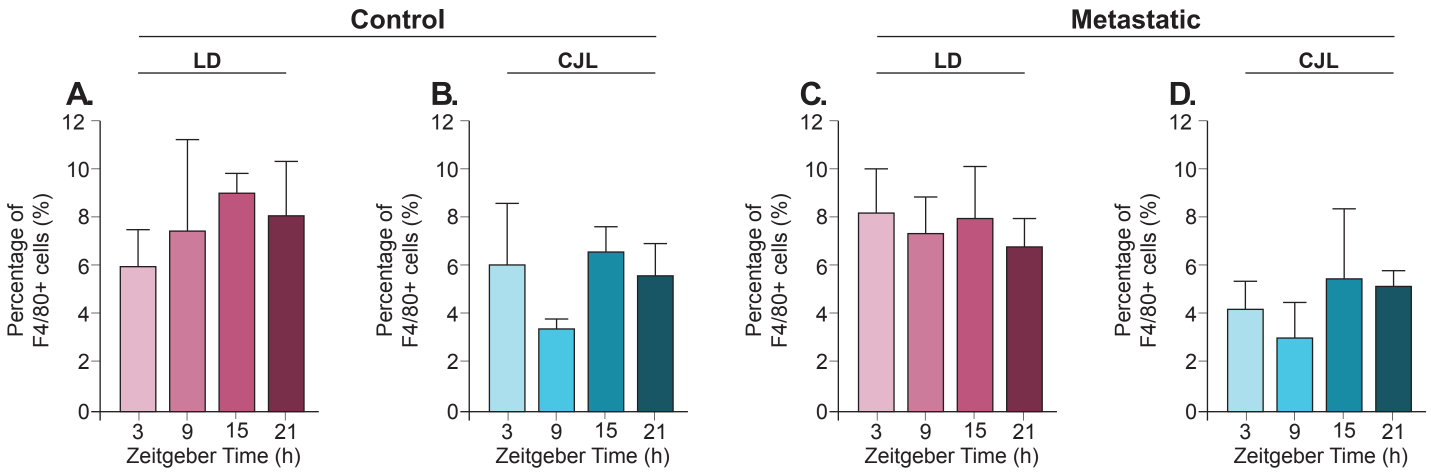
Control
Metastatic
LD
CJL
LD
CJL
A.
B.
C.
D.
Percentage of F4/80+ cells (%)
0
2
4
6
8
10
12
3
9
15
21
Zeitgeber Time (h)

**Supplementary Table 1. Statistical analysis of circadian clock inhibitor effects on MLg cell migration.**

| Post-hoc 2-way ANOVA | | | | | | | | |
|---|---|---|---|---|---|---|---|---|
| GSK2945 | | | KL001 | | | PF670 | | |
| comparison | p value | | comparison | p value | | comparison | p value | |
| Ctrl Unsync vs. Ctrl Sync | >0.9999 | ns | Ctrl Unsync vs. Ctrl Sync | 0.9996 | ns | Ctrl Unsync vs. Ctrl Sync | 0.9967 | ns |
| Ctrl Unsync vs. GSK2945 Unsync | <0.0001 | **** | Ctrl Unsync vs. KL001 Unsync | >0.9999 | ns | Ctrl Unsync vs. PF670 Unsync | >0.9999 | ns |
| Ctrl Unsync vs. GSK2945 Sync | <0.0001 | **** | Ctrl Unsync vs. KL001 Sync | >0.9999 | ns | Ctrl Unsync vs. PF670 Sync | >0.9999 | ns |
| Ctrl Unsync vs. TGF-β Unsync | <0.0001 | **** | Ctrl Unsync vs. KL001 + TGF-β Unsync | 0.0001 | *** | Ctrl Unsync vs. PF670 + TGF-β Unsync | 0.0007 | *** |
| Ctrl Unsync vs. TGF-β Sync | <0.0001 | **** | Ctrl Unsync vs. KL001 + TGF-β Sync | 0.0005 | *** | Ctrl Unsync vs. PF670 + TGF-β Sync | 0.0246 | * |
| Ctrl Unsync vs. TGF-β + GSK2945 Unsync | <0.0001 | **** | Ctrl Unsync vs. TGF-β Unsync | 0.0007 | *** | Ctrl Unsync vs. TGF-β Unsync | <0.0001 | **** |
| Ctrl Unsync vs. TGF-β + GSK2945 Sync | <0.0001 | **** | Ctrl Unsync vs. TGF-β Sync | 0.0003 | *** | Ctrl Unsync vs. TGF-β Sync | <0.0001 | **** |
| Ctrl Sync vs. GSK2945 Unsync | <0.0001 | **** | Ctrl Sync vs. KL001 Unsync | 0.9845 | ns | Ctrl Sync vs. PF670 Unsync | 0.9754 | ns |
| Ctrl Sync vs. GSK2945 Sync | <0.0001 | **** | Ctrl Sync vs. KL001 Sync | 0.9987 | ns | Ctrl Sync vs. PF670 Sync | 0.9983 | ns |
| Ctrl Sync vs. TGF-β Unsync | <0.0001 | **** | Ctrl Sync vs. KL001 + TGF-β Unsync | <0.0001 | **** | Ctrl Sync vs. PF670 + TGF-β Unsync | <0.0001 | **** |
| Ctrl Sync vs. TGF-β Sync | <0.0001 | **** | Ctrl Sync vs. KL001 + TGF-β Sync | 0.0001 | *** | Ctrl Sync vs. PF670 + TGF-β Sync | 0.0023 | ** |
| Ctrl Sync vs. TGF-β + GSK2945 Unsync | <0.0001 | **** | Ctrl Sync vs. TGF-β Unsync | 0.0004 | *** | Ctrl Sync vs. TGF-β Unsync | <0.0001 | **** |
| Ctrl Sync vs. TGF-β + GSK2945 Sync | <0.0001 | **** | Ctrl Sync vs. TGF-β Sync | 0.0001 | *** | Ctrl Sync vs. TGF-β Sync | <0.0001 | **** |
| GSK2945 Unsync vs. GSK2945 Sync | 0.0923 | ns | KL001 Unsync vs. KL001 Sync | >0.9999 | ns | PF670 Unsync vs. PF670 Sync | 0.9999 | ns |
| GSK2945 Unsync vs. TGF-β Unsync | <0.0001 | **** | KL001 Unsync vs. KL001 + TGF-β Unsync | 0.0001 | *** | PF670 Unsync vs. PF670 + TGF-β Unsync | <0.0001 | **** |
| GSK2945 Unsync vs. TGF-βSync | 0.7621 | ns | KL001 Unsync vs. KL001 + TGF-β Sync | 0.0006 | *** | PF670 Unsync vs. PF670 + TGF-β Sync | 0.0049 | ** |

| | | | | | | | | |
|---|---|---|---|---|---|---|---|---|
| GSK2945 Unsync vs. TGF-β + GSK2945 Unsync | 0.0668 | ns | KL001 Unsync vs. TGF-β Unsync | 0.0008 | *** | PF670 Unsync vs. TGF-β Unsync | <0.0001 | **** |
| GSK2945 Unsync vs. TGF-β + GSK2945 Sync | 0.9221 | ns | KL001 Unsync vs. TGF-β Sync | 0.0003 | *** | PF670 Unsync vs. TGF-β Sync | <0.0001 | **** |
| GSK2945 Sync vs. TGF-β Unsync | <0.0001 | **** | KL001 Sync vs. KL001 + TGF-β Unsync | 0.0001 | *** | PF670 Sync vs. PF670 + TGF-β Unsync | <0.0001 | **** |
| GSK2945 Sync vs. TGF-βSync | 0.0009 | *** | KL001 Sync vs. KL001 + TGF-β Sync | 0.0005 | *** | PF670 Sync vs. PF670 + TGF-β Sync | 0.0013 | ** |
| GSK2945 Sync vs. Tgf-β + GSK2945 Unsync | >0.9999 | ns | KL001 Sync vs. TGF-β Unsync | 0.0007 | *** | PF670 Sync vs. TGF-β Unsync | <0.0001 | **** |
| GSK2945 Sync vs. TGF-β + GSK2945 Sync | 0.7015 | ns | KL001 Sync vs. TGF-β Sync | 0.0003 | *** | PF670 Sync vs. TGF-β Sync | <0.0001 | **** |
| TGF-β Unsync vs. TGF-βSync | 0.016 | * | KL001 + TGF-β Unsync vs. KL001 + TGF-β Sync | 0.5151 | ns | PF670 + TGF-β Unsync vs. PF670 + TGF-β Sync | 0.8185 | ns |
| TGF-β Unsync vs. TGF-β + GSK2945 Unsync | <0.0001 | **** | KL001 + TGF-β Unsync vs. Tgf-β Unsync | >0.9999 | ns | PF670 + TGF-β Unsync vs. TGF-β Unsync | <0.0001 | **** |
| TGF-β Unsync vs. TGF-β + GSK2945 Sync | <0.0001 | **** | KL001 + TGF-β Unsync vs. Tgf-β Sync | >0.9999 | ns | PF670 + TGF-β Unsync vs. TGF-β Sync | <0.0001 | **** |
| TGF-βSync vs. TGF-β+ GSK2945 Unsync | 0.0006 | *** | KL001 + TGF-β Sync vs. Tgf-β Unsync | 0.744 | ns | PF670 + TGF-β Sync vs. TGF-β Unsync | <0.0001 | **** |
| TGF-βSync vs. TGF-β + GSK2945 Sync | 0.1163 | ns | KL001 + TGF-β Sync vs. Tgf-β Sync | 0.9899 | ns | PF670 + TGF-β Sync vs. TGF-β Sync | <0.0001 | **** |
| TGF-β+ GSK2945 Unsync vs. TGF + GSK2945 Sync | 0.6149 | ns | TGF-β Unsync vs. TGF-β Sync | >0.9999 | ns | TGF-β Unsync vs. TGF-β Sync | 0.0005 | *** |

| 2-way ANOVA | | | | | |
|---|---|---|---|---|---|
| **NIH PER2KO** | | | **MLg shBmal1** | | |
| **comparison** | **p value** | | **Comparison** | **p value** | |
| Ctrl Per2KO Unsync vs. Ctrl Per2KO Sync | 0.9825 | ns | shCtrl unsync vs. shCtrl sync | 0.6834 | ns |
| Ctrl Per2KO Unsync vs. Per2KO TGF unsync | 0.0606 | ns | shCtrl unsync vs. shBmal1 unsync | >0.9999 | ns |
| Ctrl Per2KO Unsync vs. Per2KO TGF sync | >0.9999 | ns | shCtrl unsync vs. shBmal1 sync | >0.9999 | ns |
| **Ctrl Per2KO Unsync vs. Control WT Unsync** | <0.0001 | **** | **shCtrl unsync vs. shCtrl TGF unsync** | **<0.0001** | **** |
| **Ctrl Per2KO Unsync vs. Control WT Sync** | <0.0001 | **** | shCtrl unsync vs. shCtrl TGF sync | 0.9998 | ns |
| **Ctrl Per2KO Unsync vs. WT TGF Unsync** | <0.0001 | **** | **shCtrl unsync vs. shBmal1 TGF unsync** | **<0.0001** | **** |
| **Ctrl Per2KO Unsync vs. WT TGF Sync** | <0.0001 | **** | **shCtrl unsync vs. shBmal1 TGF sync** | **<0.0001** | **** |
| **Ctrl Per2KO Sync vs. Per2KO TGF unsync** | 0.0044 | ** | shCtrl sync vs. shBmal1 unsync | 0.7435 | ns |
| Ctrl Per2KO Sync vs. Per2KO TGF sync | 0.9138 | ns | shCtrl sync vs. shBmal1 sync | 0.8593 | ns |
| **Ctrl Per2KO Sync vs. Control WT Unsync** | <0.0001 | **** | **shCtrl sync vs. shCtrl TGF unsync** | **<0.0001** | **** |
| **Ctrl Per2KO Sync vs. Control WT Sync** | <0.0001 | **** | shCtrl sync vs. shCtrl TGF sync | 0.0556 | ns |
| **Ctrl Per2KO Sync vs. WT TGF Unsync** | <0.0001 | **** | **shCtrl sync vs. shBmal1 TGF unsync** | **<0.0001** | **** |
| **Ctrl Per2KO Sync vs. WT TGF Sync** | <0.0001 | **** | **shCtrl sync vs. shBmal1 TGF sync** | **<0.0001** | **** |
| Per2KO TGF unsync vs. Per2KO TGF sync | 0.1297 | ns | shBmal1 unsync vs. shBmal1 sync | >0.9999 | ns |
| **Per2KO TGF unsync vs. Control WT Unsync** | <0.0001 | **** | **shBmal1 unsync vs. shCtrl TGF unsync** | **<0.0001** | **** |
| **Per2KO TGF unsync vs. Control WT Sync** | <0.0001 | **** | shBmal1 unsync vs. shCtrl TGF sync | 0.9994 | ns |

| | | | | | |
|---|---|---|---|---|---|
| **Per2KO TGF unsync vs. WT TGF Unsync** | <0.0001 | **** | **shBmal1 unsync vs. shBmal1 TGF unsync** | **<0.0001** | **** |
| **Per2KO TGF unsync vs. WT TGF Sync** | <0.0001 | **** | **shBmal1 unsync vs. shBmal1 TGF sync** | **<0.0001** | **** |
| **Per2KO TGF sync vs. Control WT Unsync** | <0.0001 | **** | **shBmal1 sync vs. shCtrl TGF unsync** | **<0.0001** | **** |
| **Per2KO TGF sync vs. Control WT Sync** | <0.0001 | **** | shBmal1 sync vs. shCtrl TGF sync | 0.9955 | ns |
| **Per2KO TGF sync vs. WT TGF Unsync** | <0.0001 | **** | **shBmal1 sync vs. shBmal1 TGF unsync** | **<0.0001** | **** |
| **Per2KO TGF sync vs. WT TGF Sync** | <0.0001 | **** | **shBmal1 sync vs. shBmal1 TGF sync** | **<0.0001** | **** |
| **Control WT Unsync vs. Control WT Sync** | 0.0095 | ** | **shCtrl TGF unsync vs. shCtrl TGF sync** | **<0.0001** | **** |
| **Control WT Unsync vs. WT TGF Unsync** | <0.0001 | **** | **shCtrl TGF unsync vs. shBmal1 TGF unsync** | **<0.0001** | **** |
| Control WT Unsync vs. WT TGF Sync | 0.1718 | ns | shCtrl TGF unsync vs. shBmal1 TGF sync | >0.9999 | ns |
| **Control WT Sync vs. WT TGF Unsync** | <0.0001 | **** | **shCtrl TGF sync vs. shBmal1 TGF unsync** | **<0.0001** | **** |
| **Control WT Sync vs. WT TGF Sync** | <0.0001 | **** | **shCtrl TGF sync vs. shBmal1 TGF sync** | **<0.0001** | **** |
| **WT TGF Unsync vs. WT TGF Sync** | <0.0001 | **** | **shBmal1 TGF unsync vs. shBmal1 TGF sync** | **<0.0001** | **** |

**Supplementary Table 2. Statistical analysis of genetic circadian disruption effects on cell migration.**

**Supplementary Table 3. MetaCycle analysis of clock and MMP gene rhythmicity in MLg cells.**

| Gene ID | Condition | Synchrony | p value | period | phase | mesor | amplitude |
|---|---|---|---|---|---|---|---|
| *Bmal1* | Control | **Sync** | **0.003** | **25.566** | **16.003** | **2.025** | **1.047** |
| | | Unsync | 0.255 | 25.031 | 17.734 | 2.095 | 0.625 |
| | TGF-β | **Sync** | **0.025** | **25.285** | **18.267** | **1.924** | **0.708** |
| | | Unsync | 0.576 | 24.169 | 14.606 | 5.819 | 2.077 |
| | GSK2945 | Sync | 0.067 | 26.333 | 11.007 | 1.185 | 0.291 |
| | | Unsync | 0.079 | 21.892 | 8.371 | 1.548 | 0.471 |
| *Mmp3* | Control | **Sync** | **0.00001** | **25.077** | **15.280** | **2.024** | **0.803** |
| | | Unsync | 0.260 | 25.362 | 15.937 | 4.887 | 3.040 |
| | TGF-β | **Sync** | **0.021** | **27.574** | **0.412** | **0.284** | **0.217** |
| | | **Unsync** | **0.003** | **23.197** | **1.286** | **0.841** | **0.813** |
| | GSK2945 | Sync | 0.107 | 26.333 | 26.034 | 0.500 | 0.239 |
| | | **Unsync** | **0.002** | **26.333** | **23.165** | **1.320** | **0.677** |
| *Mmp2* | Control | **Sync** | **0.001** | **20.899** | **12.741** | **1.094** | **0.382** |
| | | **Unsync** | **0.046** | **25.551** | **19.503** | **1.047** | **0.264** |
| | TGF-β | **Sync** | **0.001** | **26.180** | **14.459** | **3.428** | **1.818** |
| | | Unsync | 0.823 | 26.581 | 17.075 | 6.646 | 0.484 |
| | GSK2945 | **Sync** | **0.021** | **25.565** | **3.219** | **0.662** | **0.470** |
| | | Unsync | 0.163 | 25.153 | 9.663 | 1.069 | 0.088 |
| *Mmp9* | Control | **Sync** | **0.037** | **24.672** | **13.480** | **0.986** | **0.328** |
| | | Unsync | 0.149 | 26.333 | 19.009 | 2.978 | 1.131 |
| | TGF-β | **Sync** | **0.0001** | **26.242** | **12.693** | **2.589** | **1.535** |
| | | Unsync | 0.157 | 26.791 | 16.712 | 8.626 | 1.999 |
| | GSK2945 | Sync | 0.455 | 24.169 | 3.486 | 0.534 | 0.145 |
| | | **Unsync** | **0.009** | **26.447** | **23.997** | **1.590** | **0.647** |

**Supplementary Table 4. MetaCycle analysis of core clock gene oscillations in lung tissue.**

| Gene ID | Condition | Light schedule | P value | period | phase | Mesor | Amplitude |
|---|---|---|---|---|---|---|---|
| ***Bmal1*** | Control lung | **LD** | **0.041** | **24.098** | **22.408** | **0.270** | **0.369** |
| | | CJL | 0.822 | 21.645 | 2.447 | 0.245 | 0.121 |
| | Metastatic lung | LD | 0.668 | 25.333 | 21.792 | 0.275 | 0.235 |
| | | CJL | 0.413 | 21.597 | 9.666 | 0.165 | 0.103 |
| ***Cry2*** | Control lung | **LD** | **0.047** | **22.563** | **10.984** | **1.640** | **0.903** |
| | | CJL | 0.957 | 22.666 | 4.156 | 2.236 | 1.044 |
| | Metastatic lung | LD | 0.893 | 21.666 | 18.33 | 2.079 | 1.218 |
| | | **CJL** | **0.012** | **22.085** | **8.88** | **2.207** | **1.465** |
| ***Cry1*** | Control lung | **LD** | **0.006** | **22.654** | **7.336** | **1.123** | **0.843** |
| | | CJL | 0.858 | 21.666 | 13.632 | 1.352 | 0.650 |
| | Metastatic lung | LD | 0.983 | 25.333 | 10.946 | 0.949 | 0.184 |
| | | CJL | 0.431 | 24.875 | 9.806 | 0.690 | 0.273 |

**Supplementary Table 5. MetaCycle analysis of pathway component rhythmicity in control and metastatic lungs.**

| Gene ID | Condition | Light schedule | P value | period | phase | Mesor | Amplitude |
|---|---|---|---|---|---|---|---|
| ***Tnf-α*** | Control lung | LD | 0.530 | 21.666 | 19.901 | 0.385 | 0.311 |
| | | CJL | 0.280 | 24.767 | 18.452 | 1.289 | 1.033 |
| | Metastatic lung | LD | 0.594 | 23.588 | 0.349 | 0.374 | 0.231 |
| | | CJL | 0.997 | 23.671 | 23.043 | 0.162 | 0.024 |
| ***Tgf-β*** | Control lung | LD | 0.949 | 25.425 | 18.613 | 0.663 | 0.310 |
| | | CJL | 1 | 22.197 | 7.831 | 0.650 | 0.012 |
| | Metastatic lung | LD | 0.980 | 23.875 | 19.450 | 0.480 | 0.147 |
| | | CJL | 0.814 | 22.856 | 15.528 | 1.047 | 0.720 |
| ***Ccl2*** | Control lung | LD | 0.800 | 21.179 | 0.506 | 0.739 | 0.255 |
| | | **CJL** | **0.039** | **20.502** | **10.538** | **0.815** | **0.607** |
| | Metastatic lung | LD | 0.782 | 21.666 | 16.278 | 0.840 | 0.727 |
| | | CJL | 0.253 | 21.666 | 5.358 | 0.957 | 0.811 |
| ***Mmp3*** | Control lung | **LD** | **0.004** | **24.460** | **22.201** | **0.732** | **0.551** |
| | | CJL | 0.546 | 24.767 | 19.725 | 1.258 | 0.600 |
| | Metastatic lung | LD | 0.999 | 25.333 | 0.013 | 1.030 | 0.004 |
| | | CJL | 0.979 | 21.728 | 19.527 | 0.607 | 0.177 |
| ***Mmp9*** | Control lung | LD | 0.236 | 0.236 | 0.236 | 0.236 | 1.142 |
| | | CJL | 0.997 | 0.997 | 0.997 | 0.997 | 0.219 |
| | Metastatic lung | LD | 0.787 | 0.787 | 0.787 | 0.787 | 0.261 |
| | | CJL | 0.953 | 0.953 | 0.953 | 0.953 | 0.527 |
| ***Mmp2*** | Control lung | LD | 0.999 | 0.999 | 15.150 | 0.666 | 0.187 |
| | | CJL | 0.131 | 0.131 | 20.128 | 1.352 | 0.642 |
| | Metastatic lung | **LD** | **0.027** | **24.027** | **7.690** | **0.484** | **0.257** |
| | | CJL | 0.111 | 0.111 | 19.123 | 0.483 | 0.166 |
| ***Mob1a*** | Control lung | LD | 0.590 | 20.777 | 8.526 | 0.872 | 0.577 |
| | | CJL | 0.811 | 21.666 | 14.840 | 1.252 | 1.137 |
| | Metastatic lung | LD | 0.233 | 24.122 | 2.475 | 0.669 | 0.227 |
| | | CJL | 0.874 | 24.725 | 13.526 | 0.626 | 0.245 |
| ***Tead4*** | Control lung | LD | 0.996 | 24.333 | 18.126 | 0.552 | 0.261 |
| | | CJL | 0.955 | 21.666 | 14.526 | 1.162 | 0.534 |
| | Metastatic lung | LD | 0.595 | 24.875 | 0.852 | 1.537 | 0.474 |
| | | CJL | 0.875 | 21.666 | 16.262 | 1.258 | 0.293 |
| ***Smad4*** | Control lung | LD | 0.553 | 25.098 | 0.814 | 0.634 | 0.487 |
| | | CJL | 0.248 | 22.060 | 10.470 | 0.778 | 0.425 |
| | Metastatic lung | LD | 0.251 | 22.819 | 17.399 | 0.970 | 0.523 |
| | | CJL | 0.128 | 22.840 | 13.772 | 0.913 | 0.633 |
| ***Smad7*** | Control lung | LD | 0.998 | 25.574 | 18.789 | 0.459 | 0.250 |
| | | CJL | 0.740 | 21.666 | 1.506 | 1.403 | 1.176 |
| | Metastatic lung | LD | 0.951 | 25.632 | 3.273 | 0.586 | 0.514 |

| | | | | | | | |
|---|---|---|---|---|---|---|---|
| | | CJL | 0.185 | 24.560 | 10.684 | 0.427 | 0.300 |
| *Snail* | Control lung | LD | 0.463 | 21.666 | 2.915 | 0.916 | 0.527 |
| | | CJL | 0.994 | 21.666 | 6.626 | 1.379 | 0.498 |
| | Metastatic lung | LD | 0.999 | 25.333 | 24.104 | 0.627 | 0.0171 |
| | | CJL | 0.965 | 21.523 | 15.516 | 0.642 | 0.262 |
| *Zeb1* | Control lung | LD | 0.604 | 22.343 | 6.083 | 1.071 | 1.122 |
| | | CJL | 0.095 | 22.041 | 14.093 | 1.353 | 0.954 |
| | Metastatic lung | LD | 0.997 | 22.006 | 3.951 | 1.278 | 0.690 |
| | | CJL | 0.999 | 24.333 | 15.192 | 0.956 | 0.251 |
| *Fibronectin* | Control lung | LD | 0.178 | 22.269 | 6.203 | 1.178 | 1.066 |
| | | CJL | 0.878 | 25.333 | 16.799 | 0.999 | 0.206 |
| | Metastatic lung | LD | 0.944 | 21.580 | 7.721 | 1.551 | 1.078 |
| | | CJL | 0.999 | 21.059 | 5.387 | 0.960 | 0.275 |

| Post-hoc 2-way ANOVA | | | | | | | | |
|---|---|---|---|---|---|---|---|---|
| **Verteporfin at 0h** | | | **Verteporfin at 6h** | | | **Verteporfin at 12h** | | |
| **comparison** | **p value** | | **comparison** | **p value** | | **comparison** | **p value** | |
| Ctrl Unsync vs. Ctrl Sync | >0.9999 | ns | Ctrl Unsync vs. Ctrl Sync | >0.9999 | ns | Ctrl -Dex vs. Ctrl +Dex | >0.9999 | ns |
| Ctrl Unsync vs. Tgf-β + verte0h Unsync | 0.9992 | ns | Ctrl Unsync vs. Tgf-β + verte6h Unsync | 0.9983 | ns | **Ctrl -Dex vs. Tgf-β + verte12h Unsync** | **<0.0001** | ******** |
| Ctrl Unsync vs. Tgf-β + verte0h Sync | 0.9994 | ns | Ctrl Unsync vs. Tgf-β + verte6h Sync | >0.9999 | ns | **Ctrl -Dex vs. Tgf-β + verte12h sync** | **0.0006** | ******* |
| **Ctrl Unsync vs. TGF-β Unsync** | **<0.0001** | ******** | **Ctrl Unsync vs. TGF-β Unsync** | **<0.0001** | ******** | **Ctrl -Dex vs. TGF-βUnsync** | **<0.0001** | ******** |
| **Ctrl Unsync vs. TGF-β Sync** | **<0.0001** | ******** | **Ctrl Unsync vs. TGF-β Sync** | **0.0004** | ******* | **Ctrl -Dex vs. TGF-βSync** | **<0.0001** | ******** |
| Ctrl Unsync vs. Verteporfin Unsync | 0.8053 | ns | Ctrl Unsync vs. Verteporfin Unsync | 0.9392 | ns | Ctrl -Dex vs. Verteporfin Unsync | 0.5655 | ns |
| Ctrl Unsync vs. Verteporfin Sync | >0.9999 | ns | Ctrl Unsync vs. Verteporfin Sync | >0.9999 | ns | Ctrl -Dex vs. Verteporfin Sync | >0.9999 | ns |
| Ctrl Sync vs. Tgf-β + verte0h Unsync | >0.9999 | ns | Ctrl Sync vs. Tgf-β + verte6h Unsync | 0.9999 | ns | **Ctrl +Dex vs. Tgf-β + verte12h Unsync** | **<0.0001** | ******** |
| Ctrl Sync vs. Tgf-β + verte0h Sync | 0.9934 | ns | Ctrl Sync vs. Tgf-β + verte6h Sync | >0.9999 | ns | **Ctrl +Dex vs. Tgf-β + verte12h sync** | **0.0003** | ******* |
| **Ctrl Sync vs. TGF-β Unsync** | **<0.0001** | ******** | **Ctrl Sync vs. TGF-β Unsync** | **<0.0001** | ******** | **Ctrl +Dex vs. TGF-βUnsync** | **<0.0001** | ******** |
| **Ctrl Sync vs. TGF-β Sync** | **<0.0001** | ******** | **Ctrl Sync vs. TGF-β Sync** | **0.0007** | ******* | **Ctrl +Dex vs. TGF-βSync** | **<0.0001** | ******** |
| Ctrl Sync vs. Verteporfin Unsync | 0.9197 | ns | Ctrl Sync vs. Verteporfin Unsync | 0.9819 | ns | Ctrl +Dex vs. Verteporfin Unsync | 0.7562 | ns |
| Ctrl Sync vs. Verteporfin Sync | >0.9999 | ns | Ctrl Sync vs. Verteporfin Sync | >0.9999 | ns | Ctrl +Dex vs. Verteporfin Sync | >0.9999 | ns |
| Tgf-β + verte0h Unsync vs. Tgf-β + verte0h Sync | 0.4476 | ns | Tgf-β + verte6h Unsync vs. Tgf-β + verte6h Sync | >0.9999 | ns | **Tgf-β + verte12h Unsync vs. Tgf-β + verte12h sync** | **<0.0001** | ******** |
| **Tgf-β + verte0h Unsync vs. TGF-β Unsync** | **<0.0001** | ******** | **Tgf-β + verte6h Unsync vs. TGF-β Unsync** | **<0.0001** | ******** | **Tgf-β + verte12h Unsync vs. TGF-βUnsync** | **<0.0001** | ******** |

| | | | | | | | | |
|---|---|---|---|---|---|---|---|---|
| Tgf-β + verte0h Unsync vs. TGF-β Sync | 0.0012 | ** | Tgf-β + verte6h Unsync vs. TGF-β Sync | 0.0115 | * | Tgf-β + verte12h Unsync vs. TGF-βSync | 0.6 | ns |
| Tgf-β + verte0h Unsync vs. Verteporfin Unsync | >0.9999 | ns | Tgf-β + verte6h Unsync vs. Verteporfin Unsync | >0.9999 | ns | Tgf-β + verte12h Unsync vs. Verteporfin Unsync | <0.0001 | **** |
| Tgf-β + verte0h Unsync vs. Verteporfin Sync | >0.9999 | ns | Tgf-β + verte6h Unsync vs. Verteporfin Sync | >0.9999 | ns | Tgf-β + verte12h Unsync vs. Verteporfin Sync | <0.0001 | **** |
| Tgf-β + verte0h Sync vs. TGF-β Unsync | <0.0001 | **** | Tgf-β + verte6h Sync vs. TGF-β Unsync | <0.0001 | **** | Tgf-β + verte12h sync vs. TGF-β Unsync | <0.0001 | **** |
| Tgf-β + verte0h Sync vs. TGF-β Sync | <0.0001 | **** | Tgf-β + verte6h Sync vs. TGF-β Sync | 0.0007 | *** | Tgf-β + verte12h sync vs. TGF-βSync | <0.0001 | **** |
| Tgf-β + verte0h Sync vs. Verteporfin Unsync | 0.0963 | ns | Tgf-β + verte6h Sync vs. Verteporfin Unsync | 0.9869 | ns | Tgf-β + verte12h sync vs. Verteporfin Unsync | <0.0001 | **** |
| Tgf-β + verte0h Sync vs. Verteporfin Sync | 0.825 | ns | Tgf-β + verte6h Sync vs. Verteporfin Sync | >0.9999 | ns | Tgf-β + verte12h sync vs. Verteporfin Sync | <0.0001 | **** |
| TGF-β Unsync vs. TGF-β Sync | <0.0001 | **** | TGF-β Unsync vs. TGF-β Sync | <0.0001 | **** | TGF-β Unsync vs. TGF-βSync | <0.0001 | **** |
| TGF-β Unsync vs. Verteporfin Unsync | <0.0001 | **** | TGF-β Unsync vs. Verteporfin Unsync | <0.0001 | **** | TGF-β Unsync vs. Verteporfin Unsync | <0.0001 | **** |
| TGF-β Unsync vs. Verteporfin Sync | <0.0001 | **** | TGF-β Unsync vs. Verteporfin Sync | <0.0001 | **** | TGF-β Unsync vs. Verteporfin Sync | <0.0001 | **** |
| TGF-β Sync vs. Verteporfin Unsync | 0.0084 | ** | TGF-β Sync vs. Verteporfin Unsync | 0.0337 | * | TGF-β Sync vs. Verteporfin Unsync | 0.0015 | ** |
| TGF-β Sync vs. Verteporfin Sync | 0.0003 | *** | TGF-β Sync vs. Verteporfin Sync | 0.0022 | ** | TGF-β Sync vs. Verteporfin Sync | <0.0001 | **** |
| Verteporfin Unsync vs. Verteporfin Sync | 0.9991 | ns | Verteporfin Unsync vs. Verteporfin Sync | >0.9999 | ns | Verteporfin Unsync vs. Verteporfin Sync | 0.9922 | ns |

**Supplementary Table 6. Statistical analysis of YAP inhibitor temporal effects on cell migration.**

**Supplementary Table 7. Primer sequences for quantitative real-time PCR analysis.**

| Gene | Primer | Sequence (5' to 3') | Product (bp) |
|---|---|---|---|
| ***Bmal1*** | Forward | CCAACCTTCCCGCAGCTAAC | 155 |
| | Reverse | TCCTCCGCGATCATTCGACC | |
| ***Cry1*** | Forward | ATCATTGGCGTGGACTAC | 79 |
| | Reverse | TCTGCTTCATTCGTTCA | |
| ***Cry2*** | Forward | GGTTGCCTGTTTCCTGACTC | 195 |
| | Reverse | TTGGGATCTGTCCTCCTACC | |
| ***Mmp3*** | Forward | GCTCATGAACTTGGCCACTC | 156 |
| | Reverse | TGCTGTGGGAGTTCCATAGAGG | |
| ***Mmp9*** | Forward | CTTCCAGTACCAAGACAAAGCC | 249 |
| | Reverse | CAGCTAGCACCTTTCCCTCG | |
| ***Mmp2*** | Forward | CAGGAGACAAGTTCTGGAGATACA | 227 |
| | Reverse | GCCCAGCCAGTCTGATTTGA | |
| ***Tnf-α*** | Forward | GACAGTGACCTGGACTGTGG | 132 |
| | Reverse | GAGACAGAGGCAACCTGACC | |
| ***Tgf-β*** | Forward | ATTTGGCTTGAGATGGTTGG | 110 |
| | Reverse | CTTCGGGTGAGACCACAAAT | |
| ***Ccl2*** | Forward | CGGCTGGAGCATCCACGTGTT | 191 |
| | Reverse | TGGGGTCAGCACAGACCTCTC | |
| ***Mob1a*** | Forward | GGCATCATTAGAAGAATCAGCAG | 146 |
| | Reverse | GGCAACATAACAGCTTGCCT | |
| ***Tead4*** | Forward | GCCCATCGACAATGATGCAG | 270 |
| | Reverse | CCGTTCCTCGAAACGGTCC | |
| ***Smad4*** | Forward | ATAGCTCCAGCCATCAGTCTGTC | 254 |
| | Reverse | AAATGGTTAGGGCGTCCGTG | |
| ***Smad7*** | Forward | TTCTCAAACCAACTGCAGGCT | 185 |
| | Reverse | GACACAGTAGAGCCTCCC | |
| ***Zeb1*** | Forward | AGCCAAACGGAAACCAGGAT | 182 |
| | Reverse | GTGTCTCAACAGTGAGCTGC | |
| ***Snail*** | Forward | GTCCAGCTGTAACCATGCCT | 106 |
| | Reverse | TGTCACCAGGACAAATGGGG | |
| ***Fibronectin*** | Forward | CACCAACCCTGGGTATGACA | 168 |
| | Reverse | ACATTCGGCAGGTATGGTCT | |
| ***Hprt*** | Forward | TGTTGGATACAGGCCAGAC | 196 |
| | Reverse | TGGCAACATCAACAGGACTC | |
